# Quantitatively mapping local quality of super-resolution microscopy by rolling Fourier ring correlation

**DOI:** 10.1101/2022.12.01.518675

**Authors:** Weisong Zhao, Xiaoshuai Huang, Jianyu Yang, Guohua Qiu, Liying Qu, Yue Zhao, Shiqun Zhao, Ziying Luo, Xinwei Wang, Yaming Jiu, Heng Mao, Xumin Ding, Jiubin Tan, Ying Hu, Leiting Pan, Liangyi Chen, Haoyu Li

**Affiliations:** Innovation Photonics and Imaging Center, School of Instrumentation Science and Engineering, Harbin Institute of Technology, Harbin 150080, China; Biomedical Engineering Department, Peking University, Beijing 100191, China; The Key Laboratory of Weak-Light Nonlinear Photonics of Education Ministry, School of Physics and TEDA Institute of Applied Physics, Frontiers Science Center for Cell Responses, Nankai University, Tianjin 300071, China; State Key Laboratory of Membrane Biology, Beijing Key Laboratory of Cardiometabolic Molecular Medicine, Institute of Molecular Medicine, National Biomedical Imaging Center, Center for Life Sciences, School of Future Technology, Peking University, Beijing 100871, China; Department of Control Science and Engineering, Harbin Institute of Technology, Harbin 150001, China; Unit of Cell Biology and Imaging Study of Pathogen Host Interaction, The Center for Microbes, Development and Health, Key Laboratory of Molecular Virology and Immunology, Institut Pasteur of Shanghai, Chinese Academy of Sciences, Shanghai 200031, China; School of Mathematical Sciences, Peking University, Beijing 100871, China; Key Laboratory of Ultra-Precision Intelligent Instrumentation of Ministry of Industry and Information Technology, Harbin Institute of Technology, Harbin 150080, China; School of Life Science and Technology, Harbin Institute of Technology, Harbin 150001, China; International Cancer Institute, Peking University, Beijing 100191, China; Beijing Academy of Artificial Intelligence, Beijing 100871, China; PKU-IDG/McGovern Institute for Brain Research, Beijing 100871, China

## Abstract

In fluorescence microscopy, computational algorithms have been developed to suppress noise, enhance contrast, and even enable super-resolution (SR). However, the local quality of the images may vary on multiple scales, and these differences can lead to misconceptions, which is especially intractable in emerging deep-learning ones. Current mapping methods fail to finely estimate the local quality, challenging to associate the SR scale content. Here, we develop a rolling Fourier ring correlation (rFRC) framework to evaluate the reconstruction uncertainties down to SR scale. To visually pinpoint regions with low reliability, a filtered rFRC is combined with a modified resolution scaled error map (RSM), offering a comprehensive and concise map for further examination. We demonstrate their performances on various SR imaging modalities, and the resulting quantitative maps enable better SR images integrated from different reconstructions. Beyond that, we provide a strategy for learning-based restorations, allowing a direct detection of both data and model uncertainties, and expect the representative cases can inspire further advances in this rapidly developing field.

---

By implementing fluorescent probes and combining specific excitation and emission protocols, super-resolution (SR) fluorescence microscopy breaks the diffraction limit of resolution^1^, in which many methods heavily depend on image calculation and processing that retrieve SR information^1, 2^. Intrinsically, the noise and distortions in raw images caused by the photophysics of fluorophores^3–5^, the chemical environment of the sample^3, 4, 6^, and the optical setup conditions^4, 7–10^, may influence the qualities of the final SR images^11–13^. Because these factors are related to specific experimental configurations, a reliable and reference-free estimation of the image quality is invaluable to subsequent analysis, especially at the SR scale.

To evaluate the global effective resolution in situ, the Fourier ring correlation (FRC)^14^ describes the highest reliable cut-off frequency of an image. This effective resolution, or equivalently the spectral signal-to-noise ratio (SNR), is one crucial SR image quality metric, reflecting the authentic resolvability or the uncertainty^15^. However, the local resolution may vary dramatically over the imaging field. For example, in single-molecule localization microscopy (SMLM)^16–18^, the practical resolutions at different local regions are generally determined by the corresponding molecule active intensity and density, as well as the local background level^11^. To measure this resolution heterogeneity, the block-wise FRC calculation^14, 19^ was introduced, but it is still too coarse to describe the SR scale spatial separation of the resolution variation. The upscaled resolvability of SR imaging requires a more elaborate evaluation.

Here, we propose a rolling Fourier ring correlation (rFRC) method to draw the resolution heterogeneity directly in the SR domain, which allows for a mapping at an unprecedented SR scale and seamlessly correlates the resolution map with the SR content. Moreover, the variations of different SR reconstruction methods are usually on a fine scale, and our rFRC provides a prerequisite for assessing these methods objectively. Thus, it enables advancing process procedures to improve image restoration quality, such as fusing SMLM images reconstructed by different algorithms to yield SR images with better quality. Although we are limited to calculate the errors as without ground-truth comparing, we can measure the uncertainties by this rFRC to uncover the errors contained in the corresponding SR images. In other words, the lower spectral SNR (effective resolution) gives a higher probability of the error existence^15^, and thus we can use it to represent the uncertainty revealing the error distribution.

As a model-independent assessment, the rFRC using two independent captures may fail to identify regions that were always incorrectly restored during different reconstructions, possibly due to systematic image processing bias (model bias). On the other hand, the resolution-scaled error map (RSM)^19^ can evaluate reconstruction errors against the simultaneously acquired high SNR wide-field reference, assuming a spatially invariant Gaussian kernel and homogenous illumination. However, RSM suffers from false-negative identifications when the assumptions fail, and its detectable error scale is limited by the diffraction barrier. In this sense, RSM can only estimate the large-scale errors, such as the complete absence and distortion of structures, possibly induced by model bias, which can be a complementary module. We also accompany our rFRC with a truncated RSM, namely PANEL (pixel-level analysis of error locations), pinpointing the regions with low reliability for subsequent biological profiling.

We applied our quantitative maps in many SR approaches, including SMLM, SR radial fluctuations (SRRF)^20^, structured illumination microscopy (SIM)^21^, and deconvolution^22, 23^ (**Supplementary Note 2-5**), demonstrating its effectiveness. Beyond that, we explored that our rFRC can also be applied to evaluate the local restoration qualities of the deep-learning methods. At present, the importance of reconstruction uncertainty is attracting more attentions, as the out-of-distribution tests leading to hallucinations far from the truth. Based on rFRC mapping, we offer an effective solution for both data uncertainty and model uncertainty estimation. To study the special mechanism of the model and data uncertainties, we designed a simulated experiment with intentionally induced model uncertainty, and we found a large increase in detected data uncertainty, indicating the leakage of model uncertainty. This phenomenon suggests that the data uncertainty may be even more crucial for deep-learning microscopies. Finally, for expecting our method can be a routinely used local quality evaluation tool, it has been implemented as an open-source framework; the related source codes and the out-of-the-box Fiji/ImageJ^24^ plugin are available on GitHub (**Methods**).

## RESULTS

### The rFRC mapping and PANEL pinpointing

To provide local quality measurements directly at the SR scale, we transformed the conventional FRC into a rolling FRC (rFRC) map (**Fig. 1a**, **Methods**). The rFRC calculation is similar to that of a moving filter on an image. We assigned the corresponding FRC resolution in each block by sliding a window through the image. (i) To calculate the FRC of pixels at the image boundaries, we padded the input image symmetrically around a half size of the block (Step 1, **Fig. 1a**). (ii) To eliminate background-induced false negatives, we avoided calculating areas indistinguishable from the background (**Methods**, **Supplementary Fig. 2**), in which we assigned calculated the FRC resolution to the center pixel of each block only if its mean intensity was larger than a given threshold (Steps 2-4, **Fig. 1a**). To avoid overconfident and unstable determinations from small image blocks, in this work we used the 3σ curve^25^ as criterion (**Methods**, **Supplementary Note 1.1**). (iii) Afterward, the same procedure was repeated block by block throughout the entire image. Using the above rFRC as the metric we can quantitatively map the uncertainties in the SR reconstructions at their SR scale (**Supplementary Note 1.2**). We also offer two colormaps that may be more suitable for human intuition^26^ to display the uncertainties (shifted Jet, sJet; and black Jet, bJet) (Step 5, **Fig. 1a**, **Methods**, **Supplementary Fig. 3a**). In addition to local quality assessment, we calculate two global metrics, the rFRC value, a dimensionless metric with values starting at 0 reflecting the deterioration rate across the imaging field, and the rFRC resolution, representing the averaged resolution (**Methods**).

**Fig. 1.**
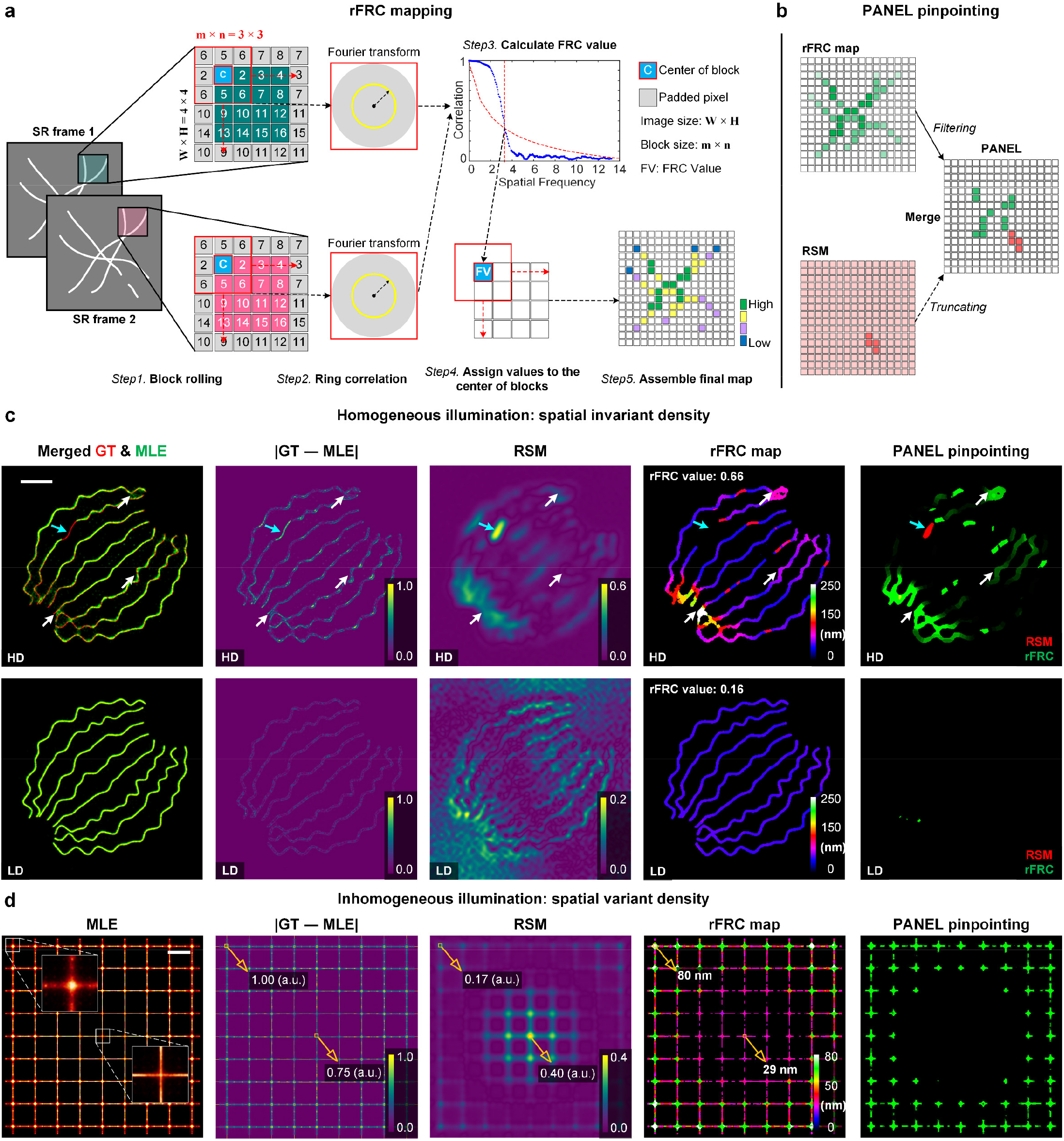
Overview of rFRC mapping and PANEL pinpointing. (**a**, **b**) Workflows. (**a**) The workflow of the rFRC map. **Step 1**, the symmetrically padded (gray pixels) two input images are clipped to small subsets for FRC calculation. The center pixel with an intensity lower than the background threshold will be skipped; otherwise, the following Steps 2-5 will be executed. **Steps 2-3**, FRC calculation, ring correlation in Fourier domain (Step 2), and FRC resolution determination (Step 3). **Step 4**, assign the obtained FRC resolutions to the corresponding center pixels. **Step 5**, assemble the final rFRC map and render it with the corresponding color map. (**b**) PANEL pinpointing. To highlight regions with low reliability, the rFRC map with values under the Otsu-determined threshold; and the normalized RSM with values under 0.5 will be filtered. Its abstract version can be seen in **Supplementary Fig. 1**. (**c**, **d**) Validations. (**c**) Simulations of 2D-SMLM with homogeneous illumination (inducing overall spatial invariant active density), with high-density (‘HD’, top) and low-density (‘LD’, bottom) emitting fluorophores in each frame. From left to right: Merged MLE reconstructions (green channel) and ground-truth images (red channel); Spatial subtractions between ground-truth images and MLE reconstructions; Spatial subtractions between wide-field ground-truth images and wide-field images generated from the MLE reconstructions, a.k.a. the RSM; The rFRC maps of two MLE reconstructions from odd frames (MLE_odd_) and even frames (MLEeven), respectively; The full PANEL visualizations (RSM corresponding to the red channel and rFRC map to the green channel). Cyan and white arrows represent the errors found by the RSM or rFRC map, respectively. (**d**) 2D-SMLM simulation with inhomogeneous illumination (high intensity in the center and decreasing illumination toward the edges). From left to right: The MLE result; Spatial subtraction between ground-truth image and MLE result; The RSM; The rFRC map. Scale bars: (**c**) 500 nm; (**d**) 1 μm.

In addition, we realized that the rFRC may not identify the regions that were always incorrectly restored during different reconstructions due to the model bias. For example, if the two reconstructed images lost an identical component, the rFRC may indicate a false positive in the corresponding region. To moderate this issue, we combined a modified RSM (**Methods**, **Supplementary Fig. 4**) with our rFRC to constitute the PANEL (**Fig. 1b**, **Methods**), for pinpointing such regions with low reliability. As small intensity fluctuations can lead to potential false negatives, we truncated the RSM with a hard threshold (0.5, **Methods**), only including prominent artifacts such as misrepresentations or the disappearance of structures. To filter the regions with high quality (high FRC resolution), we adopted the Otsu-based^27^ segmentation to highlight regions giving a higher probability of the error existence (**Methods**, **Supplementary Fig. 3b**). We then merged the filtered rFRC map (green channel) and RSM (red channel) to create the composite PANEL map (**Fig. 1b**). Note that our PANEL cannot fully pinpoint the unreliable regions induced by the model bias at present, which would require more extensive characterization and correction routines based on the underlying theory of the corresponding models^10, 28–30^.

### Validating with simulations

To test our quantitative maps with known ground truth, we used simulated datasets of SMLM from the EPFL challenge^11^ (**Methods**). These datasets consisted of high-density (HD, 361 frames) and low-density (LD, 12000 frames) emitters per frame to simulate excessively low or optimal illumination intensity conditions. The images were divided into two statistically independent subsets, yielding two SR reconstructions obtained using the maximum likelihood estimation (MLE)^31^, for our rFRC mapping (**Methods**, 4^th^ column in **Fig. 1c**). Between MLE reconstructions and ground-truth images, their differences in space indicate the locations and scales of different artifacts (first two columns in **Fig. 1c**). From this spatial difference and localization uncertainty maps (**Supplementary Fig. 5a**, **5b**), we found that the reconstructed SR image under the HD condition was much more blurred than that of the LD condition, possibly due to more overlapping emitters being excited simultaneously. This was exactly affirmed by the larger rFRC value (**Methods**) of the HD-MLE image than that of the LD-MLE image (0.66 versus 0.16, **Fig. 1c**), in which the rFRC map uncovered all the subtle errors (as pointed by white arrows, 4^th^ column in **Fig. 1c**). In contrast, the previous RSM cannot detect such subtle errors, and is influenced by noise-induced random intensity fluctuations (3^rd^ column in **Fig. 1c**). On the other hand, we note that rFRC failed to detect the filament’s missing part, mimicking defective local illumination or labeling (cyan arrows, **Fig. 1c**). That was revealed by the filtered RSM, highlighting the necessity of PANEL combination for pinpointing different types of errors (last column in **Fig. 1c**).

To demonstrate the dependence of the SMLM reconstruction quality on the illumination intensity, we synthesized a regular grid illuminated by a Gaussian beam with high power in the center and low power toward the edges (**Methods**, **Supplementary Fig. 5c**, **5d**). Under this circumstance, molecule blinkings at the center were better separated temporally than those at the edges^19^ (1^st^ column in **Fig. 1d**, **Supplementary Fig. 5d**), which was clearly revealed on the rFRC map (4^th^ column in **Fig. 1d**, 29 nm at the center and 80 nm at the edge). In contrast, because the space-invariant reconstructed PSF assumption did not hold up here, RSM provided incorrectly estimated errors (3^rd^ column in **Fig. 1d**, 0.40 a.u. at the center and 0.17 a.u. at the edge, **Supplementary Fig. 5e**), and it was opposite to the reference (2^nd^ column in **Fig. 1d**, 0.75 a.u. at the center and 1.00 a.u. at the edge).

Although the RSM is incompatible with volumetric datasets, the rFRC can be directly extended to a 3D version when applying plane-by-plane calculations (**Methods**). Here, we presented the simulated 3D dataset from the EPFL SMLM challenge^32^, including both LD and HD emitters (**Methods**, **Supplementary Fig. 6**). Similarly, compared to the reconstruction with LD emitters per frame, rFRC analysis demonstrated lower quality with HD emitters (3D rFRC value *LD:* 2.2, *HD:* 4.5, **Supplementary Fig. 6b**), confirming the real experimental experience.

### Evaluating resolution heterogeneity of localization microscopy

Next, we examined the experimental SMLM microtubule datasets (**Methods**, **Fig. 2a**, **Supplementary Fig. 7**). As visualized by the rFRC, the SR microtubule images obtained by large-field STORM^33^ (**Fig. 2a**, left), small-field SMLM^11^ (**Supplementary Fig. 7a**, **7b**), and SRRF^20^ (**Supplementary Fig. 7c**) demonstrated significantly lower resolutions at filament intersections (right at **Fig. 2a**, **Supplementary Fig. 7**) and perinuclear region of the cell (right at **Fig. 2a**). This is because the regions with more complex structures will exhibit more simultaneous emitters per area, inducing a relatively degraded resolution. In detail, as can be seen in **Fig. 2b**, the perinuclear region contains the most dense cytoskeleton, and its surrounding region is the transitional subregion with the microtubules becoming sparser. The peripheral subregion is further out, which exists in the thin perimeter areas of the cell^34^, and the microtubules appear as an expansive network. Interestingly, such three-stage structural distribution-induced resolution heterogeneity is successfully mapped by our rFRC (right at **Fig. 2a**, **Fig. 2b**). Overall, these rFRC maps offered a more intuitive interpretation for the real resolutions of the corresponding images from localization microscopy.

**Fig. 2.**
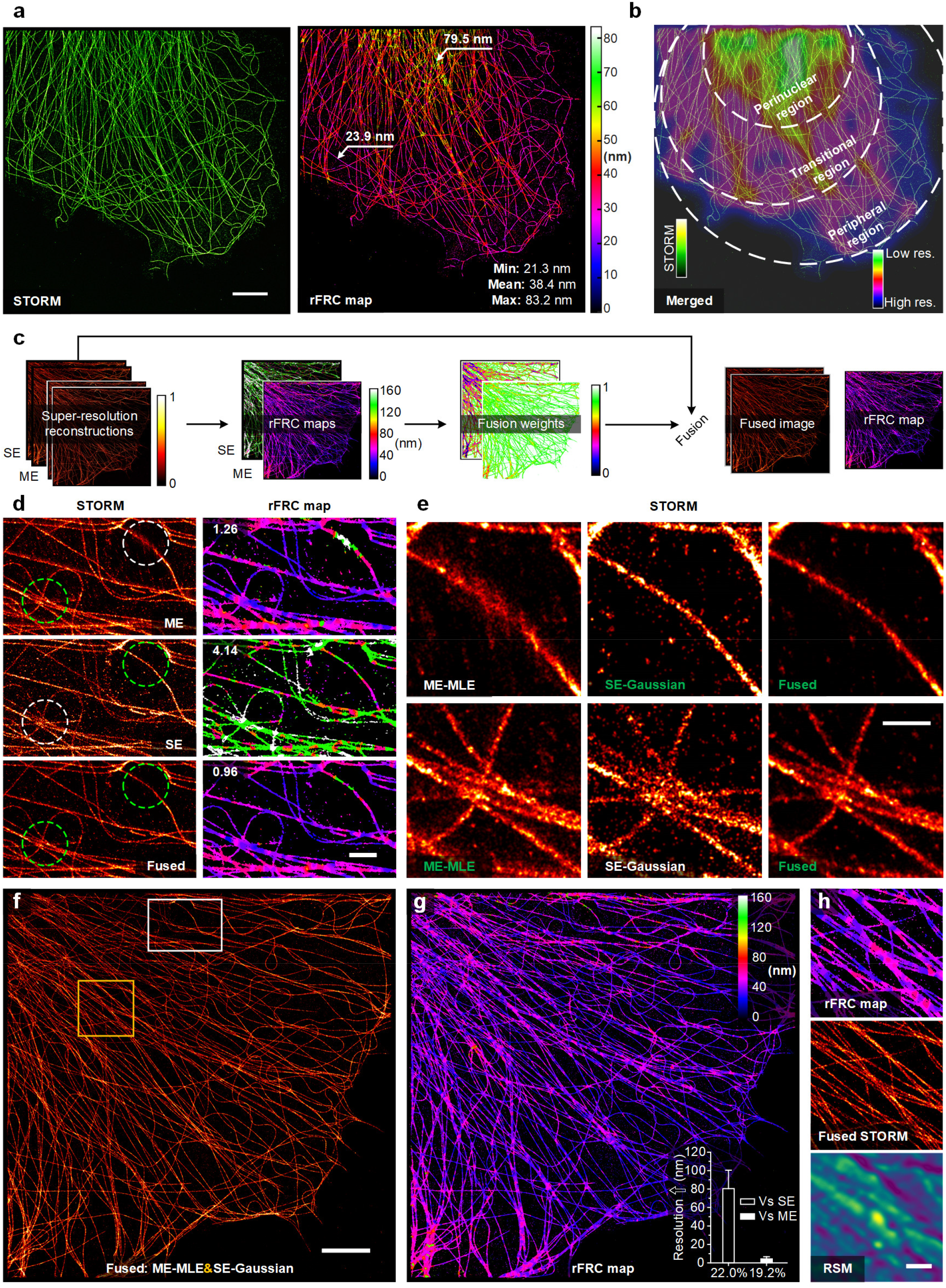
Evaluation and optimal fusion of STORM using the rFRC map. (**a**) STORM result of α-tubulin labeled with Alexa Fluor 647 in a COS-7 cell (left) and its rFRC map (right). (**b**) The merged view of STORM result (green hot) and Gaussian averaged rFRC map (shifted jet), highlighting the three-stage distribution. (**c**) Schematic of the STORM fusion. ‘ME’: Multi-emitter MLE result; ‘SE’: single-emitter Gaussian fitting result. (**d**) STORM results (COS-7 cells, α-tubulin labeled with Alexa Fluor 647, left) and their rFRC maps (right) are shown from top to bottom, which are magnified views of the white box in (**f**). From top to bottom: ‘ME’ result; ‘SE’ result; the fused result from the ‘ME’ and ‘SE’ reconstructions. The corresponding rFRC values are marked on the top left of the rFRC maps. (**e**) Magnified views of the dashed circles in (**c**). (**f**) The entire view of the fused STORM result (COS-7 cells, α-tubulin labeled with Alexa Fluor 647). (**g**) rFRC map of (**f**). The inset shows the improved resolution achieved by fusion compared with the SE (80.55 ± 1.52 nm at 22.0% region, hollow) and ME (4.28 ± 0.14 nm at 19.2% region, white solid) results. (**h**) Enlarged regions enclosed by the yellow box in (**f**). The results of the rFRC map, fused STORM, and RSM are shown from top to bottom. Error bars, s.e.m.; res.: resolution; scale bars: (**a**, **f**) 5 μm; (**d**, **e**) 500 nm; (**h**) 1 μm.

### Optimal fusion of SMLM

Since all current reconstruction algorithms assume homogenous HD or LD emitters per frame, the heterogeneity of resolution is becoming a major problem^19^. By identifying positions of high localization uncertainty with rFRC map, we can compare the local performances of different restoration algorithms, and fuse different regional reconstructions (**Methods**, **Fig. 2c**). To do so, the resolution heterogeneity and potential artifacts can be minimized. By integrating a high-density simulated dataset reconstructed by the multi-emitter MLE (ME-MLE)^31^ and the FALCON (fast localization algorithm based on a continuous-space formulation)^35^, the fused image demonstrated better PSNR (peak signal to noise ratio), SSIM (structural similarity), and rFRC values (**Supplementary Fig. 5f**-**5i**). To further evaluate its performance in real samples, we analyzed immunolabeled α-tubulin filaments in fixed COS-7 cells imaged with 2D-STORM and restored them with either the ME-MLE or single-emitter Gaussian fitting^33^ (SE-Gaussian) (**Methods**, **Fig. 2d**). Although the ME-MLE method performed better at approximating complex structures (HD emitters) and provided a lower overall rFRC value, the SE-Gaussian algorithm seemed to excel in reconstructing some simple structures (LD emitters) (**Fig. 2d, 2e**). By combining regions with the lowest local rFRC values between reconstructions from either of the two algorithms (**Methods**), the new composite SR image demonstrated better visual quality and the lowest overall rFRC value (0.96 versus 1.26 and 4.14, **Fig. 2d**). Moreover, the fused image exhibited a more homogenous distribution of spatial resolution than that obtained either by ME-MLE or SE-Gaussian alone (**Fig. 2f, 2g**), reinforcing its superior performance in the entire FOV. Specifically, this rFRC map-guided image fusion led to a substantially improved resolution (the inset in **Fig. 2g**) in replaced regions than the SE-Gaussian method (80.55 ± 1.52 nm, hollow), and significant increases in local resolutions than the ME-MLE method (4.28 ± 0.14 nm, white solid). In contrast, because the RSM method was incapable of revealing errors of SR ranges, it failed to identify such intricate structures from the STORM image (**Fig. 2h**).

Similarly, the rFRC was used to composite fusion to clathrin-coated pits (CCPs) in COS-7 cells under 2D-STORM. The merged SR image showed better quality and higher mean resolution (**Supplementary Fig. 8**). Beyond that, we further provided the rFRC map procedure for the 3D-STORM^33^ reconstruction (**Supplementary Fig. 9a**), in which significant uncertainties also occurred at interweaved filaments (**Supplementary Fig. 9b-9d**). Under this 3D configuration, the overall most accurate axial planes for microtubule (**Supplementary Fig. 9d**, **9e**) and actin filaments (**Supplementary Fig. 9f-9k**) were located at the focal planes.

### Evaluating diverse optical super-resolution microscopies

After establishing the validity and superiority of our method in SMLM, we extended our analysis to other non-pointillism SR methods. In theory, because the weight of the optical transfer function (OTF) decreases gradually with its spatial frequency, the noise will dominate the high-frequency components while the low-frequency deep inside the OTF support remains stable. The subsequent reconstructions will apply more amplifications at higher frequencies than lower ones, leading to significant fluctuations in high-frequency components, which renders intricate structures more profoundly affected by noise. Moreover, the variations of different SR reconstruction methods are usually on their SR scale, and thus an evaluation on the corresponding level is essential. Here, our rFRC offers a well-timed solution to detect these uncertainties at high spatial frequencies.

Hessian-SIM^36^ using the Hessian matrix continuity on the Wiener-SIM^37^ results to reduce random, non-continuous artifacts induced by the live-cell low SNR imaging conditions. We applied the rFRC map to differentiate such subtle variations in fidelity between conventional Wiener-SIM and Hessian-SIM (rFRC value, 1.36 versus 1.24) (**Methods**, **Supplementary Note 2**), and in contrast, the RSM detected identical qualities (RSE value, 0.27 versus 0.27). Richardson-Lucy (RL) deconvolution^22, 23^ has been well applied in many optical microscopes^13^ to improve the image contrast, deblurring the estimate of sample density with each iteration. However, the RL deconvolution may amplify noise under excessive iterations, and thus require back-and-forth visual inspection for iteration determination. With a finer assessment, we used rFRC map to moderate this challenge in two aspects. First, we applied RL deconvolution to process the total internal reflection fluorescence (TIRF) image and then calculated its corresponding rFRC value of each iteration to determine the optimal iteration times objectively (**Supplementary Note 3.1**). Second, we performed RL deconvolution with excessive iterations on a simulated wide-field image, in which the amplified noise induced snowflake-like artifacts. To adaptively filter the high-frequency artifacts, we used local cutoff frequencies determined by the rFRC map to low-pass filter various block-box areas within the entire image, producing optimal quality against the global filter (**Supplementary Note 3.2**). Finally, we even employed our evaluation on coherent imaging reconstructions, and obtained an accurate quality rating map contrasted to ground-truth, pinpointing all subtle untrustworthy regions (**Supplementary Note 4**).

### The model and data uncertainties in learning-based applications

Characterizing the uncertainty of the corresponding network representations is crucial for further quantitative analysis. The Bayesian neural network (BNN) framework^38^ can be trained to obtain the uncertainties by learning the distribution overweight (**Supplementary Note 7.2**), and two types of uncertainty are defined in its concept, i.e., the data uncertainty and the model uncertainty. Generally, the data uncertainty is induced by the combined effects of noise or sampling, and the model uncertainty is caused by the existing distance between the established model and its real-world counterpart. Drawing on approximate implementations of BNN^38, 39^, in several learning-based microscopies, the data and model uncertainties can be measured by modifications to the original training procedures and network architecture^40–44^. Notably, the data uncertainty is learned and predicted by the network itself using distributional approximations^40–42, 44^. Thus it is hard to ensure the stability and rationality of the obtained data uncertainty^44^, and these modifications may compromise the network performance.

In optical SR modalities, we have shown how the rFRC mapping (from two independent captures) model-independently measured the data uncertainties with no need for modifications. When considering the learning-based restorations, we also envisage that the model uncertainty can be directly detected by the ensemble disagreement^39^ of independently training repeated models on the same dataset with multiple random initializations and optimizations^43^. By applying the rFRC map to two predictions from two inputs (data sampling) and two models (network training), respectively, we can monitor both the data uncertainty and the model uncertainty (**Supplementary Note 7.2**). To test our strategy, two deep neural networks (*f*_A_ and *f*_B_) were trained independently, with high-resolution images (120 nm PSF) as ground-truth and corresponding low-resolution (240 nm PSF) as input (**Fig. 3a-3e**). When we supplied input images of different resolutions, this represented out-of-distribution data since the image transformations were specific to the 120 nm PSF and 240 nm PSF data pair. Therefore, to artificially increase the potential model uncertainty, we created 300 nm PSF images as the out-of-distribution test data.

**Fig. 3.**
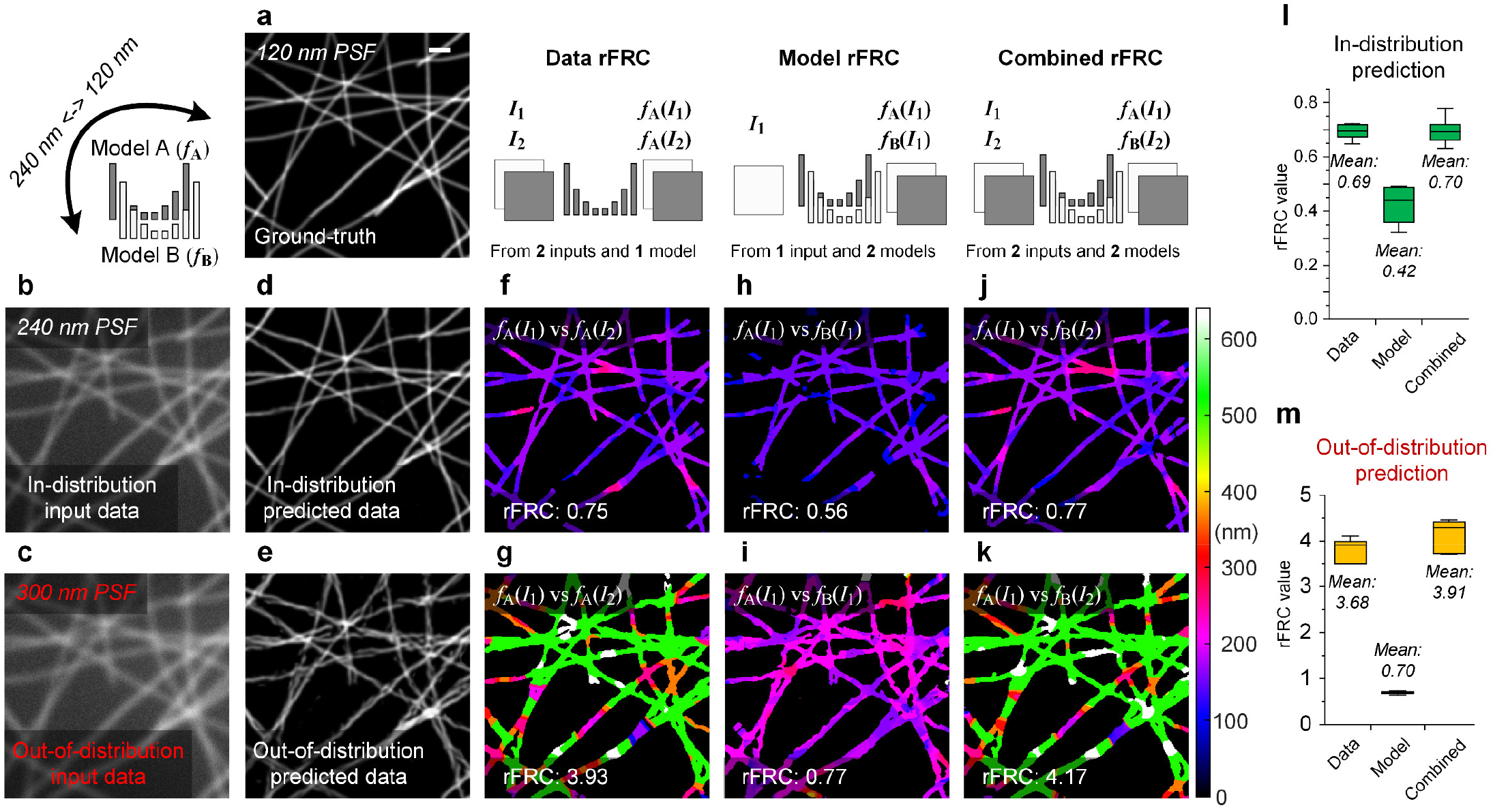
Data and model uncertainty quantifications of learning-based restoration. (**a**) The synthetic tubulin structures were convoluted with a 120 nm PSF and down-sampled 2 times (pixel size 40 nm) as ground truth. (**b**) The structures were convoluted with a 240 nm PSF, and down-sampled 2 times before adding 10% Gaussian noise to be the training dataset and in-distribution test image. (**c**) The structures were convoluted with a 300 nm PSF, and down-sampled 2 times before adding 10% Gaussian noise to be the out-of-distribution test image. (**d**, **e**) The network predictions of in-distribution input (**d**, rFRC value: 0.75; rFRC resolution: 161.0 nm), and out-of-distribution input (**e**, rFRC value: 3.93; rFRC resolution: 453.2 nm). (**f**, **g**) The ‘Data rFRC’ maps of two predictions from two in-distribution inputs (**f**), and two out-of-distribution inputs (**g**). (**h**, **i**) The ‘Model rFRC’ maps of two model predictions from in-distribution input (**h**, rFRC value: 0.56; rFRC resolution: 143.2 nm), and out-of-distribution input (**i**, rFRC value: 0.77; rFRC resolution: 203.2 nm). (**j**, **k**) The ‘Combined rFRC’ maps of two model predictions from two in-distribution inputs (**j**, rFRC value: 0.77; rFRC resolution: 163.0 nm), and out-of-distribution inputs (**k**, rFRC value: 4.17; rFRC resolution: 475.6 nm). (**l**, **m**) rFRC values of in-distribution and out-of-distribution predictions. Scale bar: (**a**) 1 μm.

#### Data uncertainty evaluated by rFRC (Data rFRC)

Here we sampled the input image twice (*I*_1_ and *I*_2_), yielding two corresponding SR reconstructions (*f*_A_(*I*_1_) vs. *f*_A_(*I*_2_)) from the same model (*f*_A_) (**Fig. 3f, 3g**). The result from 240 nm PSF input (240-result) has a rather small rFRC resolution distribution, indicating a high-quality SR reconstruction (**Fig. 3f**). On the other hand, upon inputting the out-of-distribution data, i.e., 300 nm PSF, the result (300-result) contained much more hallucinations, such as smooth filaments became broken pieces due to the enlargement of structures by the 300 nm PSF (**Fig. 3e**). This is highlighted by the ‘Data rFRC’ mapping demonstrating lower resolution distribution, especially at the intersections of the filaments (**Fig. 3g**).

Notably, because the deep models have no stationary form with only learning the representations of training data, the model uncertainty and data uncertainty will be not mutually exclusive^38^. Compared to the correct (in-distribution) input data, the predictions from the out-of-distribution input data are more sensitive to noise or other potential influences, leading the predictions more prone to fluctuations. Therefore, the model uncertainty can leak into the data uncertainty^38^, which allows our rFRC to indirectly detect the leaked model uncertainty from the data uncertainty (see also another univocal example in **Supplementary Note 7.1**). Although our rFRC method only examined pure data uncertainty in **Fig. 3f** and **3g**, we successfully appreciated the potential hallucinations induced by the out-of-distribution data, explicitly showing the leakage of model uncertainty into the data uncertainty.

#### Model uncertainty evaluated by rFRC (Model rFRC)

To detect the model uncertainty directly, we applied the rFRC mapping on two network output images (*f*_A_(*I*_1_) vs. *f*_B_(*I*_1_)) from the same input (*I*_1_), in which *f*_A_(*I*_1_) and *f*_B_(*I*_1_) are predicted from two models trained repeatedly (*f*_A_ and *f*_B_) (**Fig. 3h, 3i**). The network was nearly free from the model uncertainty when inputting the in-distribution data. The corresponding structures were predicted accurately (**Fig. 3d**), confirmed by the high-resolution distribution from the ‘Model rFRC’ mapping (**Fig. 3h**). When presenting the network with the out-of-distribution images, the predictions still approximated the corresponding structures from the expected ‘240 nm to 120 nm’ transformation (**Fig. 3e**). As a result, the corresponding rFRC map represents an overall lower resolution distribution, indicating the model uncertainty increased by networks’ ignorance of the out-of-distribution data (**Fig. 3i**). Interestingly, compared to the ‘Data rFRC’ (rFRC values, 0.75 vs. 3.93), we found the overall resolution decrease of ‘Model rFRC’ induced by the out-of-distribution test is relatively small (rFRC values, 0.56 vs. 0.77), and it might reflect that most model uncertainty has been leaked to the data uncertainty.

#### Combined uncertainty evaluated by rFRC (Combined rFRC)

To monitor the combination of both data and model uncertainties, we used the rFRC mapping on the two predictions (*f*_A_(*I*_1_) vs *f*_B_(*I*_2_)), in which we employed two inputs (*I*_1_ and *I*_2_) for two models trained repeatedly (*f*_A_ and *f*_B_) (**Fig. 3j, 3k**). We found that the overall distributions of these rFRC maps are highly consistent with the results in ‘Data rFRC’. The metrics of ‘Data rFRC’ and ‘Combined rFRC’ from in-distribution predictions are almost identical (**Fig. 3l**), in which the mean rFRC values are 0.69 and 0.70, respectively. This suggests that the model uncertainty of in-distribution input is close to zero. In contrast, when examining the out-of-distribution predictions, the metrics of ‘Combined rFRC’ are slightly larger than that of ‘Data rFRC’ (**Fig. 3m**), in which the mean rFRC values are 3.68 and 3.91, respectively. The difference (3.68 versus 3.91) may indicate that there is still a tiny amount of residual model uncertainty after its leaking into the data uncertainty.

### Assessments of learning-based restorations

To illustrate the general applicability, we applied rFRC to several learning-based SR microscopes with experimental datasets, evaluating the local qualities of the network-extrapolated SR information (**Fig. 4**). For example, artificial neural network accelerated PALM (ANNA-PALM)^45^ reconstructs dense images from sparse localization images, significantly reducing the total number of raw frames. In ANNA-PALM reconstruction, MLE reconstructions of individual molecules within 25 frames were used as the input (**Methods**, **Fig. 4a-4e**, **Supplementary Fig. 10**), and the resultant SR image (**Fig. 4a**) resembled those generated by MLE of all molecules emitted within 500 frames (**Fig. 4b**). In this case, the experimental configurations of our tested input^11^ are very different from the open-sourced model^45^, and thus this out-of-distribution input will induce a large model uncertainty. Although the repeatedly trained models are unavailable, the leaked model uncertainty can be detected by our rFRC map (using two captures) indirectly with the RSM contributing to perceiving the remnant part. We found rFRC map (**Fig. 4c**) can indicate all the subtle reconstruction uncertainties at filament intersections (cyan arrows, **Fig. 4e**), with the RSM finding the missing bulk structures (magenta arrows, **Fig. 4e**). All these local restoration qualities were assessed (**Fig. 4d**) without the 500-frame MLE image, and they were confirmed by comparing the ANNA-PALM image (**Fig. 4a**) with the ground truth (**Fig. 4b**).

**Fig. 4.**
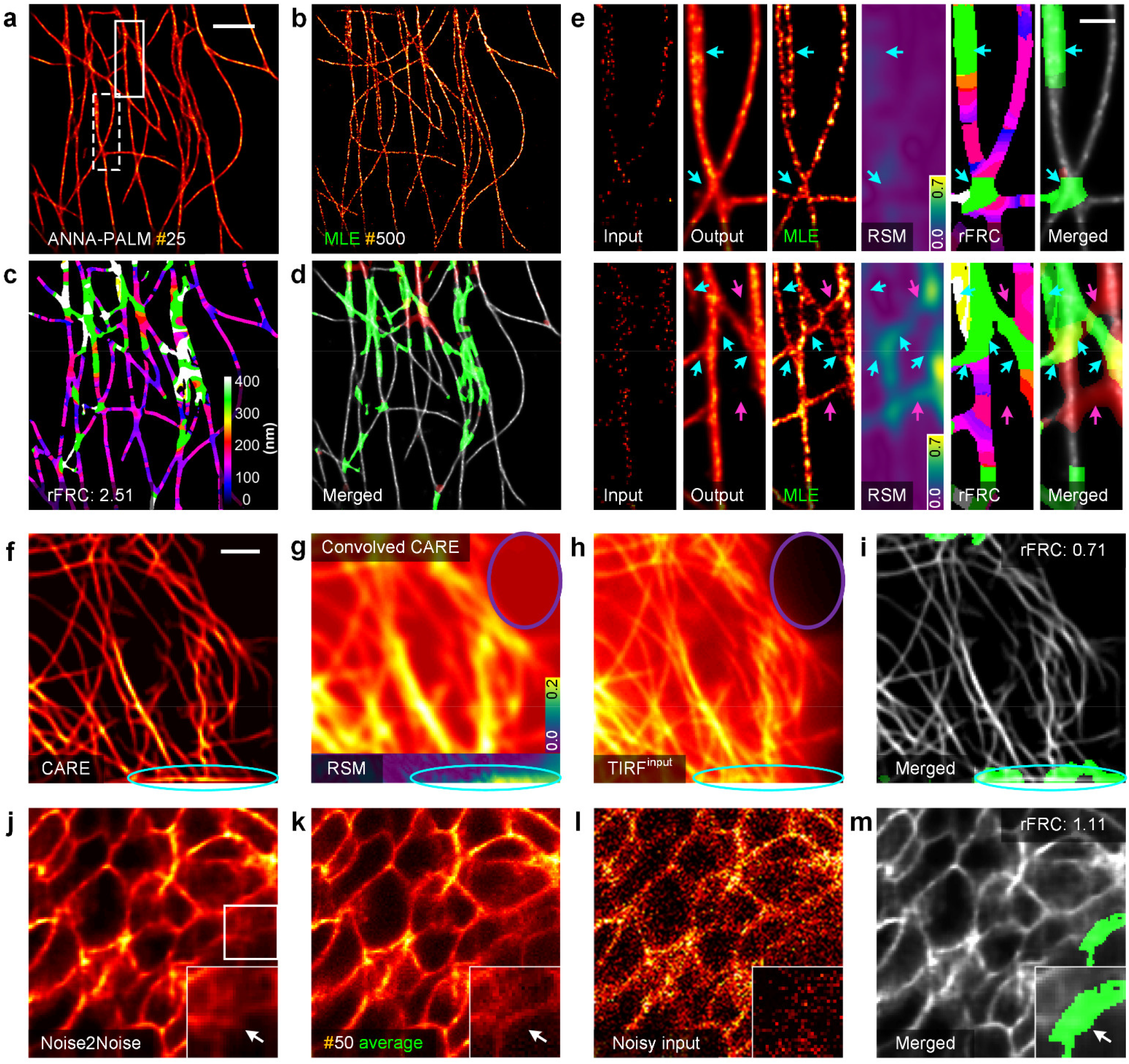
Assessments of diverse learning-based reconstructions. (**a**) ANNA-PALM output (MLE reconstruction with 25 frames of tubulin as input). (**b**) MLE reconstruction with full 500 frames. (**c**) rFRC map of (**a**). (**d**) Merged image of the PANEL (green channel) and ANNA-PALM (gray channel) results. (**e**) From left to right: Enlarged regions of input sparse MLE reconstruction, ANNA-PALM output, full dense MLE reconstruction, RSM, rFRC map, and merged PANEL visualization map from the white solid (top) and dashed (bottom) box in (**a**). Cyan and magenta arrows represent the errors detected by rFRC and RSM, respectively. (**f**) CARE output of GFP-tagged microtubules in live HeLa cells (raw TIRF image as input). (**g**) CARE convolved back to its original low-resolution scale (top) and its RSM (bottom) result. (**h**) Corresponding TIRF image. (**i**) Merged image of the PANEL (green channel) and CARE (gray channel) results. (**j**) Noise2Noise result of EGFP-labeled *Tg (sox10:megfp)* zebrafish at 2 days postfertilization. (**k**) Ground-truth reference image generated by averaging 50 noise images with identical content. (**l**) Representative noisy input. (**m**) Merged image of PANEL (green channel) and Noise2Noise (gray channel) results. Centerline, medians; limits, 75% and 25%; whiskers, maximum and minimum; error bars, s.e.m.; scale bars: (**a**, **f**) 2 μm (**c**) 500 nm.

The content-aware image restoration (CARE)^40^ network framework enables more effective denoising and deconvolution. We directly reproduced the results using the open-sourced model and data^40^ (**Methods**), and thus the corresponding predictions should be nearly free from the model uncertainty (**Fig. 4f-4i**). The rFRC from two captures successfully detected the unreliable regions (cyan circle in **Fig. 4i**), in which the laterally displaced microtubules predicted at the boundary were due to edging effects but not real structures (cyan circle in **Fig. 4f**). Interestingly, by processing the predicted image back with a presumably space-invariant PSF and background (**Supplementary Fig. 11**), we identified excessive background fluorescence in a region (purple circle in **Fig. 4g**), absent in both the original image (**Fig. 4f**) and the TIRF reference (**Fig. 4h**), and our truncation operation on the RSM eliminated this potential false negative (the absence of red-color components in **Fig. 4i**).

Next, we explored the capability of our method in evaluating the noise-removal effects of the learning-based approaches. For example, Noise2Noise is a widely known unsupervised method that is superior in denoising noisy images without needing clean images^46^. Here we used fluorescence microscopy denoising (FMD)^47^ datasets to train the Noise2Noise network (**Methods**, **Fig. 4j-4m**, **Supplementary Fig. 12**). Briefly, we fed the network with two images with independent noise but identical structure details, one for the input and the other for the output target. Because the corresponding wide-field reference of this task is unavailable, we used the rFRC map only. Interestingly, the suspicious area (in green) was amplified in the rFRC map (**Fig. 4m**), and it was also verified by the least overlapped region between the model prediction and the average of 50 noisy images (**Fig. 4j-4k**, white arrows). Finally, by applying a single-frame rFRC strategy, we also showed that it could reveal the hallucinations in learning-transformed SR-SIM images from wide-field images (**Methods**, **Supplementary Note 8.2**).

## DISCUSSION

In deep learning applications, more and more microscopists have noticed the importance of model uncertainty. However, different from the *high-level* vision tasks, the learning-based microscopies intend to inverse the degraded images to their high-quality counterpart. This *low-level* forward (image degradation) and backward (learning to inverse the degradation) process will be more affected by the data qualities. We demonstrate that most model uncertainty has leaked into the data uncertainty in **Fig. 3** and **Supplementary Note 7.2**. This phenomenon suggests that the data uncertainty is even more crucial for learning-based microscopies.

According to the underlying theories of different modalities, the corresponding model uncertainties can be minimized by optical system calibrations^10, 29, 30^, or suppressed by a specifically designed learning strategy^28^ and enough training data^38^. On the other hand, data uncertainty is fundamentally inevitable and difficult to remove, and there is still no effective method for its routine evaluation. In this work, we use rFRC with two independent captures to measure the data uncertainty in general, and in particular for learning-based applications, we also provide a strategy to reveal both model and data uncertainties.

Without a reference, a map of uncertainty down to the SR scale will be crucial for extracting reliable and quantitative information from biological images. When the spatially different uncertainties revealed by rFRC, the way may be paved for these state-of-the-art imaging methods to be widely adopted in cell biological studies. Based on our analysis (**Fig. 2**), we uncover that the resolution heterogeneity can be a sought-after issue to be discussed in future methodological developments and even biological studies. Assisted by rFRC, we anticipate the developers and users can optimize the resolution heterogeneity and evaluate the performances for specific experiments. In addition, we also expect our rFRC can be broadly used as a cross-modality tool, evaluating the resolution heterogeneity for other typical localization microscopies, such as ultrasound localization microscopy^48^ and the recently emerged localization atomic force microscopy^49^, offering well-founded systemic improvement schemes.

When two independent frames are unavailable, we also introduced two alternative single-frame strategies (**Supplementary Note 8**). For optical imaging modalities, we followed ref^50^ and divided a single frame into four subsets to create two image pairs for the rFRC calculation (details in **Supplementary Note 8.1**). For learning-based methods, we added independent noise to raw data to create the required two-frame input (details in **Supplementary Note 8.2**). Finally, to avoid the potential false negative, it is suggested that the rFRC mapping skipped the background areas, requiring the background threshold determination. In addition to the global threshold background filter strategy, we adopted an adaptive method for local thresholds calculation to adapt to more modalities (**Methods**, **Supplementary Fig. 2c**). Regarding the modalities without the requirement of additional postprocessing, the rFRC map can also provide a fine resolution-map for imaging quality evaluation and further optimization (**Supplementary Note 5**). For 3D data, we applied rFRC mapping on volumetric datasets in a slice-by-slice manner to visualize the quality variations on each plane. The 3D extension of our method would require 3D rolling operation and the Fourier shell correlation (FSC)^25^ calculation to further incorporate axial information.

In addition to the reference-free objective quality rating, we also expect our rFRC map can become a generalized metric in the presence of ground-truth, similar to the structural similarity (SSIM)^51^, to assess image quality closer to the human perception (**Supplementary Note 6**), which may be equally important in the SR microscopy field. By developing an open-source ImageJ plug-in, and libraries in different programming languages, we enable wide users to apply our method. We hope this metric will benefit image-based biological profiling and inspire further advances in the rapidly developing field of computational microscopes.

## METHODS

### FRC calculation

The FRC method measures the statistical correlation between two bidimensional signals over a series of concentric rings in the Fourier domain. It can be regarded as a function of the spatial frequency *q_i_*:

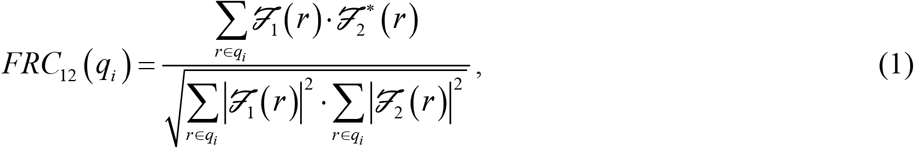

where 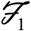 and 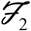 denote the discrete Fourier transforms (DFTs) of the two signals and 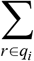 represents the summation over the pixels on the perimeter of circles of corresponding spatial frequency *q_i_*.

Before calculation, a Hanning window is used to suppress the edge effects and other spurious correlations caused by the DFT calculation. The rectangular images should be zero-padded to produce squares to calculate the FRC curve. To calculate the discrete values of the corresponding spatial frequencies, it is necessary to define the discretization of the spatial frequencies of the FRC curve. The maximum frequency *f*_max_ is half the inverse of the pixel size (*p_s_*), i.e., *f*_max_ = 1/(2 *p_s_*). Then the average filter with a half-width of the average window (equal to 3 frequency bins) is applied to smooth this noisy FRC curve.

When the FRC curve drops below a given threshold, the corresponding frequency is defined as the effective cutoff frequency (COF), whereas the resolution is the inverse of the effective COF. This threshold for FRC indicates the spatial frequency above which meaningful information beyond random noise can be extracted. Specifically, the common choices for the criterion/threshold are the fixed-value thresholds or the sigma-factor curves^25^. The fixed value is usually the 1/7 hard threshold, and the criterion of sigma-factor curves can be written as follows:

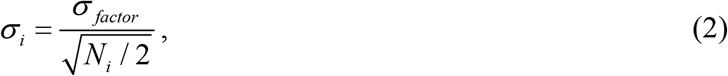

where *N_i_* represents the number of pixels in a ring of radius *q_i_* and the most commonly used *σ_factor_* is 3. If the two measurements are corrupted with excessive noise, the FRC curve can be expressed as 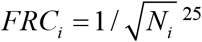.

The 1/7 hard threshold has been widely used in determining the resolution of SR images. Although this fixed-value threshold method is incompatible with statistical assumptions^25^, the resolution obtained with that criterion is approximately accurate for SMLM^14^ and the stimulated emission depletion microscopy (STED) microscopy^52^. The 1/7 threshold attains a similar result for a large image to the 3*σ* curve criterion (**Supplementary Fig. 15a**). However, this fixed threshold is overconfident for determining the resolutions of small image blocks, which is essential to map local SR errors in the reconstructions. In **Supplementary Fig. 15a**, the 1/7 threshold is smaller than all correlation values in the FRC curve and fails to yield the COF of small images (red cross). On the other hand, unlike avoiding the conservative threshold choice in resolution determination, we prefer a moderate threshold for quality mapping to reduce false positives. Therefore, we choose three standard deviations above the expected random noise fluctuations as the threshold^25^. This criterion is robust and accurate in examining small image blocks and calculating the FRC resolutions.

### rFRC map generation

#### Two-frame generation

The rFRC mapping requires two independent frames of identical contents under the same imaging conditions. For the SMLM and the SRRF modalities (**Fig. 1-2**, **Fig. 4a-4i**, **Supplementary Fig. 5-10**), these two frames were generated by splitting the raw image sequence in half (odd and even frames) and reconstructing the resulting two image subsets independently. For the SIM, FPM, RL deconvolution, STED (**Supplementary Fig. 18-22**), and some learning-based methods (**Fig. 3**, **4j-4m**), we directly imaged the identical contents twice to capture the required two frames. Regarding the two-frame unavailable configurations, we also provided two alternative strategies to produce two frames from the single accessible image in **Supplementary Note 8**.

#### rFRC Mapping

Since the FRC measures the global similarity between two images, we extend the FRC to a rolling form (rFRC) to provide the local distance measurements at the pixel level. We regard the FRC calculations as a filter in which the image is scanned block by block (64 × 64 pixels as a default size in this work), with each block assigned the corresponding FRC resolution. First, we pad the input image symmetrically around a half size of the block to calculate the FRC at the image boundaries (Step 1, **Fig. 1a**). Second, by setting the background threshold of the center pixel, we avoid FRC calculation of the background area. If the mean of the center pixels is larger than the threshold, we calculate the FRC and assign the FRC resolution to the center pixel of each block. In contrast, we set a zero value to the central pixel when it is smaller than the threshold (Steps 2-4, **Fig. 1a**). Afterward, we run this procedure block by block until the entire image is finished.

#### Background thresholding

By labeling designated structures specifically, fluorescence images confer high contrast and dark background areas containing background and readout noise. These regions, however, result in low FRC resolutions that are essentially false negatives. Therefore, we use two strategies to threshold the background (**Supplementary Fig. 2**). We determine the hard threshold according to the images by user-defined global value adapting to their data (default method) or by an iterative wavelet transform method^53^ to estimate local values automatically. For the global threshold, because different values lead to different regions being interrogated, we choose the hard threshold carefully based on two principles: 1) the removal of background; 2) the maintenance of structures. Regarding the local threshold, the background is iteratively estimated from the lowest-frequency wavelet bands of the images (**Supplementary Fig. 2c**). In each iteration, all image values above the current estimation are clipped.

#### rFRC mapping acceleration

Although the rFRC allows evaluation at the pixel level, the most delicate scale of detectable errors can only reach the highest resolution allowed by the system, which satisfies the Nyquist-Shannon sampling theorem. Thus, the smallest error should be larger than ~3 × 3 pixels. Therefore, we can skip 2~4 pixels for each rolling operation to accelerate the mapping calculation 4~16 times. The rFRC map can be resized to the original image size by bilinear interpolation for better visualization.

#### Adaptively filtering the rFRC map

The FRC calculation is not always stable and may generate aberrantly large values in neighboring pixels due to improperly determined COFs. Thus, we create an adaptive median filter to remove these inappropriate values. Instead of the standard median filter that replaces each pixel with the median of the neighboring pixels, we develop an adaptive median filter to remove only the isolated pixels with aberrantly large values, avoiding blurring of the rFRC map^13^. If the pixel intensity is larger than a preset fold (default as 2-fold) of the median in the window (default as 3-pixel), the pixel is replaced by the median value. Otherwise, the window moves to the next pixel.

#### Drift correcting

To correct relative movements between measurements, we use a method based on the phase correlation^54^. First, we calculate the cross-correlation function *CC* of the two images:

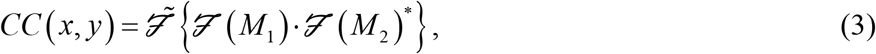

where *M*_1_ and *M*_2_ represent the two images. The peak of the *CC* is the shift between these two images that ensures the best-correlated *M*_1_ and *M*_2_. After that, we find the centroid of the distribution of intensities of the cross-correlation function to achieve subpixel accuracy. This operation is executed before the rFRC mapping.

### rFRC colormap

Choosing a proper color map to visualize error maps is another tricky question. The existing popular color maps, such as Jet, use blue to red to index the different error magnitudes. However, people usually tend to define black (dark color) as small magnitude and white (light color) as large magnitude, which is identical to the logic of the gray color map. In this sense, the Jet color map may be incompatible with human intuition^26^. On the other hand, human vision is insensitive to light or dark gray levels and sensitive to different colors. As a result, we intend to create a color map using color to index the magnitudes and with black/white zone to visualize the smallest/largest values.

First, because human eyes are more sensitive to green color, we use green to highlight errors of large magnitude. Second, human instinct usually regards bright color (white) as an effect of large magnitude and dark color (black) for small magnitudes. Therefore, we involve a black zone (0, 0, 0) and a white zone (1, 1, 1) in the color map to visualize the smallest and largest values. Taken together, we shift the Jet colormap (left panel of **Supplementary Fig. 3a**) to create the shifted Jet (sJet) color map (right panel of **Supplementary Fig. 3a**). Along with the extension of the blue color component in this sJet color map, we obtain a white zone to represent the most significant error (even larger than those highlighted in green). Because the background in the rFRC map means no error, we use the black zone for the display. As shown in **Supplementary Fig. 3a**, our sJet color map is more intuitive for visualizing errors than the original Jet color map.

In addition to the sJet colormap, we also provided another alternative colormap, i.e., Jet with the black zone (bJet, middle panel of **Supplementary Fig. 3a**) while using red color to represent large magnitude. The readers are encouraged to try these colormaps and select their favorite ones.

### rFRC value

As mentioned above, the rFRC map can be used to subtly visualize the local uncertainties down to the SR scale. Here, we also intend to give two metrics for globally evaluating the entire image quality. One metric with dimension *(resolution)* represented the averaged resolution across the entire imaging field, namely rFRC resolution, and its calculation is given as follows:

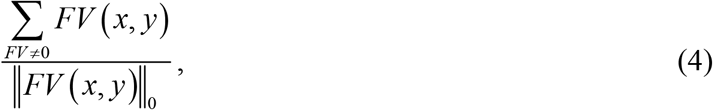

where ||*FV*||_0_ is the *l*_0_ norm, which represents the number of nonzero values in the rFRC map, and *FV* denotes the rFRC map.

Secondly, to reflect the potential deterioration rate of the reconstructed images, we provided a more generalized dimensionless metric, namely rFRC value. Here we normalize the rFRC resolution with its corresponding minimum resolution, and subtract 1 to ensure its value starting at 0:

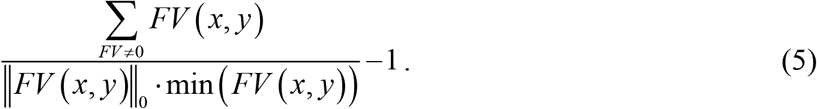

It noted that both metrics can be further extended to three dimensions, in which the (*x*, *y*) two-dimensional coordinates can be raised to three dimensions (*x*, *y*, *z)* directly (3D rFRC value).

### RSM generation

#### Image intensity rescaling and resolution scaling function (RSF) estimation

To normalize the intensity between low-resolution (LR) and high-resolution (HR) images and maximize the similarity between them, the intensity of the original HR image *I_H_* needs to be linearly rescaled:

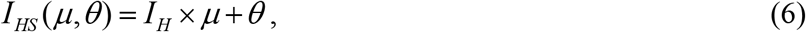

where *I_HS_* represents the HR image after linear rescaling. The values of *μ* and *θ* in **Eq. (6)** should be chosen to maximize the similarity between the LR image, *I_L_,* and *I_HS_* convolved with the RSF. Because the RSF is an unknown kernel used to transform an HR image into an LR image, it can be approximatively defined by a 2D Gaussian function with an unknown *σ.* The RSF is usually anisotropic in the *x* and *y* directions. Hence unlike its original version^19^, we set *σ* as a vector that includes two elements, i.e., *σ_x_* and *σ_y_*.

Then, to estimate *μ* and *θ* for image intensity rescaling and *σ_x_* and *σ_y_* for RSF parameterization, we jointly optimize these four variables (**Supplementary Fig. 4**), i.e., *μ*, *θ*, *σ_x_*, and *σ_y_*, to minimize the following function:

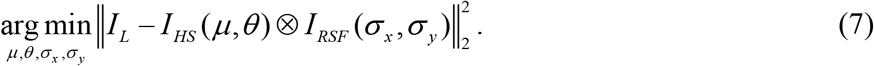

Because the gradient in **Eq. (7)** is difficult to calculate, we use a derivative-free optimizer to search for the four optimal parameters. Different from the particle swarm optimization (PSO)^55^ used previously^19^, we chose the pattern search method (PSM)^56^ to optimize **Eq. (7)**. PSO searches for substantial candidate solutions and may not be necessary for a four-parameter optimization problem. Compared to the unstable and slow metaheuristic optimization approach of PSO, the PSM is stable, computationally effective, and direct. It is commonly used in small-scale parameter optimization problems and is more suitable for our RSM estimation.

#### Metrics and pixel-wise error map of the RSM

After obtaining *μ* and *θ* (image intensity rescaling factors) and *σ_x_* and *σ_y_* (RSF parameters), we can transform the HR image *I_H_* into its LR version *I_HL_* by convolving the estimated RSF.

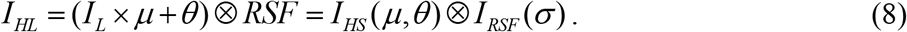

To assess the global quality of the resolution-scaled-back image *I_HL_* against the original LR image *I_L_*, we use the common root mean squared error for the resolution-scaled error (RSE)^19^ and the Pearson correlation coefficient for the resolution-scaled Pearson coefficient (RSP)^19^.

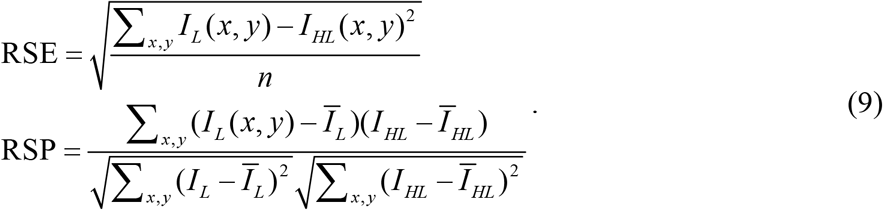

In addition, to visualize the pixelwise absolute difference, the RSM between *I_L_* and *I_HL_* can be calculated by:

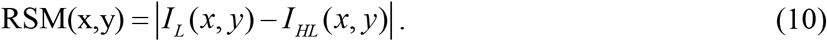

### PANEL pinpointing

To pinpoint regions with a high probability of error existence, we filter both the RSM and the rFRC to create a PANEL composite map. The small-magnitude components contained in the RSM may introduce false negatives. Therefore, we segment the RSM before integrating it into PANEL by the following equation:

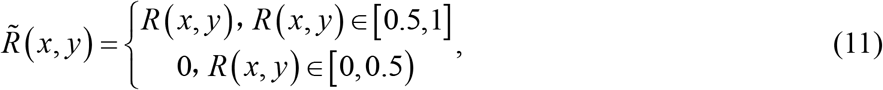

where *R*(*x, y*) represents the normalized RSM value in the *x, y* positions and 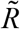 denotes the segmented RSM. After this operation, the small false negative is filtered, leaving us with strong low-resolution scale error components, focusing on the true negatives detected by the RSM. On the other hand, the rFRC map indicates the degree of uncertainty. The smallest FRC value in the map may not represent the error existence. Likewise, we introduce a segmentation method called Otsu^27^, which automatically determines the threshold by maximizing the interclass variance, performing image thresholding to filter the background in the rFRC map, and highlighting the regions with a high possibility of error existence (**Supplementary Fig. 3b**).

After that, considering human eyes more sensitive to the green color, we used the rFRC as green channel for better visualization of fine details, and leave the red channel for RSM to display large-scale components. In detail, first, the rFRC map and the RSM are normalized to a 0~1 scale. Second, we filter the rFRC map and the RSM with the ‘Otsu determined threshold’ and the ‘0.5 threshold’, respectively. Regions with values smaller than the thresholds are set to zero, and regions with larger values remain unchanged. Finally, we merge the rFRC map (green channel) and the RSM (red channel), and this operation is for qualitative pinpointing of regions with low reliability. The original rFRC map and the 0.5 threshold filtered RSM can be separated if quantitative evaluations are required.

In addition, if the datasets are three-dimensional or under a non-Gaussian convolution relation (between the low-resolution and high-resolution scales), we cannot estimate the corresponding RSMs. For these datasets, the RSM is not integrated into PANEL.

### SMLM Fusion

The RSM estimates the errors at the low-resolution scale, which is not suitable for the SMLM fusion. In contrast, the rFRC estimates the degree of errors at the SR scale and thus is a superior choice to guide the fusion of SMLM. Using the rFRC quality metric, we can fuse different localization results according to the weights of the rFRC maps, resulting in combined reconstructions that perform better than any one of the reconstructions alone.

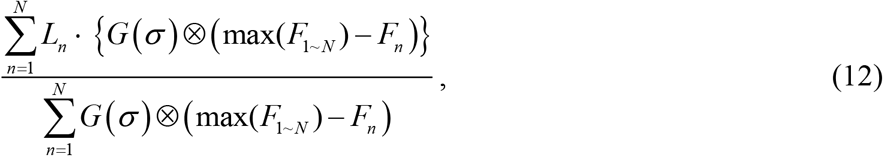

where *L_n_* is the result of the *n^th^* localization model, and *G*(*σ*) represents the Gaussian kernel with *σ* standard variance. The max(*F_l~n_*) is the maximum FRC value of the total *N* localization results, and ⊗ is the convolution operation. We use *G* (*σ* as 4 pixels) to slightly blur the rFRC map, avoiding oversharpen effects.

### STORM imaging

#### Microscope setup

After washing with phosphate buffer saline (PBS), the samples were mounted on glass slides with a standard STORM imaging buffer consisting of 5% w/v glucose, 100 × 10^-3^ M cysteamine, 0.8 mg mL^-1^ glucose oxidase, and 40 μg mL^-1^ catalase in Tris-HCl (pH 7.5)^33^. Then, data were collected by 3D-STORM^33^ carried out on a homebuilt setup based on a modified commercial inverted fluorescence microscope (Eclipse Ti-E, Nikon) using an oil-immersion objective (100×/1.45 NA, CFI Plan Apochromat λ, Nikon). Lasers at 405 nm and 647 nm were introduced into the cell sample through the objective’s back focal plane and shifted toward the edge of the objective to illuminate ~1 μm within the glass-water interface. A strong (~2 kW cm^-2^) excitation laser of 647 nm photoswitched most of the labeled dye molecules into a dark state while also exciting fluorescence from the remaining sparsely distributed emitting dye molecules for single-molecule localization. A weak (typical range: 0–1 W cm^-2^) 405 nm laser was used concurrently with the 647 nm laser to reactivate fluorophores into the emitting state. Only a small, optically resolvable fraction of fluorophores was emitting at any given instant. A cylindrical lens was put into the imaging path to introduce astigmatism to encode the depth (z) position into the ellipticity of the single-molecule images^33^. The EMCCD (iXon Ultra 897, Andor) camera recorded images at a 110-frame-rate for a frame size of 256 × 256 pixels and typically recorded ≈50000 frames for each experiment. In addition, to form the 2D-STORM imaging, we removed the cylindrical lens in the optical layout.

#### STORM reconstruction

The open-source software package Thunder-STORM^31^ and customized 3D-STORM software^33^ were used for STORM image reconstruction. Images labeled ‘ME-MLE’ and ‘SE-MLE’ were reconstructed by Thunder-STORM with maximum likelihood estimation (integrated PSF method), and multi-emitter fitting enabled (‘ME-MLE’) or not (‘SE-MLE’). The images labeled ‘SE-Gaussian’ were reconstructed with the customized 3D-STORM software by fitting local maxima with an (elliptical) Gaussian function described previously in ref^33^. Drift correction was performed post-localization, and images were rendered using a normalized Gaussian function (σ as 2 pixels).

#### Cell culture, fixation, and immunofluorescence

COS-7 cells were cultured in DMEM (GIBCO, 21063029) supplemented with 10% fetal bovine serum (FBS; GIBCO) in a humidified CO2 incubator with 5% CO2 at 37 °C, following standard tissue-culture protocols. Then, cells were seeded on 12 mm glass coverslips in a 24-well plate at ~2 × 10^4^ cells per well and cultured for 12 h. For STORM of actin filaments, a previously established fixation protocol^57^ was employed: The samples were first fixed and extracted for 1 min with 0.3% v/v glutaraldehyde and 0.25% v/v Triton X-100 in cytoskeleton buffer (CB, 10 × 10^-3^ M MES, pH 6.1, 150 × 10^-3^ m NaCl, 5 × 10^-3^ m EGTA, 5 × 10^-3^ m glucose, and 5 × 10^-3^ m MgCl_2_), postfixed for 15 min in 2% (v/v) glutaraldehyde in CB, and reduced with a freshly prepared 0.1% sodium borohydride solution in PBS. Alexa Fluor 647-conjugated phalloidin was applied at a concentration of ≈0.4 × 10^-6^ m for 1 h. The sample was briefly washed two to three times with PBS and then immediately mounted for imaging. For the imaging of other targets, samples were fixed with 3% w/v paraformaldehyde and 0.1% w/v glutaraldehyde in PBS for 20 min. After reduction to a freshly prepared 0.1% sodium borohydride solution in PBS for 5 min, the samples were permeabilized and blocked in blocking buffer (3% w/v BSA, 0.5% v/v Triton X-100 in PBS) for 20 min. Afterward, the cells were incubated with the primary antibody (described above) in a blocking buffer for 1 h. After washing in a washing buffer (0.2% w/v BSA and 0.1% v/v Triton X-100 in PBS) three times, the cells were incubated with the secondary antibody for 1 h at room temperature. Then, the samples were washed three times with the washing buffer before being mounted for imaging.

### SIM imaging

#### TIRF-SIM

Our SIM system was built upon a commercial inverted fluorescence microscope (IX83, Olympus) equipped with a TIRF objective (100×/1.7 NA, Apo N, HI Oil, Olympus) and a multiband dichroic mirror (DM, ZT405/488/561/640-phase R; Chroma) as described previously^36^. In short, laser light with wavelengths of 488 nm (Sapphire 488LP-200) and 561 nm (Sapphire 561LP-200, Coherent) and acoustic, optical tunable filters (AOTFs, AA Opto-Electronic, France) were used to combine, switch, and adjust the illumination power of the lasers. A collimating lens (focal length: 10 mm, Lightpath) was used to couple the lasers to a polarization-maintaining single-mode fiber (QPMJ-3AF3S, Oz Optics). The output lasers were then collimated by an objective lens (CFI Plan Apochromat Lambda 2× NA 0.10, Nikon) and diffracted by a pure phase grating that consisted of a polarizing beam splitter, a half-wave plate, and an SLM (3DM-SXGA, ForthDD). The diffraction beams were then focused by another achromatic lens (AC508-250, Thorlabs) onto the intermediate pupil plane, where a carefully designed stop mask was placed to block the zero-order beam and other stray light and to permit passage of ±1 ordered beam pairs only. To maximally modulate the illumination pattern while eliminating the switching time between different excitation polarizations, a homemade polarization rotator was placed after the stop mask. Next, the light passed through another lens (AC254-125, Thorlabs) and a tube lens (ITL200, Thorlabs) to be focused onto the back focal plane of the objective lens, interfering with the image plane after passing through the objective lens. Emitted fluorescence collected by the same objective passed through a dichroic mirror, an emission filter, and another tube lens. Finally, the emitted fluorescence was split by an image splitter (W-VIEW GEMINI, Hamamatsu, Japan) before being captured by a sCMOS (Flash 4.0 V3, Hamamatsu, Japan) camera.

#### Hessian-SIM

We applied the Hessian denoising algorithm^36^ without the *t* continuity constraint on the Wiener-SIM reconstruction^37^ results to obtain the Hessian-SIM images, as shown in **Supplementary Fig. 18**.

#### Cell maintenance and preparation

Human umbilical vein endothelial cells (HUVECs) were isolated and cultured in an M199 medium (Thermo Fisher Scientific, 31100035) supplemented with fibroblast growth factor, heparin, and 20% FBS or in an endothelial cell medium (ECM) (ScienCell, 1001) containing endothelial cell growth supplement (ECGS) and 10% FBS. The cells were infected with a retrovirus system to express LifeAct-EGFP. The transfected cells were cultured for 24 h, detached using trypsin-EDTA, seeded onto poly-l-lysine-coated coverslips (H-LAF10L glass, reflection index: 1.788, thickness: 0.15 mm, customized), and cultured in an incubator at 37 °C with 5% CO2 for an additional 20–28 h before the experiments. Liver sinusoidal endothelial cells (LSECs) were isolated and plated onto 100 μg/ml collagen-coated coverslips and cultured in high-glucose DMEM supplemented with 10% FBS, 1% L-glutamine, 50 U/ml penicillin, and 50 μg/ml streptomycin in an incubator at 37 °C with 5% CO2 for 6 h before imaging. Live cells were incubated with DiI (100 μg/ml, Biotium, 60010) for 15 min at 37 °C, whereas fixed cells were fixed with 4% formaldehyde at room temperature for 15 min before labeling with DiI. For the SIM imaging experiments, cells were seeded onto coverslips (H-LAF 10L glass, reflection index: 1.788, diameter: 26 mm, thickness: 0.15 mm, customized).

### STED imaging

#### Microscope setup

Image acquisition of stimulated emission depletion (STED) microscopy^58^ was achieved using a gated STED (gSTED) microscope (Leica TCS SP8 STED 3X, Leica Microsystems, Germany) equipped with a wide-field objective (100×/1.40 NA, HCX PL APO, Oil, Leica). The excitation and depletion wavelengths were 647 nm and 775 nm, respectively. All images were obtained using the LAS AF software (Leica).

#### Cell maintenance and preparation

COS-7 cells were cultured in high-glucose DMEM (GIBCO, 21063029) supplemented with 10% fetal bovine serum (FBS, GIBCO) and 1% 100 mM sodium pyruvate solution (Sigma-Aldrich, S8636) in an incubator at 37°C with 5% CO^2^ until ~75% confluency was reached. To label the microtubules in live cells shown in **Supplementary Fig. 22**, COS-7 cells were incubated with SiR-Tubulin (Cytoskeleton, CY-SC002) for ~20 mins before imaging without washing.

### Open-source datasets

In addition to the custom-collected datasets, we also used freely available simulation/experiment datasets to illustrate the broad applicability of our method.

#### 2D-SMLM simulation datasets

The *‘Bundled Tubes High Density’* (361 frames) and *‘Bundled Tubes Long Sequence’* (12000 frames) datasets from the *‘Localization Microscopy Challenge datasets’*^11^ on the EPFL website were used as the high-density and low-density 2D-SMLM simulation datasets in this work, as shown in **Fig. 1c**. The NA of the optical system was 1.4 (oil-immersion objective), and the wavelength of the fluorescence was 723 nm.

#### 3D-SMLM

The ‘*MT1.N1LD*’ (19996 frames, 3D-Astigmatism PSF) dataset from the *‘Localization Microscopy Challenge datasets*’^32^ on the EPFL website was used as the low-density 3D-SMLM simulation dataset in this work, as shown in **Supplementary Fig. 6**. The NA of the optical system was 1.49 (oil-immersion objective), and the wavelength of the fluorescence was 660 nm. All the images had a frame size of 64 × 64 pixels (pixel size as 100 nm). Then, 20 frames from this low-density dataset were averaged into one frame to generate the corresponding high-density 3D-SMLM dataset (resulting in 998 frames).

#### 2D-SMLM experimental datasets

The *‘Localization Microscopy Challenge datasets’*^11^ also contain experimental data, and 500 high-density images of tubulins were acquired from the EPFL website (**Supplementary Fig. 7a**, **7b**). The NA of the optical system was 1.3 (oil-immersion objective), and the wavelength of the fluorescence was 690 nm. The images were recorded with a camera at a 25-frame-rate for a frame size of 64 × 64 pixels (pixel size as 100 nm).

#### Live-cell SRRF datasets

The GFP-tagged microtubules in live HeLa cells were imaged by the TIRF mode with a TIRF objective (100×/1.46 NA, Plan Apochromat, Oil, Zeiss) and an additional 1.6× magnification with 488 nm laser illumination^20^ (200 frames in total). The open-source ImageJ plugin^20^ was used to reconstruct the SRRF results (**Supplementary Fig. 7c**).

### Simulations of the grid imaged by SMLM

Following ref^19^, we created a regular grid on a pixel of 10 nm in size (**Supplementary Fig. 5c**). The density of the randomly activated molecule was set as increasing gradually from the center to the sides. Then, the resulting image sequence was convoluted with a Gaussian kernel with an FWHM of 280 nm and down-sampled ten times (pixel size 100 nm). After that, Poisson and 20% Gaussian noise were injected into the image sequence (**Supplementary Fig. 5d**). Finally, the image sequence was reconstructed by Thunder-STORM with maximum likelihood estimation (integrated PSF method), which enabled the multi-emitter fitting function.

### Simulation of Fourier ptychographic microscopy (FPM)

We used the United States Air Force (USAF) resolution target as the ground-truth sample of the FPM^59^ (**Supplementary Fig. 21a**). The intensity and phase of the imaged sample were both set as those of the USAF target with a size of 240 × 240 pixels (pixel size: 406.3 nm). Illumination from different angles was provided by a 7 × 7 LED matrix, whose emission wavelength was 532 nm and distance to the sample was 90 mm. The sample was illuminated by each LED unit, filtered by the objective (4×/0.1 NA), and sampled by the camera (image size as 60 × 60 and pixel size as 1.625 μm). After the LEDs illuminated the sample, the final 49 low-resolution images were obtained. We used the image illuminated by the LED in the center as the initial image. Then, the amplitude and phase of the corresponding aperture were updated in turn in each FPM iteration. After 10 iterations, the final high-resolution complex-amplitude image (240 × 240) was obtained, the size of which was enlarged by 4× compared to the corresponding low-resolution images.

### Data generation processes of learning-based applications

#### The 240 nm PSF to 120 nm PSF image transformation

The deep neural network (DNN) was trained with 240 nm PSF convoluted images and corresponding 120 nm PSF convoluted images as the ground truth. We created synthetic tubulin structures using the random walk process to simulate two-dimensional trajectories with randomly changing orientations, and the maximal curvature was set as a limited value, respecting the known physical stiffness properties of tubulin^40^. The structures were then convoluted with a 240 nm PSF or a 120 nm PSF, and down-sampled 2 times (40 nm pixel size) as the input or ground-truth. To simulate the realistic fluorescent background, we convoluted the blurred images with a larger Gaussian kernel (FWHM as ~2.5 μm), and added it to the blurred images. Then, the Poisson noise and 10% Gaussian noise were involved in the images to produce the final input images for DNN training. Following the same procedure, additional 24 images were generated as test dataset.

#### Sparse sampling

The DNN was trained with sparsely sampled geometrical structures and corresponding intact structures as the ground truth. We chose four simple and common geometrical structures, i.e., triangles, circles, rectangles, and squares, for the simulations^60^. The spatial size and the number of structures in one input image are shown in **Supplementary Table 1**. After obtaining the structures, we randomly sampled the image at a sampling rate of 8%. We selected rectangular structures and used 5000 images as the training dataset. For each geometrical structure, we generated 200 images as a test dataset.

#### Noise2Noise

Noise2Noise^46^ is an unsupervised learning procedure to denoise noisy images without clean ones. The DNN only looks at noisy image pairs (two images with independent noise that share the exact details) during training, i.e., one as input and the other as the output target. The fluorescence microscopy denoising (FMD) dataset^47^ was used in this Noise2Noise task. We chose fixed zebrafish embryos [EGFP-labeled *Tg (sox10:megfp)* zebrafish at 2 days postfertilization] as the dataset, imaged by a commercial Nikon A1R-MP laser scanning confocal microscope at very low excitation power. This imaging configuration has 5 noise levels. The raw images had the highest noise level, and images at other noise levels were generated by averaging multiple frames (2, 4, 8, and 16) of raw images using the circular averaging method. To test extreme conditions, we chose only the raw images with the highest noise level as the input of the training set (every two raw images). For each FOV (total of 20) with 50 different noise realizations, we randomly chose 200 noise-noise data pairs. Moreover, we cropped the raw images of size 512 × 512 to four nonoverlapping patches of size 256 × 256. Finally, we obtained 20 × 200 × 4 = 16000 images as the training dataset. By averaging 50 noisy raw images, we generated the ground-truth reference to evaluate the accuracy of the Noise2Noise prediction.

### Network architecture and training procedure

#### Network architecture

The network architecture, called the U-shaped architecture (U-net), is composed of a contracting path and an expansive path^61^. In the contracting path, the input layer is followed by a successive down-convolution block, consisting of 4 × 4 kernel convolution with a stride step of 2, batch normalization (BN)^62^, and a leaky rectified linear unit (LeakyReLU) function. A convolutional layer lies at the bottom of this U-shaped structure that connects the down-convolution and up-convolution blocks. The expansive pathway combines the feature and spatial information from the contracting path through a series of up-convolution blocks (Upsampling2D operation + 4 × 4 kernel convolution with stride step of 1 + BN + ReLU) and concatenations with high-resolution features. The last layer is another convolutional layer that maps the 32 channels into one channel image. We used two U-shaped network architectures (U-net^1^ and U-net^2^) in different tasks (**Supplementary Fig. 13**). U-net^1^ has 7 down-convolution blocks and 7 up-convolution blocks, whereas U-net^2^ has 4 down-convolution blocks and 4 up-convolution blocks.

#### Training procedure

All the networks were trained using stochastic gradient descent with adaptive moment estimation (Adam)^63^. The detailed input patch size of the training images, number of epochs, batch size, number of training images, learning rate, network architecture, number of parameters, and loss function for each task were shown in **Supplementary Fig. 14** and **Supplementary Table 2**. All the training procedures were performed on a local workstation equipped with an NVIDIA Titan Xp GPU card. The related learning framework was implemented with the TensorFlow^64^ framework (version 1.8.0) and Python (version 3.6).

### Using open-source deep-learning models

#### ANNA-PALM

ANNA-PALM^45^ computationally reconstructs SR images from sparse, rapidly captured localization data. ANNA-PALM was trained using densely sampled PALM images (long sequence) as the ground truth and the corresponding sparsely sampled PALM images (short sequence) as the input. ANNA-PALM is based on a conditional GAN^65^ (cGAN^1^ in **Supplementary Fig. 14**) with U-net^1^ as the generator. We tested the performance of ANNA-PALM using the 500 high-density images of tubulins from the EPFL website^11^. The fluorophores in frames 1-25 and 26-50 were localized using the ME-MLE estimator to construct two sparse SR inputs, and then the trained ANNA-PALM model predicted the corresponding dense sampled images.

#### CARE

The CARE framework has been described in detail elsewhere^40^; it is a computational approach that can extend the spatial resolution of microscopes using the U-net^3^ architecture. We fed the open-source trained model of CARE with the two averaged images (100 frames for each) from the *open-source SRRF dataset*, generating the corresponding two super-resolved images.

#### Cross-modality super-resolution

The cross-modality imaging ability of the DNN was demonstrated previously^66^ by mapping the TIRF to the TIRF-SIM modality (TIRF2SIM) using the cGAN approach. The cGAN^2^ (**Supplementary Fig. 14**) in TIRF2SIM is based on U-net^2^ with the residual convolutional blocks (Res-Unet) as the generator. It was trained and tested using AP2-eGFP-tagged clathrin in gene-edited SUM159 cells. We used the provided ImageJ plugin and example data to reproduce the results directly (**Supplementary Fig. 27b**).

#### DFGAN-SIM

The deep Fourier channel attention network trained with the GAN strategy (DFGAN-SIM)^67^ was developed to reconstruct SIM images by inputting 9 raw frames. In **Supplementary Fig. 27g**, we employed the provided frozen network weights (‘DFGAN-SIM_MTs’ trained on microtubule images, with enconsin-mEmerald in the COS-7 cells) and their example SIM data (microtubule images in the BioSR dataset, with enconsin-mEmerald in the COS-7 cells) to reproduce the results directly. We used the images captured by our SIM system (mitochondrial cristae images, with MitoTracker Green in the COS-7 cells) to test the same model (‘DFGAN-SIM_MTs’), as shown in **Supplementary Fig. 27l**.

### Image rendering and processing

We used the custom-developed color maps, shifted Jet and black Jet (sJet and bJet), to visualize the rFRC maps in this work. The color maps ‘SQUIRREL-FRC’^19^ were used to present the FRC maps in the **Supplementary Fig. 15d**, and **17**. The color maps ‘SQUIRREL-Errors’^19^ were used to present the difference map in the second and third columns of **Fig. 1b**, **1c**, the bottom panel of **Fig. 2h**, the fourth column of **Fig. 4e**, the bottom panel of **Fig. 4g**, and the third panel of **Supplementary Fig. 7b**. The volumes in **Supplementary Fig. 9a-9c** were rendered by ClearVolume^68^. The Jet color map projection was used to show the intensity in **Supplementary Fig. 2c**, and the depth in **Supplementary Fig. 9g**. All data processing was achieved using MATLAB and ImageJ. All figures were prepared with MATLAB, ImageJ, Microsoft Visio, and OriginPro.

## Data availability

All the data that support the findings of this study are available from the corresponding author on request.

## Code availability

The updated version of this work in MATLAB library can be found at https://github.com/WeisongZhao/PANELM, and the corresponding Python library can be found at https://github.com/WeisongZhao/PANELpy. The updated ImageJ plugin and its source code can be found at https://github.com/WeisongZhao/PANELJ. The online tutorials can be found on the corresponding GitHub wikis.

## Acknowledgments

We thank National Center for Protein Sciences at Peking University in Beijing, China, for assistance with STED imaging experiments. H. L. acknowledges support by grants from the National Natural Science Foundation of China (61805057); the Young Elite Scientists Sponsorship Program (2018QNRC001); and the Natural Science Foundation of Heilongjiang Province (YQ2021F013). L. C. acknowledges support from grants from the National Natural Science Foundation of China (92054301, 81925022, 31821091, 91750203), the National Science and Technology Major Project Program (2016YFA0500400); and the Beijing Natural Science Foundation (Z20J00059). P. L. acknowledges support from grants from the National Natural Science Foundation of China (11874231); and Guangdong Major Project of Basic and Applied Basic Research (2020B0301030009). H. L. and W. Z. acknowledge support from the State Key Laboratory of Robotics and Systems. L. C. acknowledges support from the High-performance Computing Platform of Peking University. S. Z. acknowledges support from the Boya Postdoctoral Fellowship of Peking University. H. M. acknowledges support from grants from the National Natural Science Foundation of China (32071458).

## Author contributions

H. L., L. C., L. P., and W. Z. supervised the project; W. Z., H. L., and L. C. initiated and conceived the research; W. Z. developed the method; W. Z. implemented the corresponding software with the contribution of L. Q. and G. Q.; W. Z. designed the theoretical model and experiments, analyzed the data, and prepared the figures; X. H. and J. Y performed the experiments and collected the data with the contribution of S. Z.; W. Z. performed the simulations and tests of the learning-based applications with the contributions of G. Q., L. Q., Y. Z., and X. W.; Z. L. reproduced and tested the DFGAN-SIM under the supervision of H. M.; Y J., H. M., X. D., J. T., Y. H., and L. P. participated in discussions during the development of the manuscript; W. Z., H. L., and L. C. wrote and revised the manuscript with input from all authors; All authors participated in the discussions and data interpretation.

## Competing interests

L. C., H. L., and W. Z. have a pending patent application on the presented framework.

## Notes

https://github.com/WeisongZhao/PANELM

https://github.com/WeisongZhao/PANELpy

https://github.com/WeisongZhao/PANELJ

