## Supplementary Information for "Quantitatively mapping local quality of super-resolution microscopy by rolling Fourier ring correlation"

### Content

**Supplementary Figures.**---------------------------------------------------------------------------------------------------------------- **3**

**Supplementary Notes.** ---------------------------------------------------------------------------------------------------------------**17**

Supplementary Note 1 | The stability and resolvability of rFRC map. ---------------------------------------------------------------17

Supplementary Note 2 | SIM applications. ------------------------------------------------------------------------------------------21

Supplementary Note 3 | Deconvolution applications.--------------------------------------------------------------------------------23

Supplementary Note 4 | FPM applications. ------------------------------------------------------------------------------------------26

Supplementary Note 5 | STED applications.-----------------------------------------------------------------------------------------27

Supplementary Note 6 | Comparisons of rFRC, RSM, and SSIM.--------------------------------------------------------------------28

Supplementary Note 7 | The uncertainties in learning-based applications. -----------------------------------------------------30

Supplementary Note 8 | Single-frame strategies for rFRC mapping. ------------------------------------------------------------35

Supplementary Note 9 | Limitations.----------------------------------------------------------------------------------------------------41

**Supplementary Tables.** -----------------------------------------------------------------------------------------------------------**44**

Supplementary Table 1 | Parameters of geometrical structure simulations.-----------------------------------------------------44

Supplementary Table 2 | Details on network architecture, loss function, and training procedure.-----------------------------45

**References.**--------------------------------------------------------------------------------------------------------------------------**46**

#### Supplementary Figures.

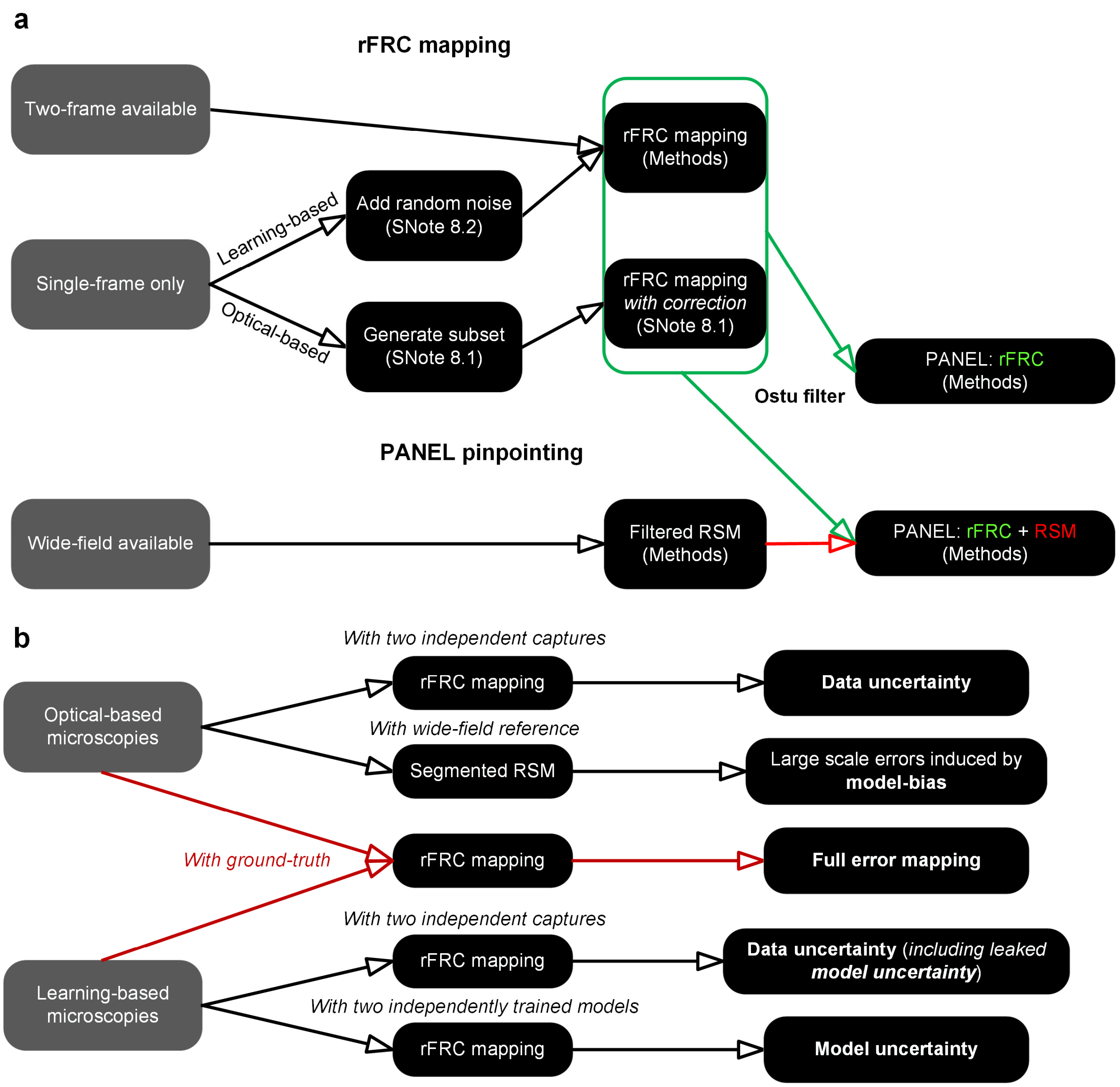


**Supplementary Fig. 1 | Abstract workflow.** (**a**) Abstract workflow. Only when the corresponding tasks satisfy two conditions, i.e., (i) belonging to 2D data and (ii) the existence of a wide-field reference, will the RSM be included in the PANEL visualization. (**b**) Our framework for estimating different types of uncertainties. At the SR scale, our method is capable of mapping (i) data uncertainty of image reconstructions without referencing the ground-truth (*Reconstruction-1* vs. *Reconstruction-2*); (ii) data uncertainty with leaked model uncertainty for deep-learning predictions without ground-truth (*Prediction-1* vs. *Prediction-2*); (iii) model uncertainty of deep-learning predictions without ground-truth (*Prediction-1* from *Model-1* vs. *Prediction-2* from *Model-2*); and (iv) full error of reconstructions/predictions with ground-truth (*Reconstruction*/*Prediction* vs. *Ground-truth*).

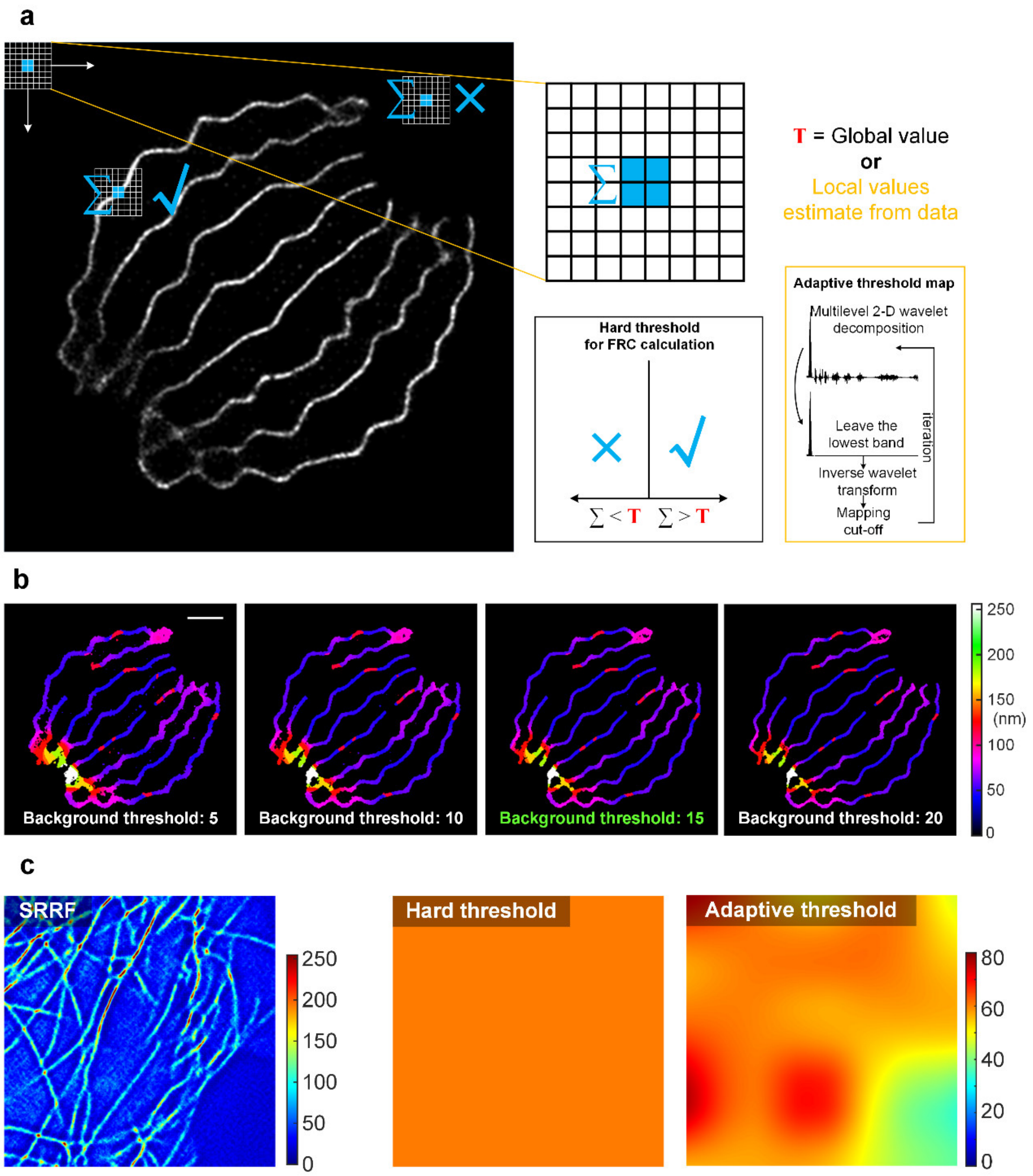


**Supplementary Fig. 2 | Two background skip strategies for rFRC mapping.** (**a**) Workflows (*c.f.*, **Fig. 1c**) of the background thresholding methods. During the rolling operation of the rFRC mapping, the intensity of center pixels from each block is summed (blue summation sign). The FRC value is calculated and assigned only if this summed value of the center pixels is larger than (blue tick sign) the threshold ('$\sum > \mathbf{T}$'); otherwise, the center pixel is set to zero (blue cross sign) ('$\sum < \mathbf{T}$'). In this work, we provided two strategies for threshold determination. One is the user-defined hard threshold for the entire image ('15' as in this representative example). The other is the iterative wavelet transform method (yellow box), which automatically estimates the local threshold values. (**b**) rFRC maps using different background thresholds (*c.f.*, **Fig. 1c**). (**c**) A representative SRRF data (left) (*c.f.*, **Supplementary Fig. 7c**) for illustration of two strategies of background thresholding (middle for hard threshold and right for adaptive threshold).

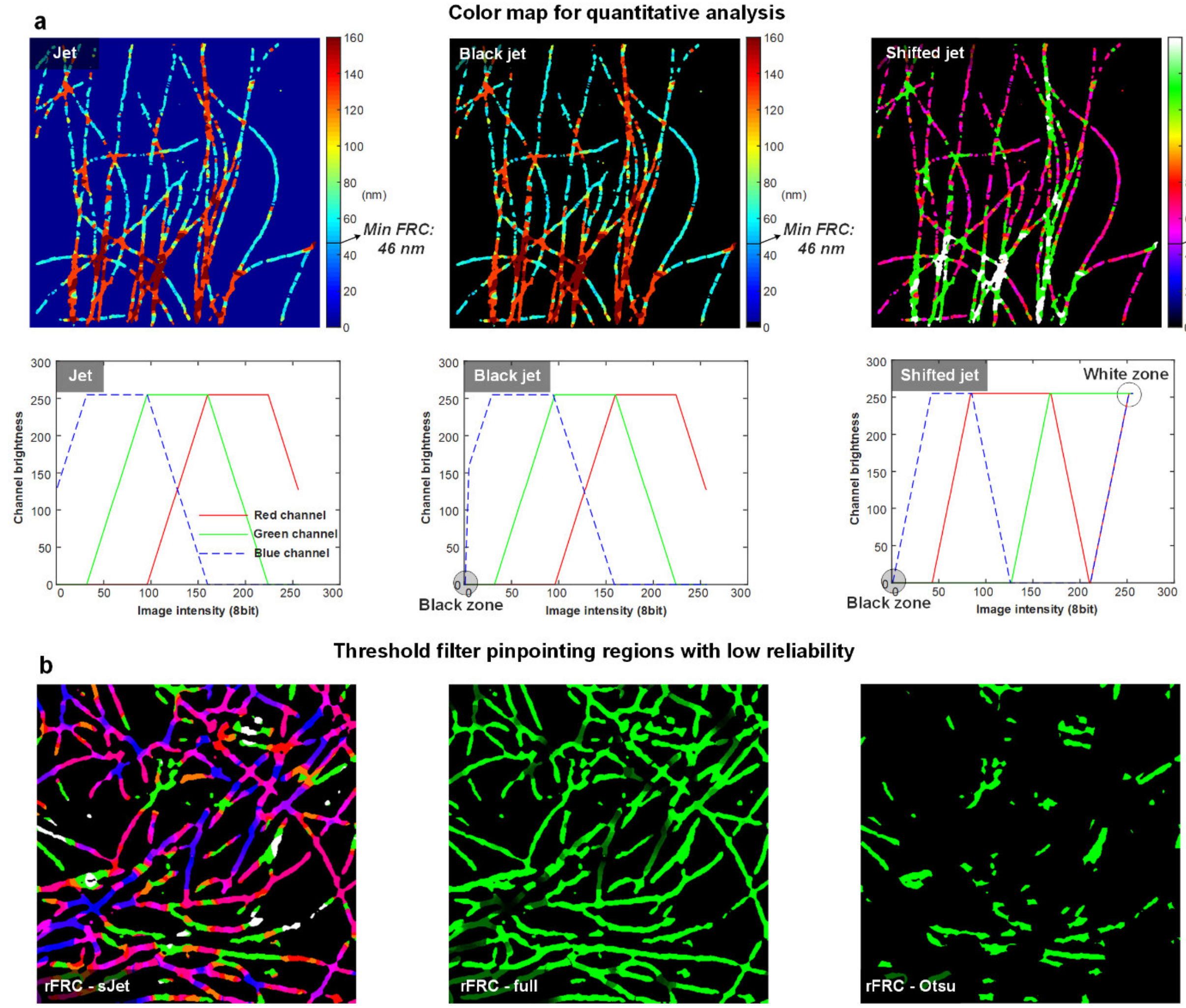


**Supplementary Fig. 3 | Color maps for map display and Otsu threshold for PANEL pinpointing.** (**a**) The representative color-coded images and color indexes of jet (left), black jet (middle), and shifted jet (right) color maps. The image is adapted from **Supplementary Fig. 7a**. (**b**) Otsu threshold for PANEL highlighting. Left: The rFRC map of the SRRF dataset in **Supplementary Fig. 7c**, displayed in the sJet color map. Middle and right: The Full rFRC map (middle) and the rFRC map after the Otsu threshold (right), regions with low reliability in the SRRF reconstruction are pointed by green.

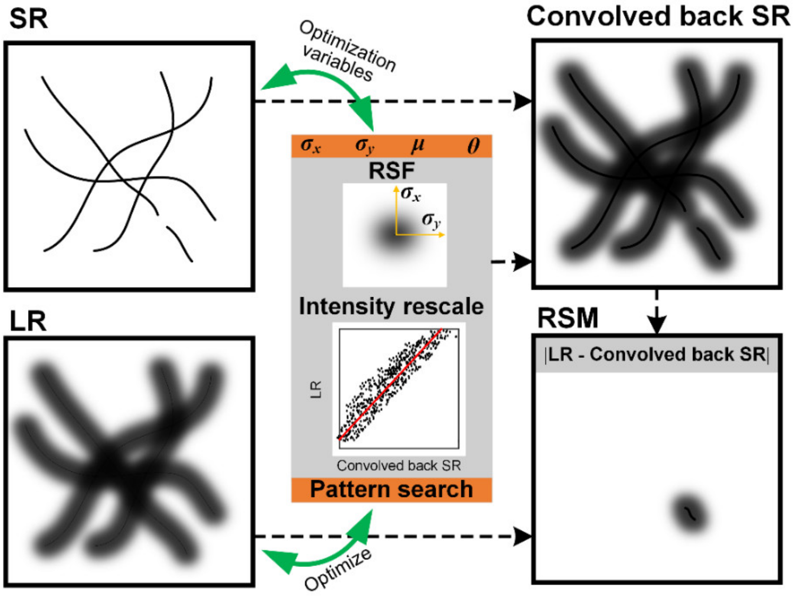


**Supplementary Fig. 4 | The workflow of the RSM.**

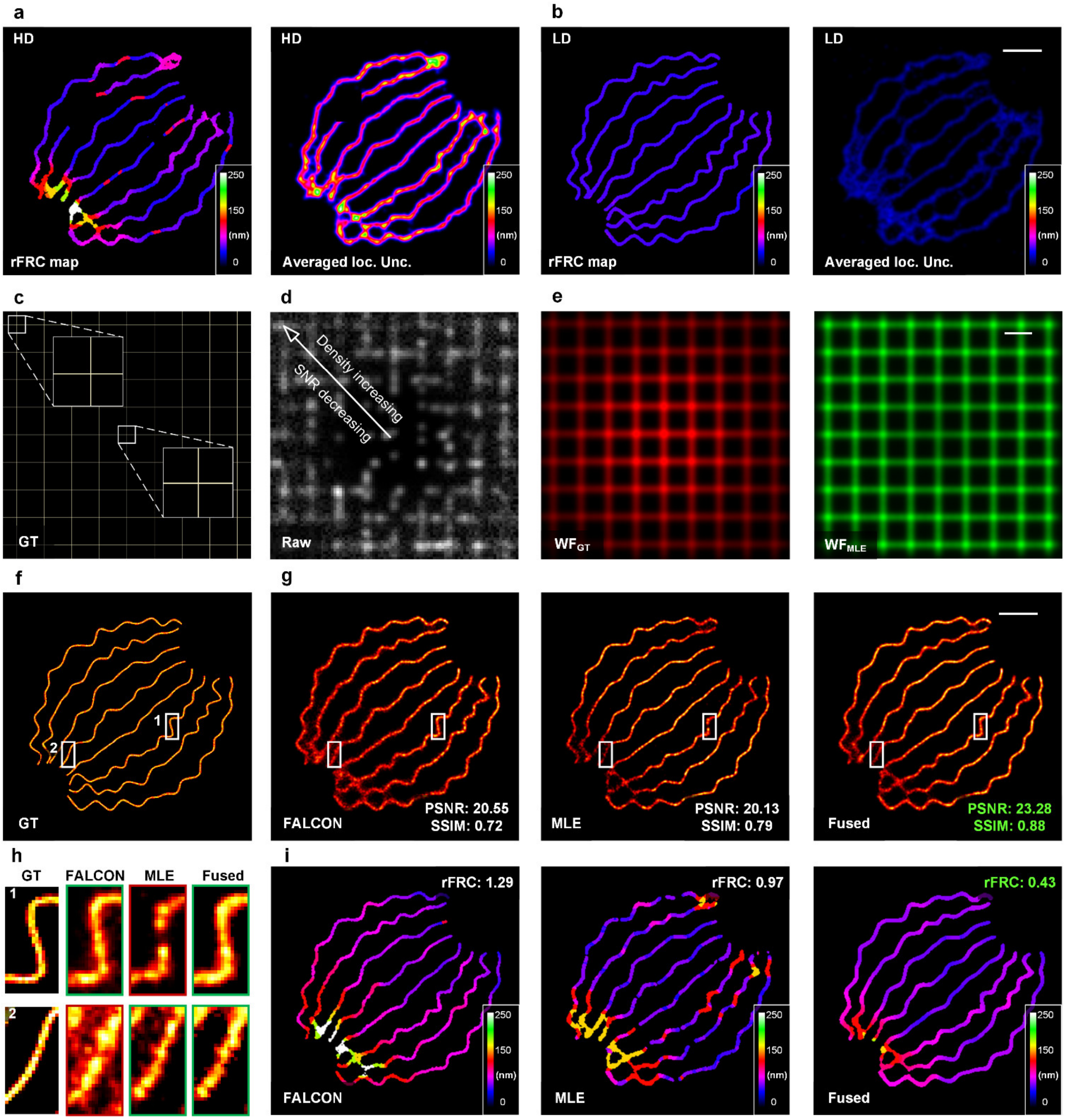


**Supplementary Fig. 5 | Full data of 2D-SMLM simulations and image fusion of SMLM data using rFRC map.** (**a**, **b**) The rFRC map versus the localization uncertainty of 2D-SMLM with high-density ('HD', **a**) and low-density ('LD', **b**) emitting fluorophores in each frame (*c.f.*, **Fig. 1c**). The overall resolution distribution of the rFRC map (left) is close to the averaged localization uncertainty map (right). For visualization, we provided the averaged localization uncertainty ('Averaged loc. unc.') map at the right, which is the raw localization uncertainty map filtered with the Gaussian function. (**c**, **d**) The ground-truth structures (**c**), and one representative raw frame (**d**) (*c.f.*, **Fig. 1d**). (**e**) The wide-field ground-truth image (WF$_{GT}$) and wide-field images generated from the MLE reconstruction (WF$_{MLE}$). (**f-i**) Image fusion of simulated 2D-SMLM data. (**f**) The ground-truth structures. (**g**) The reconstruction results of FALCON (left) and MLE (middle) algorithms, and the corresponding fused result of these two methods (right). The PSNR and SSIM values (reconstructions versus ground-truth) are labeled on the right bottom. (**h**) Enlarged regions enclosed by white boxes in (**f**) and (**g**). (**i**) rFRC maps of (**g**). Scale bars: (**b**, **g**) 500 nm; (**e**) 1 μm.

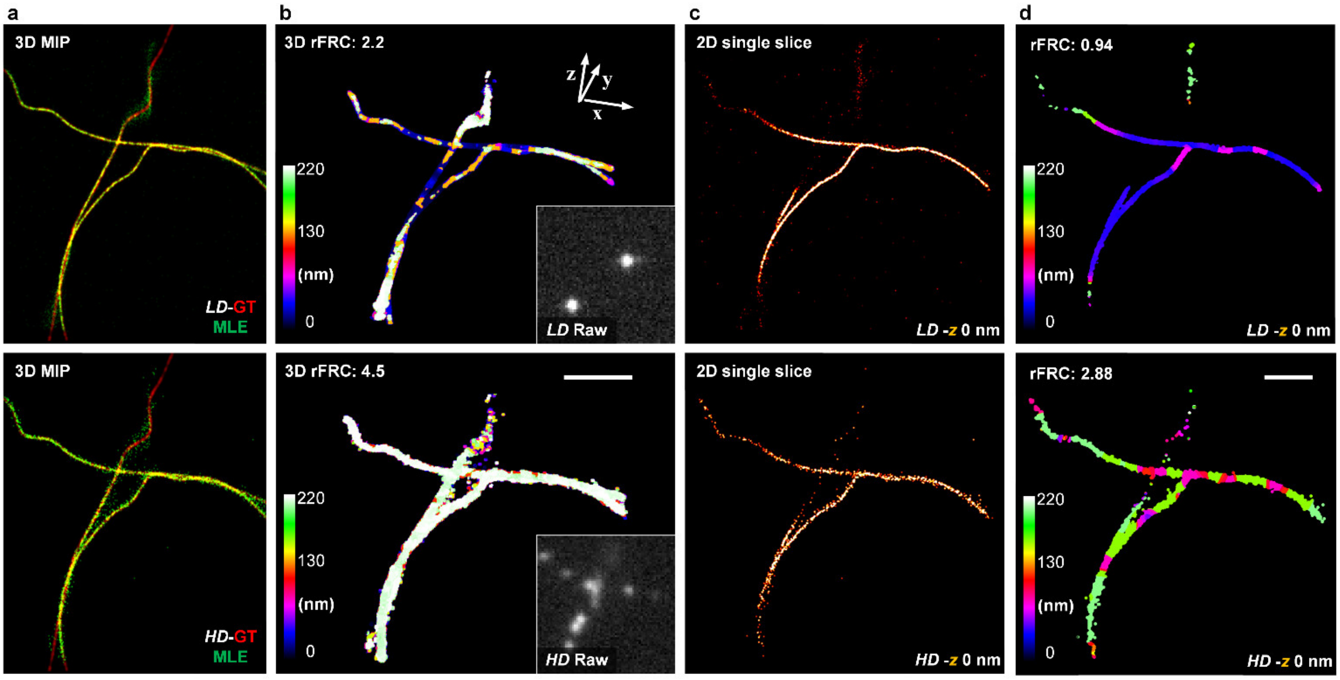


**Supplementary Fig. 6 | 3D-SMLM simulations evaluated by rFRC map.** (**a**) The merged maximum intensity projection (MIP) views of ground-truth structures (red channel, labeled as '*LD*-GT' (top) or '*HD*-GT' (bottom) for low-density or high-density emitting fluorophores), and the corresponding 3D-MLE reconstructions (green channel, labeled 'MLE'). (**b**) The rFRC maps of low-density (top) and high-density (bottom) 3D-MLE reconstructions. Insets show representative frames of low-density (top) and high-density (bottom) datasets. (**c**) Horizontal sections (at 0 nm *z* position) of 3D-MLE reconstructions (low-density at the top and high-density at the bottom). (**d**) rFRC maps of corresponding horizontal sections in (**b**). Scale bars: 1 μm.

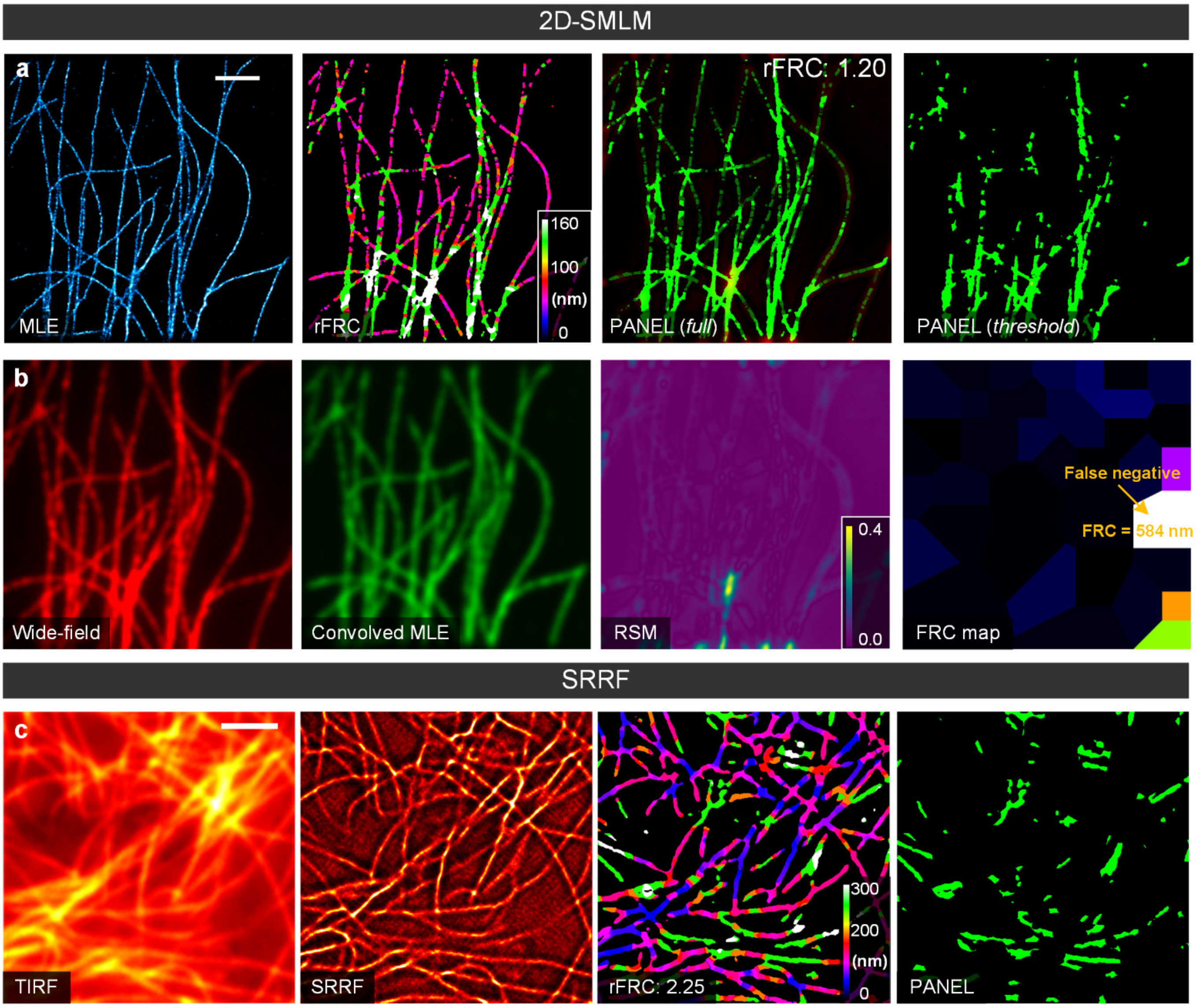


**Supplementary Fig. 7 | Open-source 2D-SMLM and SRRF experimental datasets evaluations.** (**a**) From left to right: MLE localization result of 500 high-density images of tubulins from the EPFL website (**Methods**); the rFRC map of the MLE; full merged RSM and rFRC map of the MLE; PANEL visualization. (**b**) From left to right: Corresponding wide-field image; MLE image convolved back to its original low-resolution scale; RSM of the MLE; FRC map of the MLE. (**c**) From left to right: Diffraction-limited TIRF image; SRRF reconstruction result of 100 fluctuation images (GFP-tagged microtubules in live HeLa cells, **Methods**); rFRC map of SRRF; PANEL visualization. Scale bar: 2 μm.

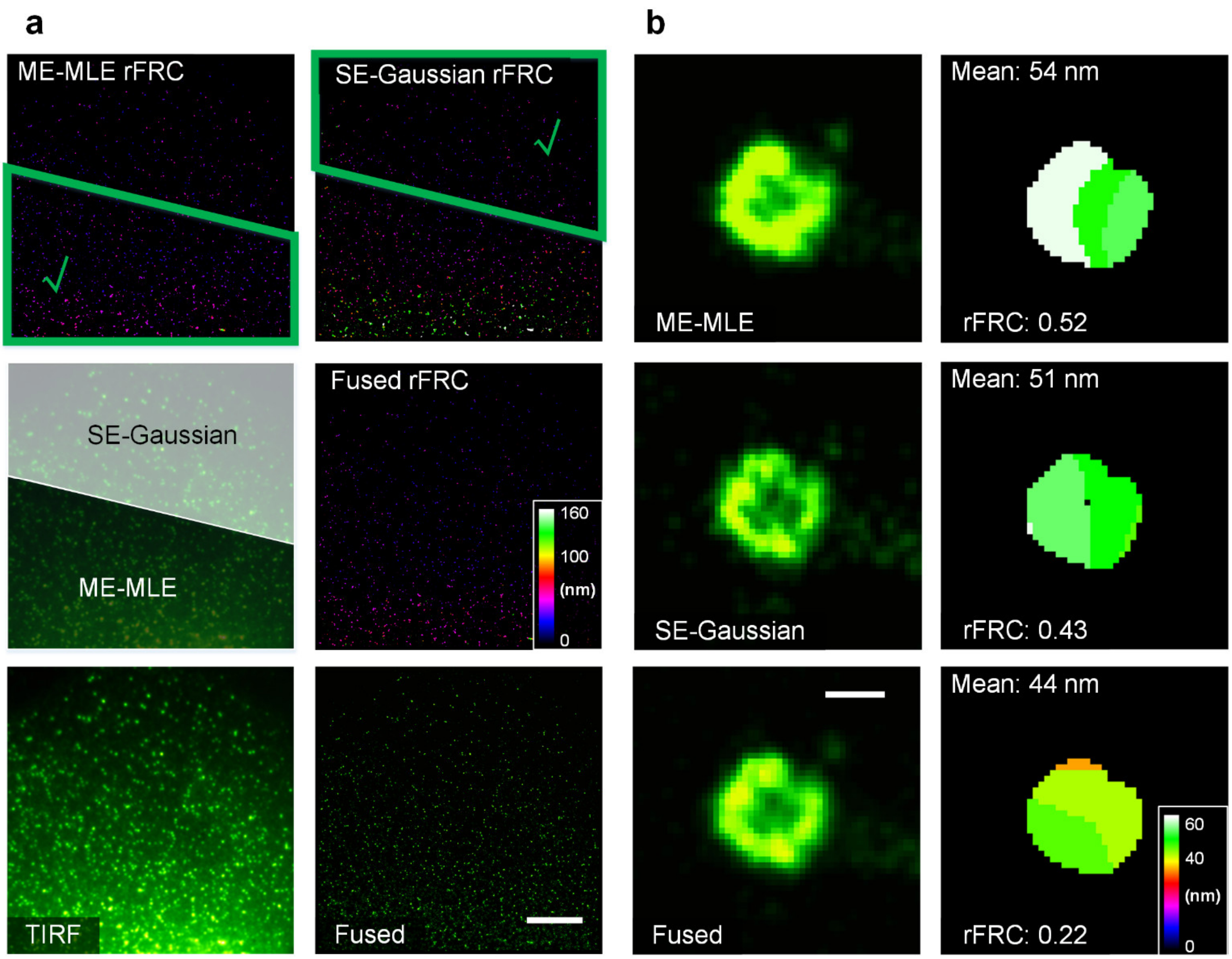


**Supplementary Fig. 8 | A representative example of STORM fusion (COS-7 cells, heavy chain clathrin-coated pits labeled with Alexa Fluor 647).** (**a**) The rFRC map of ME-MLE (top), the superiority map (middle) for fusion, and the TIRF image (bottom) are shown on the left. The rFRC maps of the SE-Gaussian (top) and fusion (middle) results, and the fusion result ('Fused', bottom) are displayed on the right. *We found that the ME-MLE method achieves superior performance in the regions containing a strong background and that the SE-Gaussian method obtains better reconstruction quality in the regions containing a weak background.* (**b**) Magnified results for a single CCP of ME-MLE (top), SE-Gaussian (middle), and fusion ('Fused,' bottom) are shown on the left, and the corresponding rFRC maps are demonstrated on the right. The mean resolutions are marked on the top left of the rFRC maps. In addition to the stable performance of fusion in the whole field of view, as highlighted in (**a**), the rFRC map assists in fusing fine structures such as a single ring-shaped CCP, enabling higher mean resolution. Scale bars: (**a**) 5 μm; (**b**) 100 nm.

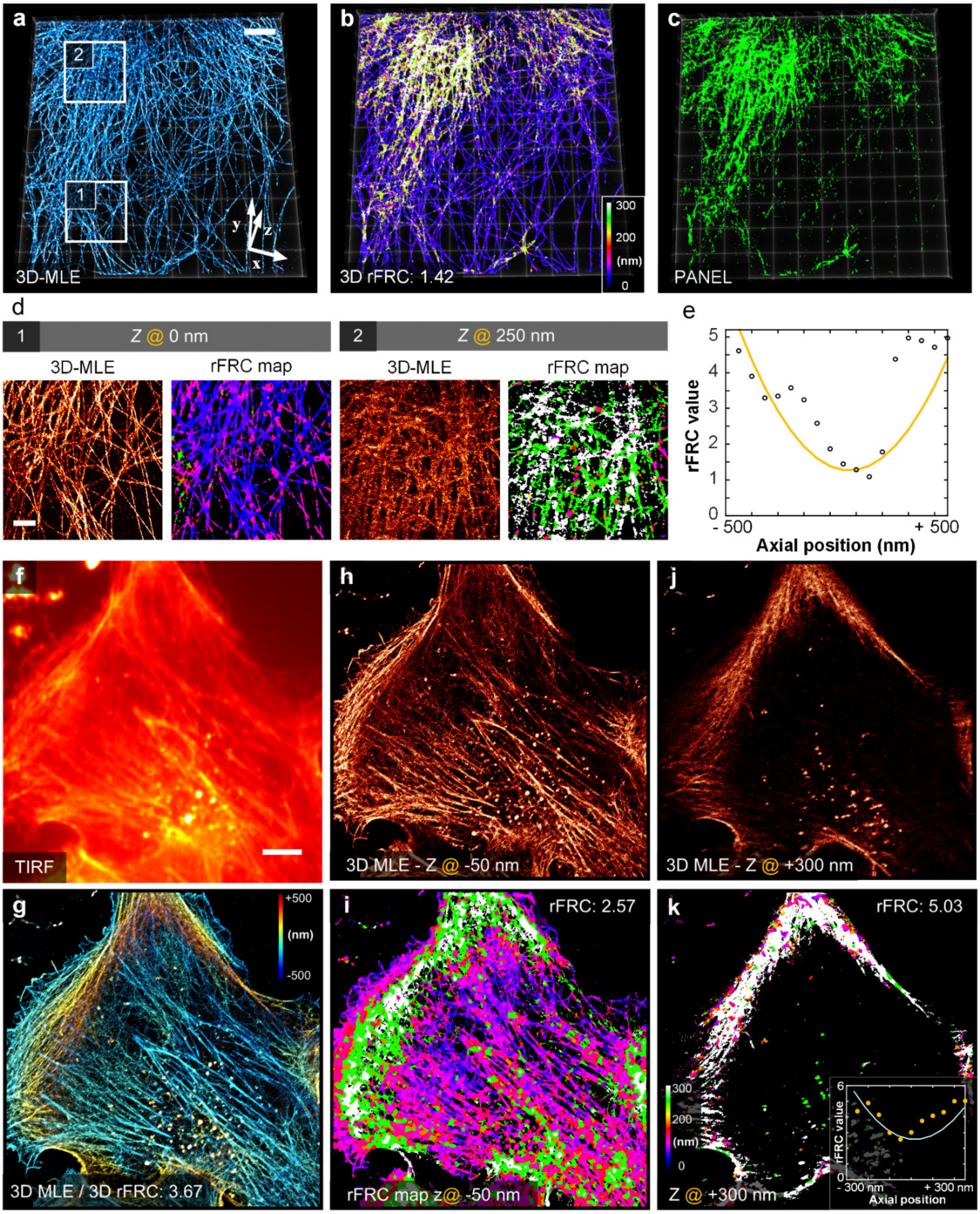


**Supplementary Fig. 9 | Evaluating 3D-STORM experiments.** (**a**) 3D-MLE reconstruction (COS-7 cells, α-tubulin labeled with Alexa Fluor 647). (**b**) 3D rFRC map of (**a**). (**c**) PANEL after the Otsu threshold of (**b**). (**d**) Corresponding magnified horizontal sections of the 3D-MLE (left) and rFRC (right) volume of the white boxes in (**a**). (**e**) The curve of the rFRC values along with the axial positions. (**f**) A maximum intensity projection (MIP) view of the TIRF (COS-7 cells, labeled with Alexa Fluor 647-phalloidin). (**g**) Depth color-coded view of 3D-MLE reconstruction. (**h**, **i**) Horizontal section of 3D-MLE reconstruction (**h**) at the -50 nm *z*-position and the corresponding rFRC map (**i**). (**j**, **k**) Horizontal section of 3D-MLE reconstruction (**j**) at the +300 nm *z*-position and the corresponding rFRC map (**k**). Scale bars: (**a**, **e**) 5 μm; (**d**) 2 μm.

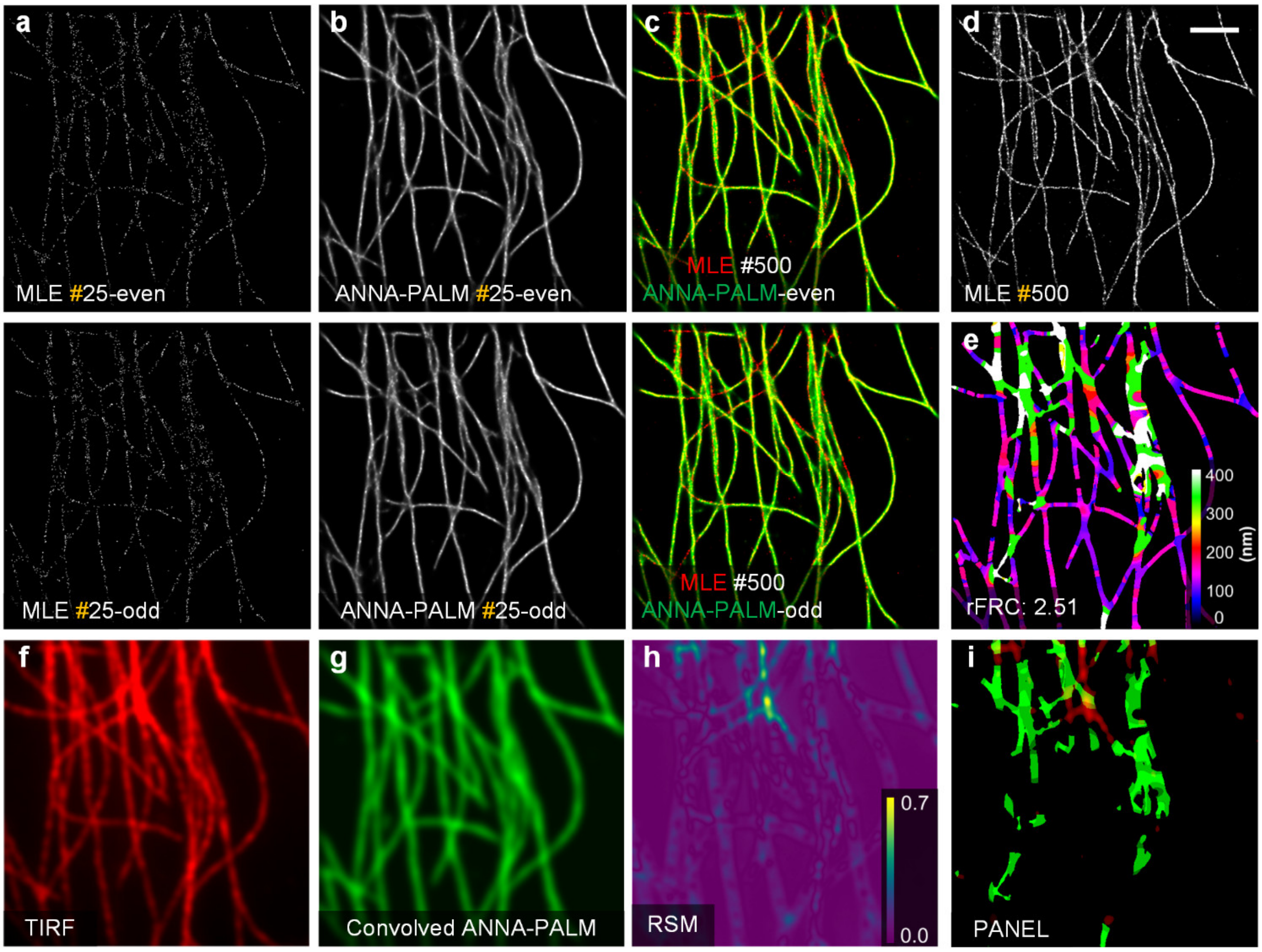


**Supplementary Fig. 10 | Full visualization of ANNA-PALM experiment (*c.f.*, Fig. 4a-4e).** (**a-c**) Sparse MLE reconstructions (**a**), ANNA-PALM results (**b**) from even frames (25 frames, top) and odd frames (25 frames, bottom), and the merged ANNA-PALM results (green channel) with full dense MLE reconstruction (red channel) (**c**). (**d**) Full MLE reconstruction. (**e**) The rFRC maps of sparse MLE reconstructions. (**f**) TIRF image. (**g**) TIRF image generated from ANNA-PALM reconstruction. (**h**) RSM. (**i**) PANEL visualization. Scale bar: 2 μm.

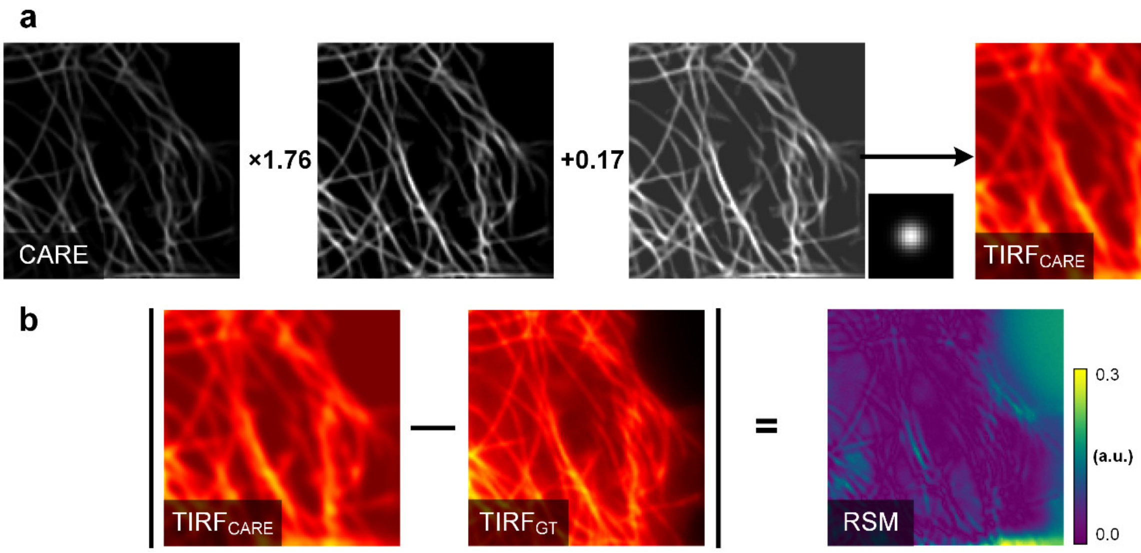


**Supplementary Fig. 11 | Visualizing the RSM estimation workflow of the CARE predicted data (*c.f.*, Fig. 4f-4i).** (**a**) The process of TIRF image generation from the CARE prediction, i.e., rescaled the intensity (CARE × 1.76 + 0.17) and convoluted the resulting image with the estimated resolution scaling function (RSF). (**b**) RSM generation, i.e., the absolute difference between the TIRF image generated from CARE prediction ($TIRF_{CARE}$) and the ground-truth TIRF ($TIRF_{GT}$) image.

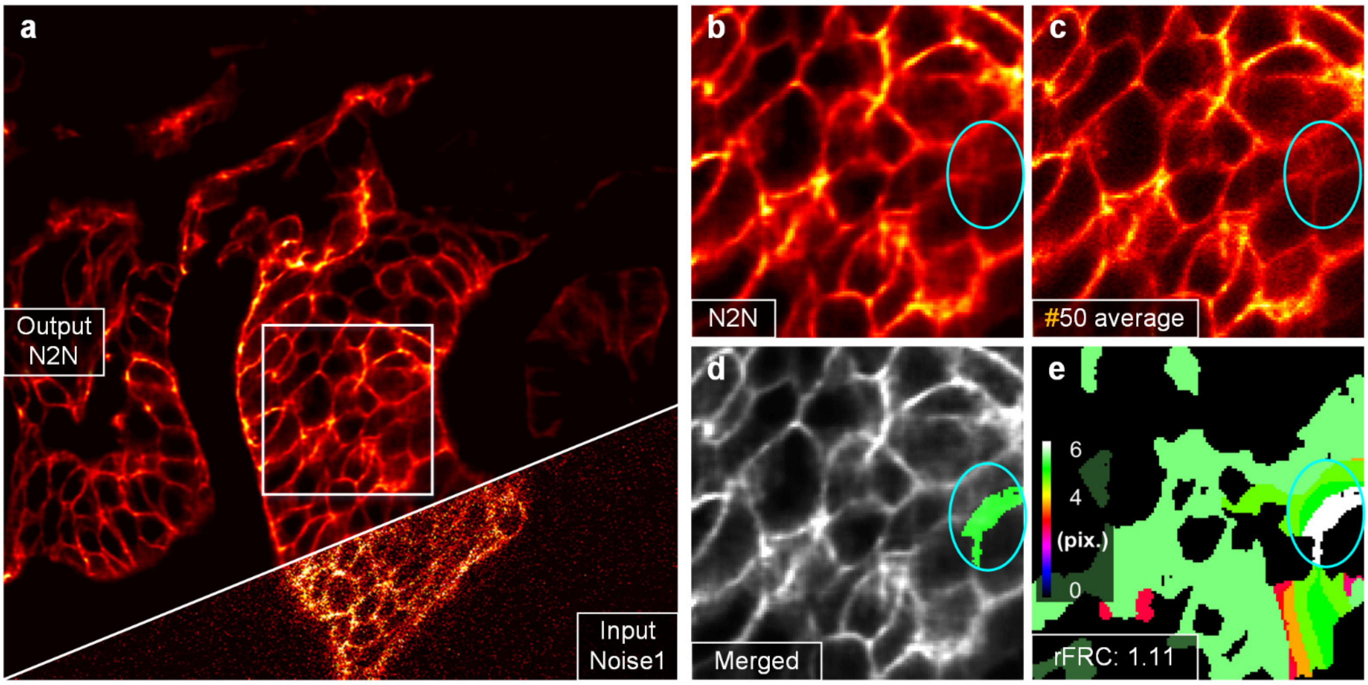


**Supplementary Fig. 12 | Full Noise2Noise experiment (*c.f.*, Fig. 4j-4m).** (**a**) Result (left top) after Noise2Noise ('N2N') and the noisy input (right bottom, 'Noise1'). (**b**) The Noise2Noise result from the white box in (**a**). (**c**) The reference image was obtained by averaging 50 noisy images with identical content. (**d**) Merged image of the PANEL (green channel) and Noise2Noise (gray channel) results. (**e**) rFRC map of (**b**). pix.: pixel.

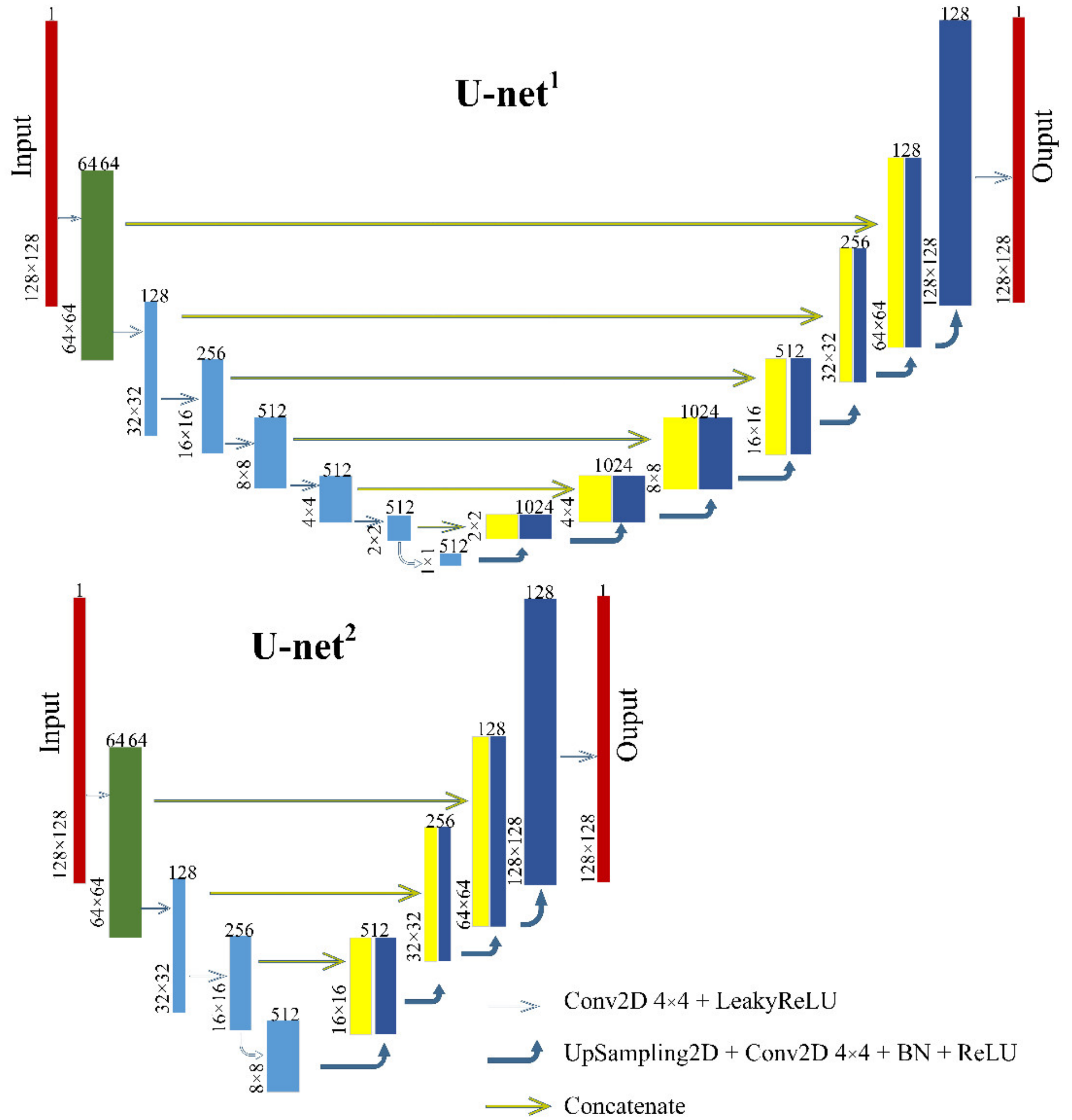


**Supplementary Fig. 13 | Neural network architectures used for different image restoration tasks.**

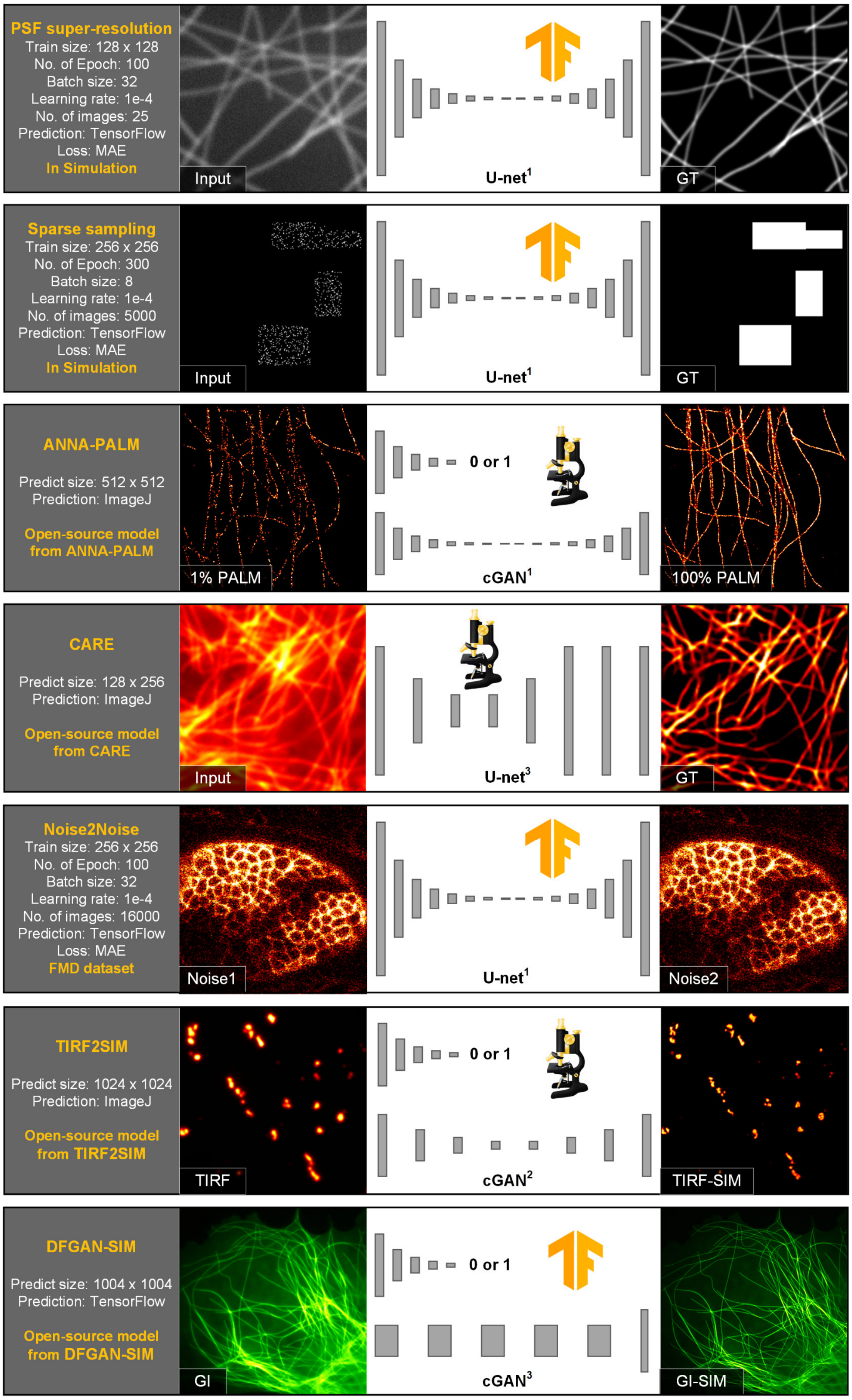


**Supplementary Fig. 14 | Overview of the network architecture, training parameter configuration, and data used for different applications.** From left to right: Task configuration; input image; topological structure of the network; and ground truth. The rows with ImageJ/Fiji icons ('microscope') indicate that we reconstructed the image using the published open-source models with the corresponding ImageJ/Fiji plugins. The rows with TensorFlow icons ('TF') indicate that we predicted the image using the corresponding model (trained by ourselves, except the last row 'DFGAN-SIM' employing the published open-source models) with the TensorFlow framework.

### Supplementary Notes.

#### Supplementary Note 1 | The stability and resolvability of rFRC map.

##### Supplementary Note 1.1 | The stability.

To avoid overconfident and unstable determination of the resolutions from small image blocks, we used the 3σ curve[1] as criterion in this work, i.e., three standard deviations above the expected random noise fluctuations as the threshold, instead of the popular-used 1/7 hard threshold[2-5]. This 3σ curve will adaptively change according to the input image block size, and thus it is more stable for local resolution estimation (**Supplementary Fig. 15a**). To test that, we created images with different block sizes (**Supplementary Fig. 15b**). In 1/7 hard threshold case, we found that the results were unstable at block size smaller than 256-pixel. Similarly, the 3σ curve from the 32-pixel block did not yield a stable resolution. However, with the larger block sizes (64-, 128, and 256-pixel), the 3σ criterion was stable and remained unchanged around the theoretical resolution. Furthermore, although a smaller block size (e.g., 32-pixel) may lead to more refined mapping, the overall distributions of these rFRC resolution maps (using different block sizes) are close to each other (**Supplementary Fig. 15c**). Therefore, to balance the mapping scale and its estimation stability, we chose a block size of 64-pixel as default in this work.

##### Supplementary Note 1.2 | The resolvability.

With a 64 × 64 block size window, overlapped image content may induce crosstalk on the resulting map. To test it and try the maximum resolvability, we simulated structures containing pairs of lines with gradually increasing spacing (**Supplementary Fig. 16**), and added different noise levels to these paired lines. After that, we applied the rFRC mapping on the resulting images and calculated the FRC resolution distributions of pixels on the left (yellow) and right lines (green) (**Supplementary Fig. 16**). In the 2-pixel case, the crosstalk between two lines is too significant for the rFRC mapping to distinguish the difference, and thus the overall distributions of FRC resolutions (yellow and green curves) are identical for different noise level. In the 4-pixel case, we observed the distributions of FRC are just separatable. Paired lines became more separable as overlaps decreased in 8-pixel and 16-pixel cases and were distinct in 32-pixel and 64-pixel cases. Images must satisfy the Nyquist sampling criteria to achieve maximal resolution, so their point spread function (PSF) should cover at least 3-pixel. Therefore, the separation of rFRC of paired lines 4-pixel apart means the minimum detectable scale of rFRC map is up to its limit. By involving the rolling operation, we have addressed a major limitation of the previous FRC map[3], which is challenging to correlate the block-wise map to the SR image content (**Supplementary Fig. 15c**, **15d**, **17**).

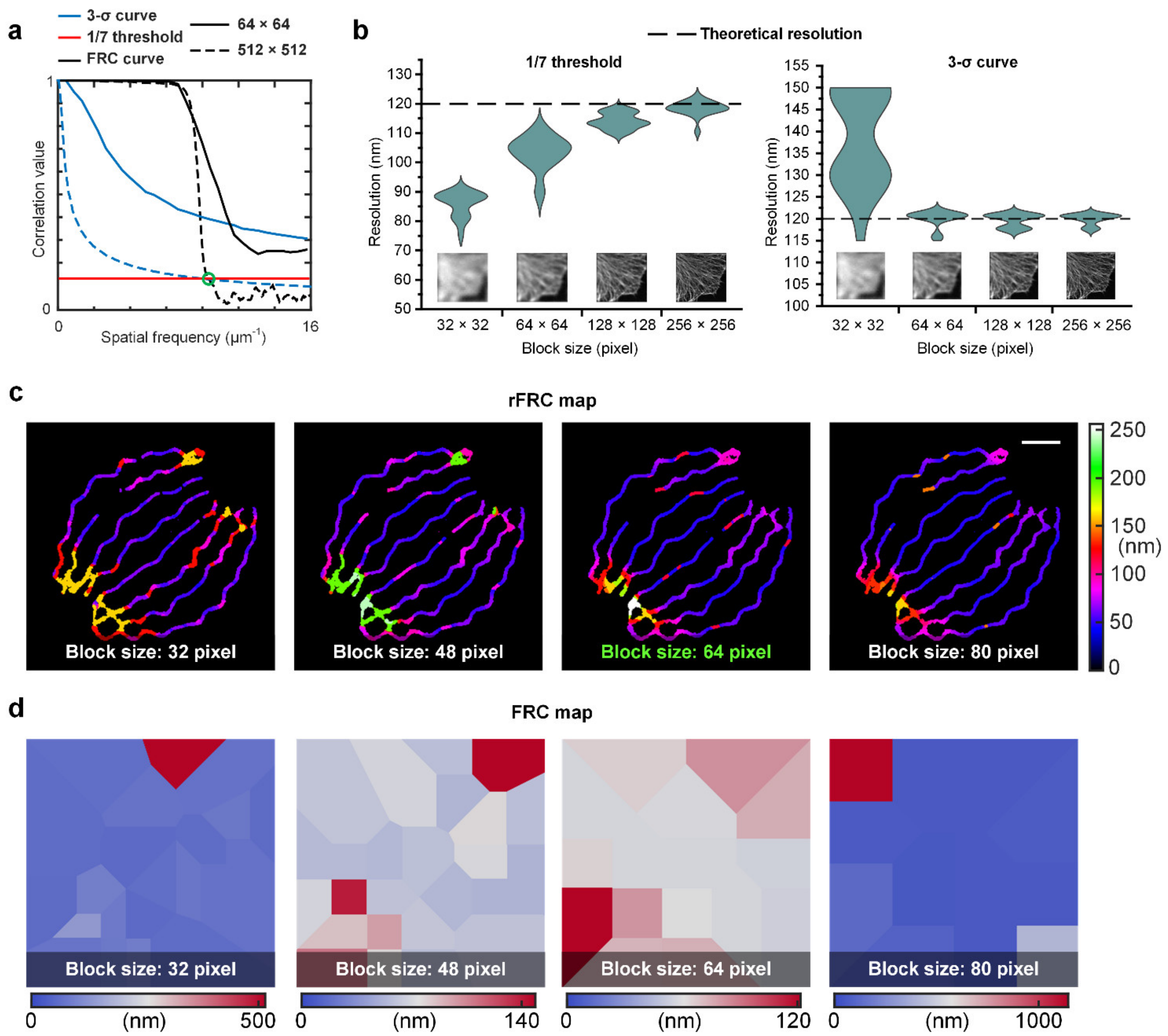


**Supplementary Fig. 15 | The stability of rFRC map.** (**a**) The FRC curve (black), 3σ threshold curve (blue), and 1/7 threshold curve (red) for a 64 × 64 pixels image (solid) and a 512 × 512 pixels image (dashed). For an image with a large size (512 × 512 pixels), the 1/7 threshold attains a similar result to the 3σ curve criterion (green circle). However, for a small image (64 × 64 pixels), the 1/7 threshold is smaller than all correlation values in the FRC curve, failing to yield the cutoff frequency. (**b**) The uncertainty of FRC calculation using different block sizes by 1/7 threshold curve (left) and 3σ threshold curve (right). We downsampled the 2D-STORM captured microtubule image (*c.f.*, **Fig. 2f**, 10 nm pixel size, 4096-pixel number) with 16, 32, 64, and 128 times to create different image sizes and convoluted the resulting images with a 120 nm PSF. After that, Poisson and 5% Gaussian noise were injected into the image. This procedure was repeated 20 times independently and the FRC calculations were performed with different criteria. (**c**) rFRC maps using different block sizes (*c.f.*, **Fig. 1c**). Although the smaller block size (e.g., 32 × 32 pixels) may enable finer mapping, the overall distributions of these rFRC resolution maps using different block sizes are close to each other. On the other hand, the overly small block size may lead to an overconfident resolution value and larger uncertainty. Therefore, to balance the compromise between mapping scale and estimation stability, we chose a block size of 64 × 64 pixels as default in this work. (**d**) FRC maps using different block sizes. Scale bar: 500 nm.

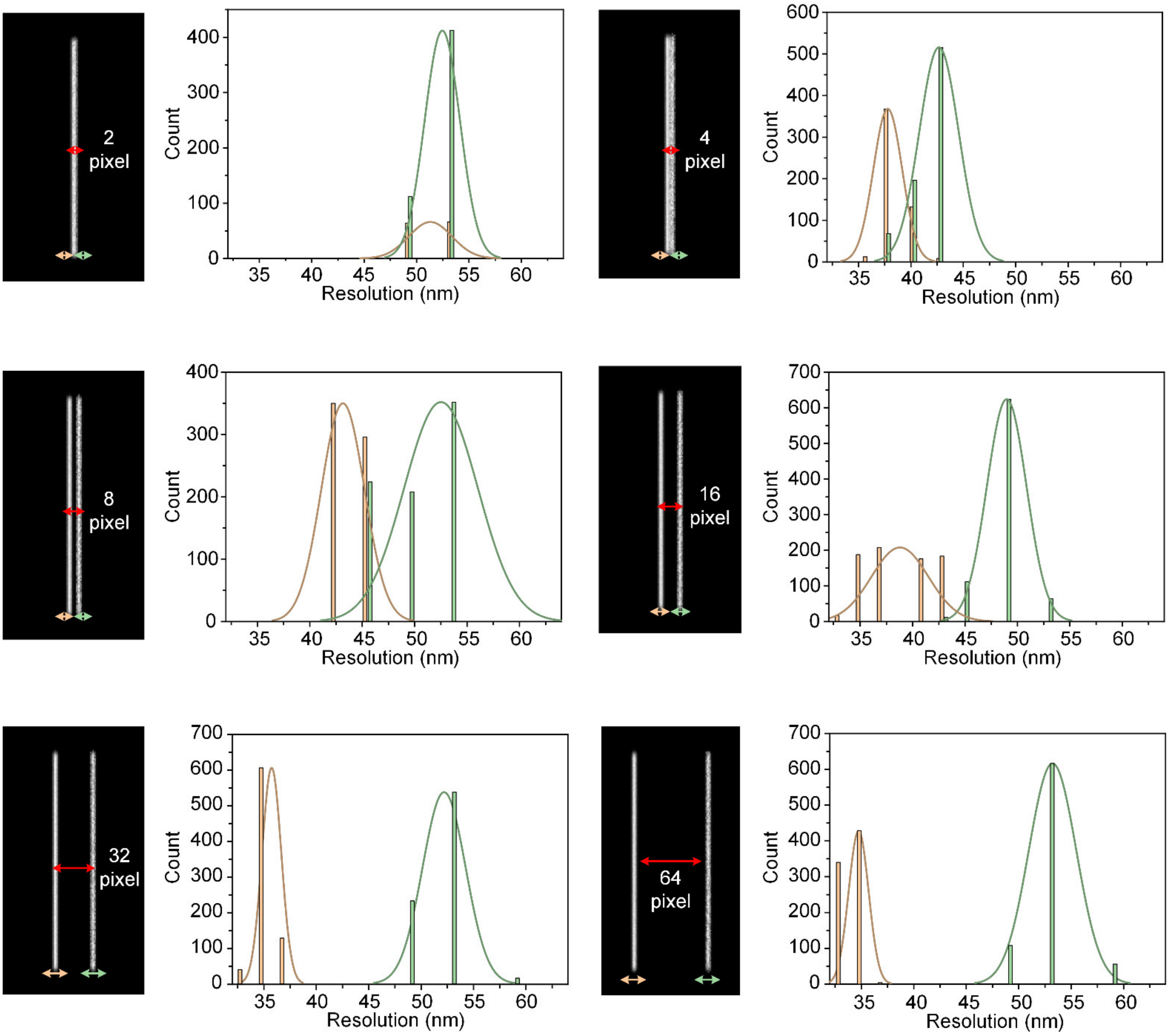


**Supplementary Fig. 16 | The resolvability of rFRC map.** We simulated structures that contained pairs of lines with spacing gradually increases, i.e., 2, 4, 8, 16, 32, 64 pixels (pointed by red arrows), and convoluted them by a PSF with a 4-pixel FWHM (pixel size 10 nm). To test the maximum resolvability of rFRC, we included different noise levels on the two lines. Specifically, we added 10% and 50% Gaussian noise on the left and right lines, respectively. After that, we applied rFRC mapping on the resulting images (left panel) and calculated the FRC value distributions (right panel) of pixels on the left (yellow) and right lines (green).

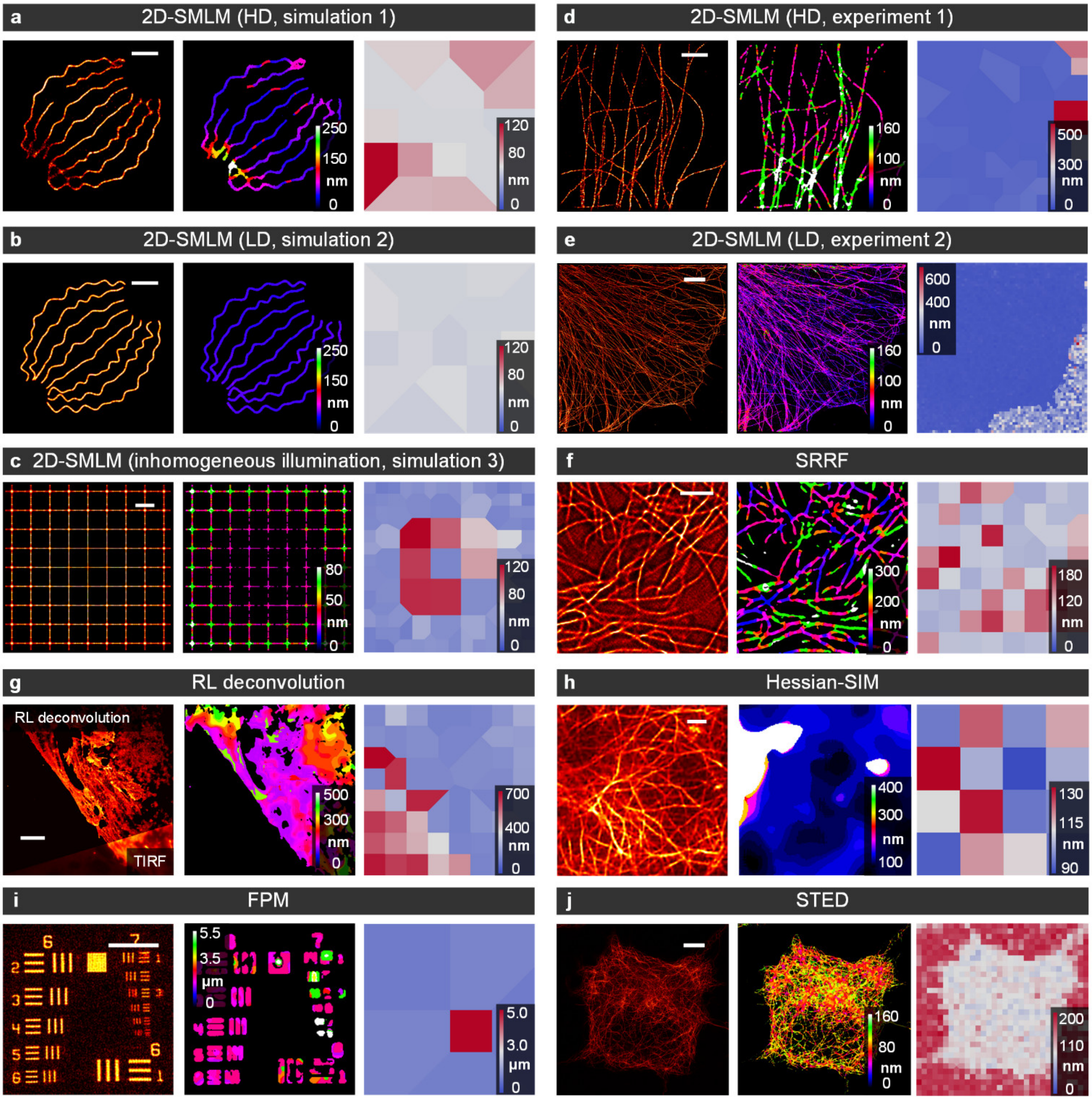


**Supplementary Fig. 17 | rFRC maps versus FRC maps from different modalities.** From left to right: Imaging data, rFRC map, and FRC map. (**a**, **b**) 2D-SMLM simulations with high-density ('HD', **a**) and low-density ('LD', **b**) emitting fluorophores in each frame (*c.f.*, **Fig. 1c**). (**c**) 2D-SMLM simulation with inhomogeneous illumination (*c.f.*, **Fig. 1d**). (**d**, **e**) 2D-SMLM experiments with high-density ('HD', **d**) (*c.f.*, **Supplementary Fig. 7a**) and low-density ('LD', **e**) (*c.f.*, **Fig. 2f**) emitting fluorophores in each frame. (**f**) SRRF experiment (*c.f.*, **Supplementary Fig. 7c**). (**g**) RL deconvolution experiment (*c.f.*, **Supplementary Fig. 19a**). (**h**) Hessian-SIM experiment (*c.f.*, **Supplementary Fig. 18b**). (**i**) FPM simulation (*c.f.*, **Supplementary Fig. 21c**). (**j**) STED experiment (*c.f.*, **Supplementary Fig. 22a**). The FRC map is based on the 1/7 fixed threshold, which may generate unstable calculations. Hence, an inverse distance weight function is involved in interpolating values in all FOVs, while the FRC resolution might not be obtained. This strategy and the calculation on background areas may generate strong false negatives in the resulting FRC map. Scale bars: (**a**, **b**) 500 nm; (**c**, **h**) 1 μm; (**d**, **f**) 2 μm; (**e**, **g**, **j**) 5 μm; (**i**) 50 μm.

#### Supplementary Note 2 | SIM applications.

In structured illumination microscopy (SIM), frequency information is unmixed and stitched from noisy data to achieve super-resolution (SR). As a result, its reconstruction is essentially an ill-posed inverse problem, in which the conventional Wiener reconstruction (Wiener-SIM) will amplify the noise, leading to significant fluctuations in high-frequency components. To moderate this issue, several regularizations were proposed to constrain the reconstruction[6]. For instance, the Hessian-SIM used the Hessian matrix continuity to eliminate random and non-continuous artifacts[6]. Note that the differences between these methods are usually at a fine scale, and thus an evaluation on the corresponding level is essential. Here our rFRC provides a prerequisite for assessing these methods objectively.

In experiments, we applied the Hessian denoising algorithm[6] on the Wiener-SIM reconstruction[7], (**Supplementary Fig. 18a**) to obtain the Hessian-SIM images (**Methods**, **Supplementary Fig. 18b**). Then, we performed the rFRC map to differentiate such subtle differences in the fidelity of conventional Wiener-SIM[7] versus Hessian-SIM[6] (rFRC value, 1.36 versus 1.24) (**Supplementary Fig. 18c**), and in contrast, the RSM detected identical qualities (RSE value, 0.27 versus 0.27) (**Supplementary Fig. 18e**). It is found that only the rFRC value can reflect the difference between Wiener-SIM and Hessian-SIM. We also found that the local qualities in SIM are correlated to the emission intensity of the fluorescent signals (**Supplementary Fig. 18d**), in which the raw images of low SNRs are susceptible to artifacts. The unreliable regions pointed by PANEL (**Supplementary Fig. 18f**) are correlated to the regions under weak illumination of TIRF. Notably, the fixed pattern artifacts of SIM caused by biased parameter estimations or configurations (model bias)[8] cannot be detected by our rFRC method.

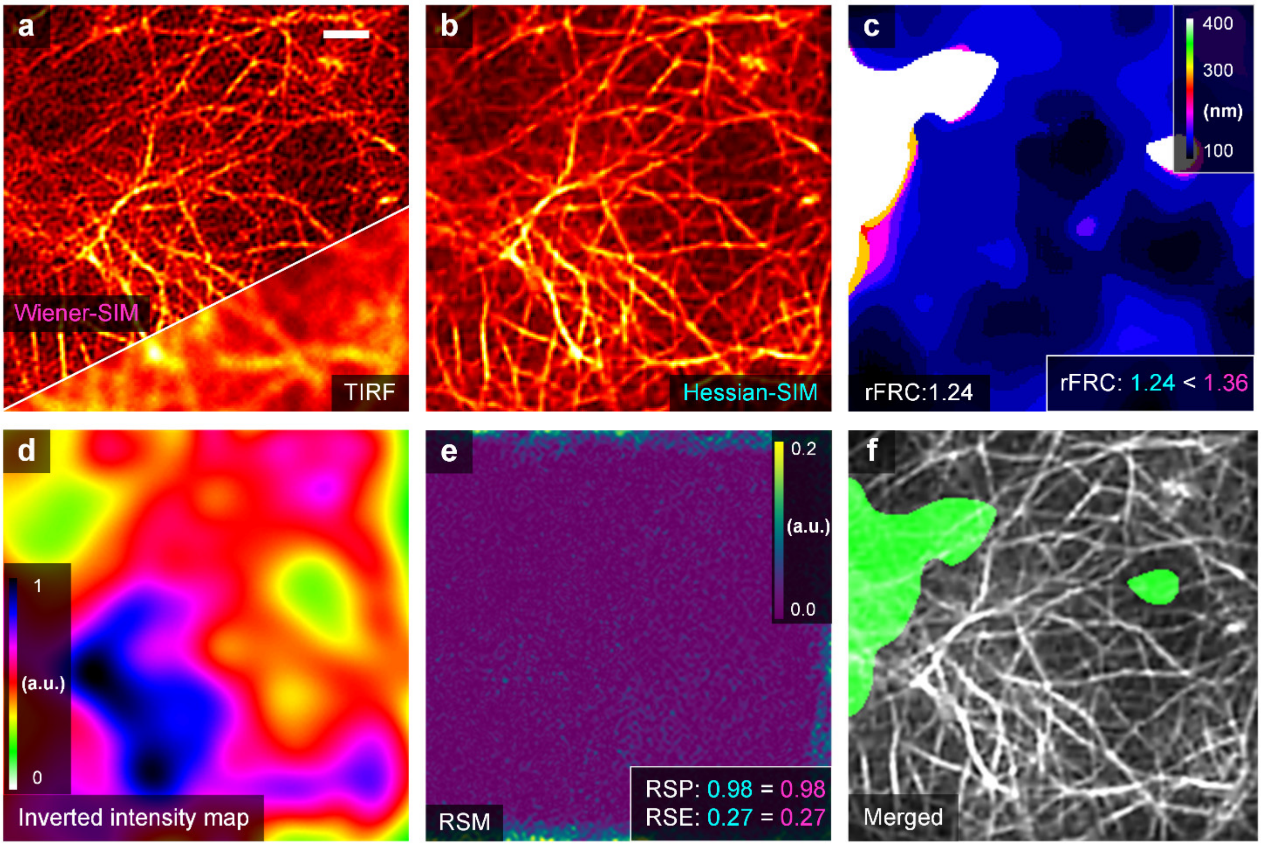


**Supplementary Fig. 18 | rFRC detects the difference between Wiener-SIM and Hessian-SIM. (a)** Representative images of live human umbilical vein endothelial cells (HUVECs) labeled with LifeAct-EGFP under Wiener-SIM (top) and TIRF (bottom) imaging. (**b**) Hessian-SIM result. (**c**) rFRC map of Hessian-SIM. The rFRC, RSP, and RSE values of Wiener-SIM (magenta) and Hessian-SIM (cyan) are shown on the bottom right. Scale bar: 1 µm.

### Supplementary Note 3 | Deconvolution applications.

#### Supplementary Note 3.1 | Determining the number of iterations by rFRC.

Richardson-Lucy (RL) deconvolution[9, 10] has been actively studied for many reasons, including its potential to improve the resolution and contrast of raw images. Nevertheless, the traditional RL algorithm risks amplifying the noise when performing excessive iterations, which extremely limits its applications. Although the noise-insulated low-frequency components may stand stable, the maximum likelihood estimation fits the noise-dominated high-frequency ones to recover the high spatial frequencies, which will lead to wide fluctuations. The common usage requires a *post hoc* visual inspection to determine the best number of iterations. Here, to ascertain the rFRC value readouts guiding this determination for the number of RL iterations, we applied RL to process the TIRF image (**Supplementary Fig. 19a**) and then calculated its corresponding rFRC value of each iteration (**Supplementary Fig. 19b**, right panel of **Supplementary Fig. 19e**). Interestingly, it is noticeable that rFRC values presented a quadratic distribution with the minimum value appearing after 80 iterations. It is similar to the peak signal-to-noise ratio distribution (PSNR, left panel of **Supplementary Fig. 19e**), in which the TIRF-SIM (**Methods**) image is used as ground truth. In contrast, the curve of the resolution-scaled Pearson coefficient (RSP)[3] failed to recapitulate this distribution (middle panel of **Supplementary Fig. 19e**). As demonstrated in **Supplementary Fig. 19d**, the RL deconvolution with 200 iterations produced snowflake-like artifacts, as indicated by the white arrows, which can be confirmed as nonexistent by the referenced TIRF-SIM image. A comprehensive comparison demonstrated that 80-iteration RL optimally enhanced the image contrast with the slightest noise-amplification-induced artifacts. The inverted illumination intensity map (**Supplementary Fig. 19c**) is proportional to the rFRC map (**Supplementary Fig. 19b**), indicating that the local quality in the results of RL deconvolution is highly correlated with the SNR.

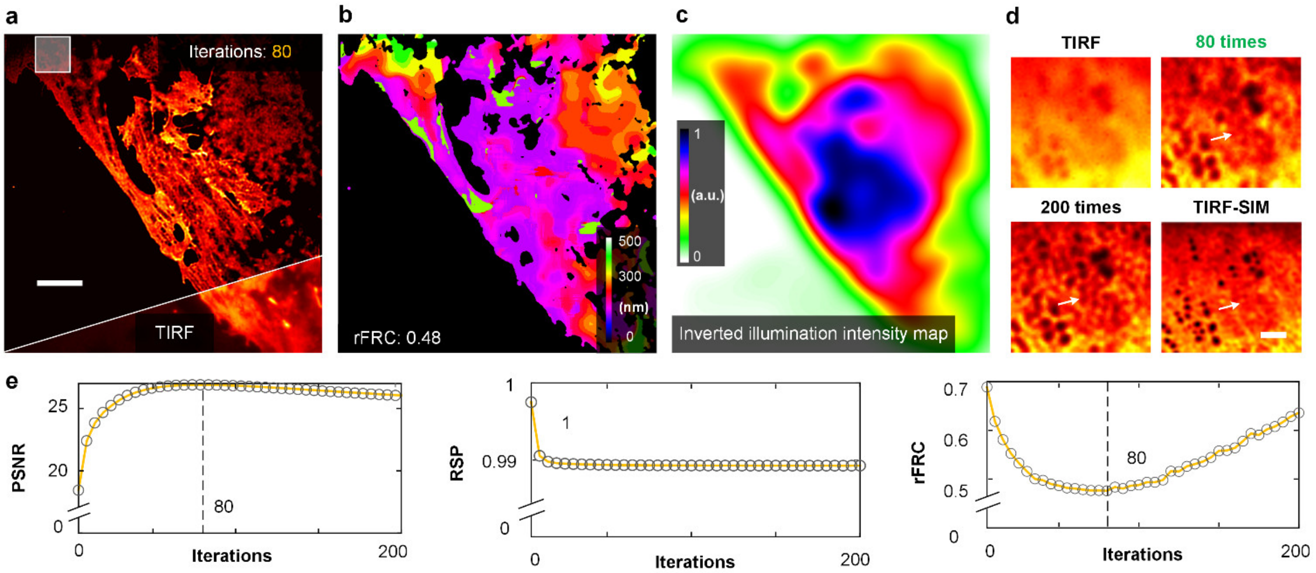


**Supplementary Fig. 19 | Determining deconvolution times by rFRC.** (**a**) Representative results of fixed liver sinusoidal endothelial cells (LSECs) labeled with DiI under RL deconvolution (top) and TIRF (bottom) imaging. (**b**) rFRC map of RL deconvolution result. (**c**) TIRF image convoluted with a large Gaussian kernel and coded with an inverted sJet colormap. (**d**) Magnified views of the white box in (**a**). The original TIRF image, RL deconvolution results with 80 and 200 iterations, and TIRF-SIM results are shown in the top left, top right, bottom left, and bottom right, respectively. (**g**) Curves of the PSNR (versus TIRF-SIM), RSP (versus TIRF), and rFRC values over iterations. (**a**) 5 μm; (**d**) 100 nm;

**Supplementary Note 3.2 | Reducing artifacts by rFRC-based adaptive low-pass filter.**

The FRC can determine the reliable cutoff frequencies (COFs) of the images, indicating that the frequency components are more prominent than the ones corrupted with noise. Because the rFRC can calculate the local COF in different areas of the image, we adaptively low-pass filtered various block-box areas within the entire image:

$$\tilde{\mathcal{F}}\left\{\mathcal{F}(I_{x,y})\cdot OTF\left(F_{x,y}\right)\right\}, \tag{31}$$

where $I_{x,y}$ represents the subset image of the input image, whose center pixel is at the spatial position ($x$, $y$). $OTF(F_{x,y})$ is the optical transfer function (OTF) with the COF $F_{x,y}$, in which the outer COF is set as 0, and the inner COF is 1. In RL deconvolution, the reconstructed image quality is highly related to the corresponding local SNR; hence, the reconstruction result usually has a spatially variant COF. A global FRC filter may not achieve the optimal result (**Supplementary Fig. 20d**, SSIM = 0.32, PSNR = 19.30); in contrast, it can be seen that our designed adaptive rFRC filter yielded a better reconstruction (**Supplementary Fig. 20e**, SSIM = 0.42, PSNR = 21.90).

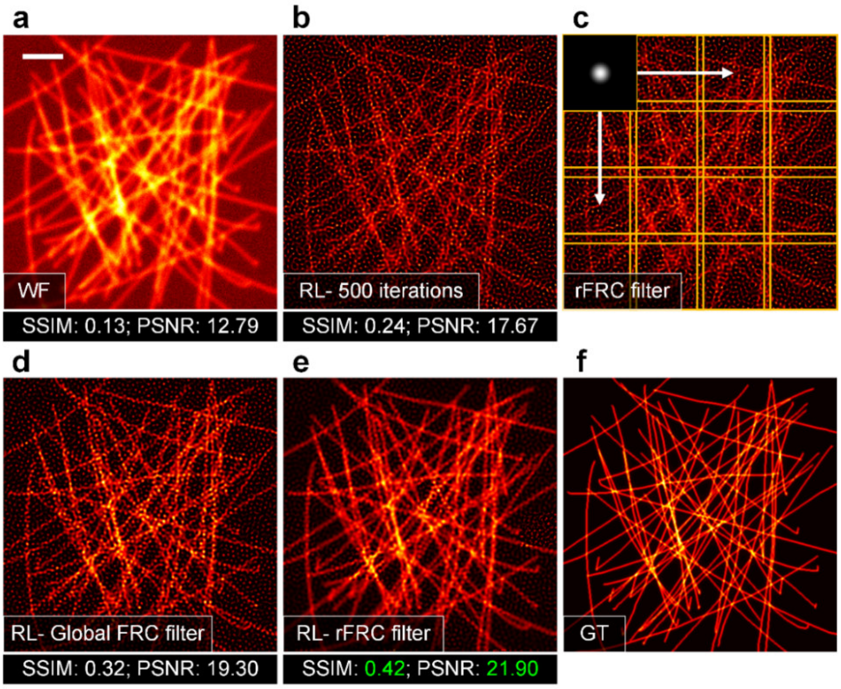


**Supplementary Fig. 20 | Adaptive low-pass filter for RL deconvolution result.** (**a**) The simulated wide-field image. The ground-truth image (**f**) is convoluted by a PSF (FWHM = 240 nm), integrated with Poisson and 10% Gaussian noise. (**b**) Image in (**a**) after RL deconvolution with 500 iterations. (**c**) The adaptive filter workflow. The block size of the filter is set as 64 × 64 pixels, and the overlap between adjacent blocks is set as 4 pixels. (**d**) Image in (**c**) after the global estimated cutoff frequency filter. (**e**) Image (**a**) after the adaptive local filter. (**f**) The ground-truth structures of (**a**). WF: wide-field; GT: ground truth. Scale bar: 2 μm.

#### Supplementary Note 4 | FPM applications.

Fourier ptychographic microscopy (FPM)[11] achieves high-resolution by iteratively stitching together a number of low-resolution images in Fourier space, and it is a coherent imaging modality through a combination of synthetic aperture and phase retrieval concepts. In specific reconstruction process, it updates the objective function between the spatial and Fourier domains iteratively with intensity or pupil constraints. In this case, the noise-contaminated high-frequency components can significantly induce the quality degradation during its spectrum extension. In this experiment, we extended our rFRC applications to FPM for assessing its reconstruction qualities. The United States Air Force (USAF) resolution target was used as the ground-truth sample (**Supplementary Fig. 21a**), and we simulated the FPM imaging process (**Methods**) to create the low-resolution result (**Supplementary Fig. 21b**) and its corresponding high-resolution FPM reconstruction (**Supplementary Fig. 21c**). In **Supplementary Fig. 21d**, it can be seen that the RSM without filtering is prone to small intensity fluctuations belonging to false negative (FP, cyan arrows). In contrast, the rFRC map (**Supplementary Fig. 21e**) accurately represents the quality of FPM reconstruction, pinpointing all the regions of true negative (TN, magenta arrows in **Supplementary Fig. 21f**).

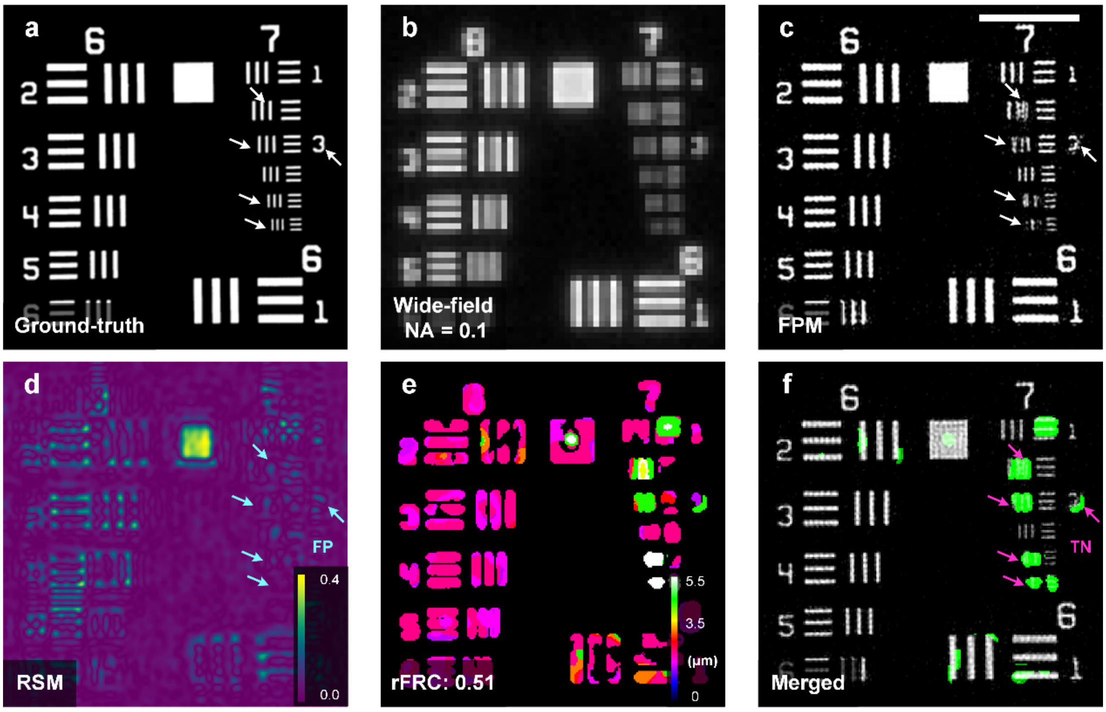


**Supplementary Fig. 21 | USAF target simulation of FPM evaluated by rFRC and PANEL.** (**a**) Simulated ground-truth. (**b**) Wide-field image of (**a**). (**c**) Corresponding FPM reconstruction. (**d**) RSM of FPM. (**e**) rFRC map of FPM. (**f**) Merged image of PANEL (green channel) and FPM (gray channel) results. FP: false positive; TN: true negative. Scale bars: 50 μm.

#### Supplementary Note 5 | STED applications.

In addition to detecting the reconstruction uncertainties, the rFRC map can provide a quantitative resolution metric. Although the original FRC map[3] was used to evaluate the resolution, it is prone to false negatives induced by the background. For example, the mean resolution of the STED[12] estimated by the previous FRC map was given as ~146 nm (**Supplementary Fig. 22b**), which might be an underestimation due to the false negatives caused by the background. In contrast, our rFRC estimated the system resolution as ~92 nm (**Supplementary Fig. 22a**), which is more reasonable. In addition, we provide the 'sJet' color map for visualizing the resolution distribution in higher contrast than that of the previously used 'SQUIRREL-FRC' color map (**Supplementary Fig. 22b**, **22c**).

More importantly, the FRC map is based on the 1/7 fixed threshold[3], while the FRC value might not be obtained in some regions, and an inverse distance weight function is involved in interpolating values in all FOVs. This strategy may generate false negatives in the resulting FRC map, as indicated by the yellow arrow in **Supplementary Fig. 7b**. Therefore, this FRC map could not reflect the fine artifacts in the MLE reconstruction. In contrast, our rFRC map used the $3\sigma$ curve to determine the cutoff frequency with applying a background threshold filter to remove the background FRC value, which effectively reduced possible false negatives. As a result, the proposed rFRC map could favorably achieve challenging SR scale quality mapping, as demonstrated in **Supplementary Fig. 7a** and **Supplementary Fig. 17**. Nevertheless, we still provided both the 1/7 hard threshold and $3\sigma$-curve-based resolution mapping features in the PANELJ Fiji/ImageJ plugin for further potential applications.

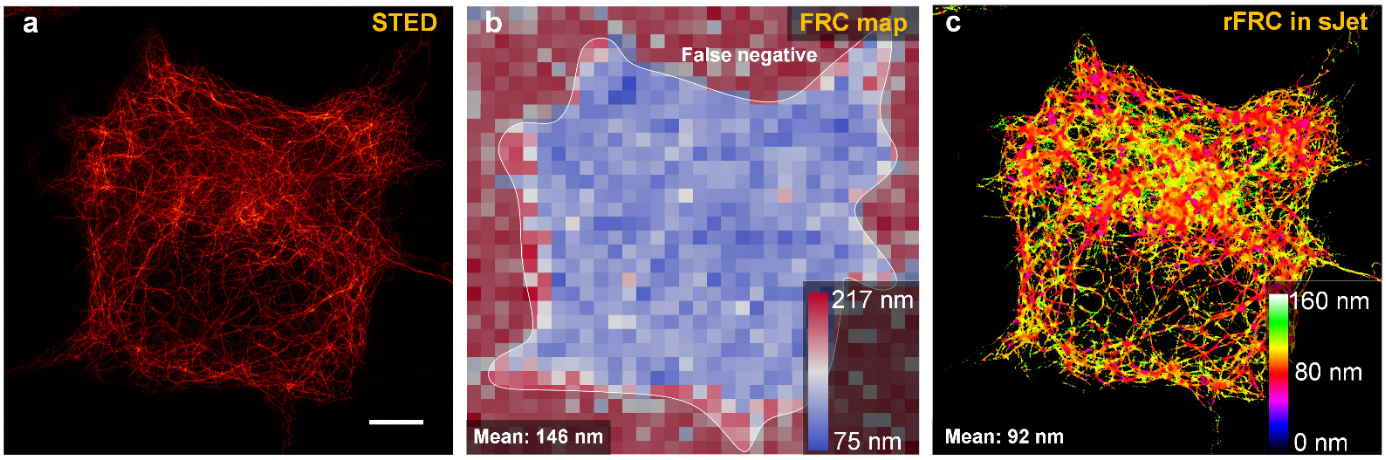


**Supplementary Fig. 22 | False-negative induced by the background.** SiR-tubulin-labeled microtubules seen under gSTED[13] (**a**); FRC map with 64-pixel block size coded with the SQUIRREL-FRC color map (**b**); rFRC map coded with the sJet color map (**c**). Scale bar: 5 μm.

#### Supplementary Note 6 | Comparisons of rFRC, RSM, and SSIM.

First, we intend to demonstrate the reliability of the proposed assessment. As seen in **Supplementary Fig. 23**, a series of filaments with different distances were convoluted with a wide-field PSF (NA=1.4). We gradually increased the noise level in the raw image (along the yellow arrow, **Supplementary Fig. 23b**) and showed the results after RL deconvolution (**Supplementary Fig. 23c**). Compared to the ground truth, significant artifacts appeared in the white dashed box in **Supplementary Fig. 23c**, which were successfully detected by our rFRC map (**Supplementary Fig. 23d**) but not the RSM (**Supplementary Fig. 23i**). Under the identical configurations of rFRC mapping, we can see these spatial methods failed to highlight such unreliable regions, in which we showed the structural similarity (SSIM)[14] map (**Supplementary Fig. 23e**) and the subtraction (**Supplementary Fig. 23f**). Beyond that, it is also worth noting that the rFRC maps formed by *Reconstruction1 and Reconstruction2* (**Supplementary Fig. 23d**) or *Reconstruction1 and Ground Truth* (**Supplementary Fig. 23g**) matched perfectly, indicating that our method can evaluate the reconstructed image quality without the ground truth.

Second, our rFRC map can also be used as a generalized metric to quantify the difference between the reconstruction and ground truth, and overcomes the natural defect of SSIM. In the SSIM map (**Supplementary Fig. 23h**), we can see the existence of strong false negatives, making the true negatives difficult to dissect. The region inside the yellow box (**Supplementary Fig. 23b**) was set as noise-free, thus there was no difference between the two independent reconstructions (**Supplementary Fig. 23d-23f**). Here, the reconstruction within this region (**Supplementary Fig. 23c**) was almost identical to the ground truth (**Supplementary Fig. 23a**). Interestingly, both the inverted SSIM map and RSM (**Supplementary Fig. 23h** and **23i**) still provided small values, indicating a false negative. In contrast, the rFRC map between *Reconstruction1* and *Ground Truth* (**Supplementary Fig. 23g**) remains empty for this region, which is fairly more reasonable.

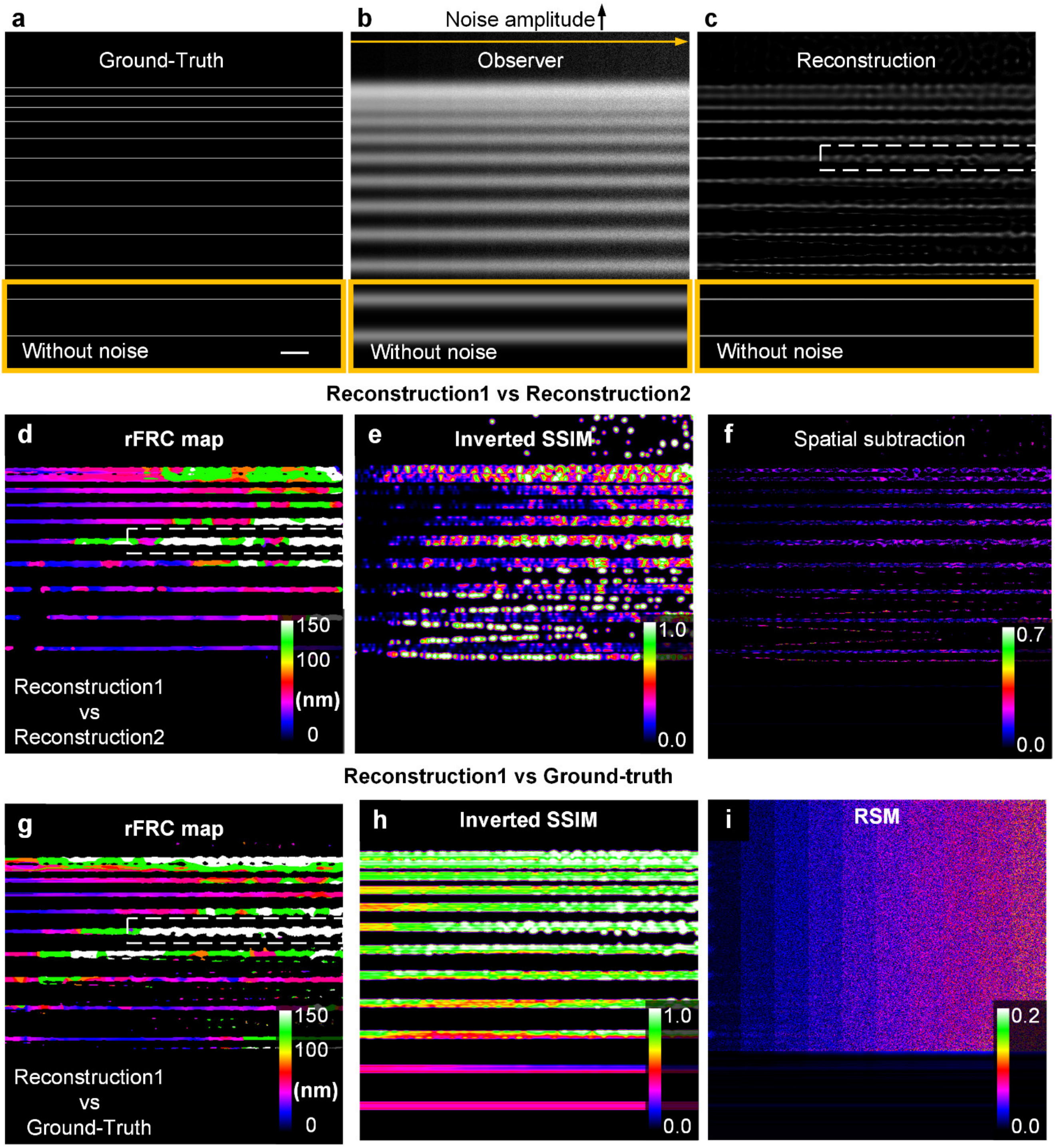


**Supplementary Fig. 23 | Synthesizing noise of different amplitudes to evaluate the performance of the rFRC, RSM, and SSIM.** (**a**) Ground-truth sample. (**b**) Wide-field image. (**c**) RL deconvolution result. (**d**) rFRC map from two reconstructions (*Reconstruction1* vs *Reconstruction2*). (**e**) Inverted SSIM map from two reconstructions (*Reconstruction1* vs *Reconstruction2*). (**f**) Spatial subtraction results from two reconstructions (*Reconstruction1* vs *Reconstruction2*). (**g**) FRC map from reconstruction and ground truth (*Reconstruction1* vs *Ground Truth*). (**g**) Inverted SSIM map from reconstruction and ground truth (*Reconstruction1* vs *Ground Truth*). (**i**) RSM of (**c**). Scale bar: 500 nm.

#### Supplementary Note 7 | The uncertainties in learning-based applications.

##### Supplementary Note 7.1 | The leaked model uncertainty detection.

In theory, as a model-independent method, the rFRC using two captures cannot directly detect the model uncertainty. However, in practice, as a purely data-driven approach, the deep learning model has no stationary form in weights that acquire only after learning from the training data. As discussed in ref[15], the model uncertainty and data uncertainty may be not mutually exclusive. To concisely demonstrate this mechanism, we synthesized four simple types of structures before sampling them to a sparse form[16] (**Methods**, **Supplementary Fig. 24**), and the structures with or without sparse sampling were used as the model input or ground truth. The rectangular shapes were used as the training dataset, and the square, triangular, and circular shapes were used to test the U-net predictions (**Methods**, **Supplementary Fig. 13**). Because it is hard to disentangle deep-learning artifacts due to mode uncertainty or data uncertainty in actual experiments, we intentionally simulated the univocal shapes to explore this problem[16]. Since the squares can be regarded as a subset of rectangles, using the sparse squares/rectangles as input, the network was free from model uncertainty and thus predicted the corresponding shapes accurately (**Supplementary Fig. 24b**, **24c**). In contrast, when presenting the network with the out-of-distribution shapes (triangular or circular), the predicted results still approximated the corresponding structures with the learned rectangular shapes[16], primarily caused by the model uncertainty (**Supplementary Fig. 24b**, **24c**).

To study how the model uncertainty leaked into the data uncertainty, we sampled identical structures twice to generate two predictions, denoted as *Prediction1* and *Prediction2* in **Supplementary Fig. 24c**. Confirming our assertion, the average intersection over union (IoU, also known as the Jaccard index) values between *Prediction1* and *Prediction2* shared a distribution identical to that between the *Ground-truth* and *Prediction1* (**Supplementary Fig. 24d** and **24e**). Moreover, the pattern of the mean rFRC values was exact opposite to those of the IoU values (**Supplementary Fig. 24f**). The predictions of the square/triangular shapes gave small rFRC metrics (1.56/1.65), and that of the triangular/circular shapes led to much larger rFRC metrics (4.19 and 4.89). Based on above proof-of-principle simulations, we can find the only probable reason for the rFRC metrics increasing, is the model uncertainty leaking into the measured data uncertainty. Therefore, using the rFRC metric on the two individual predictions, we can detect both the data and the leaked model uncertainty of learning-based approaches without knowing the ground truth.

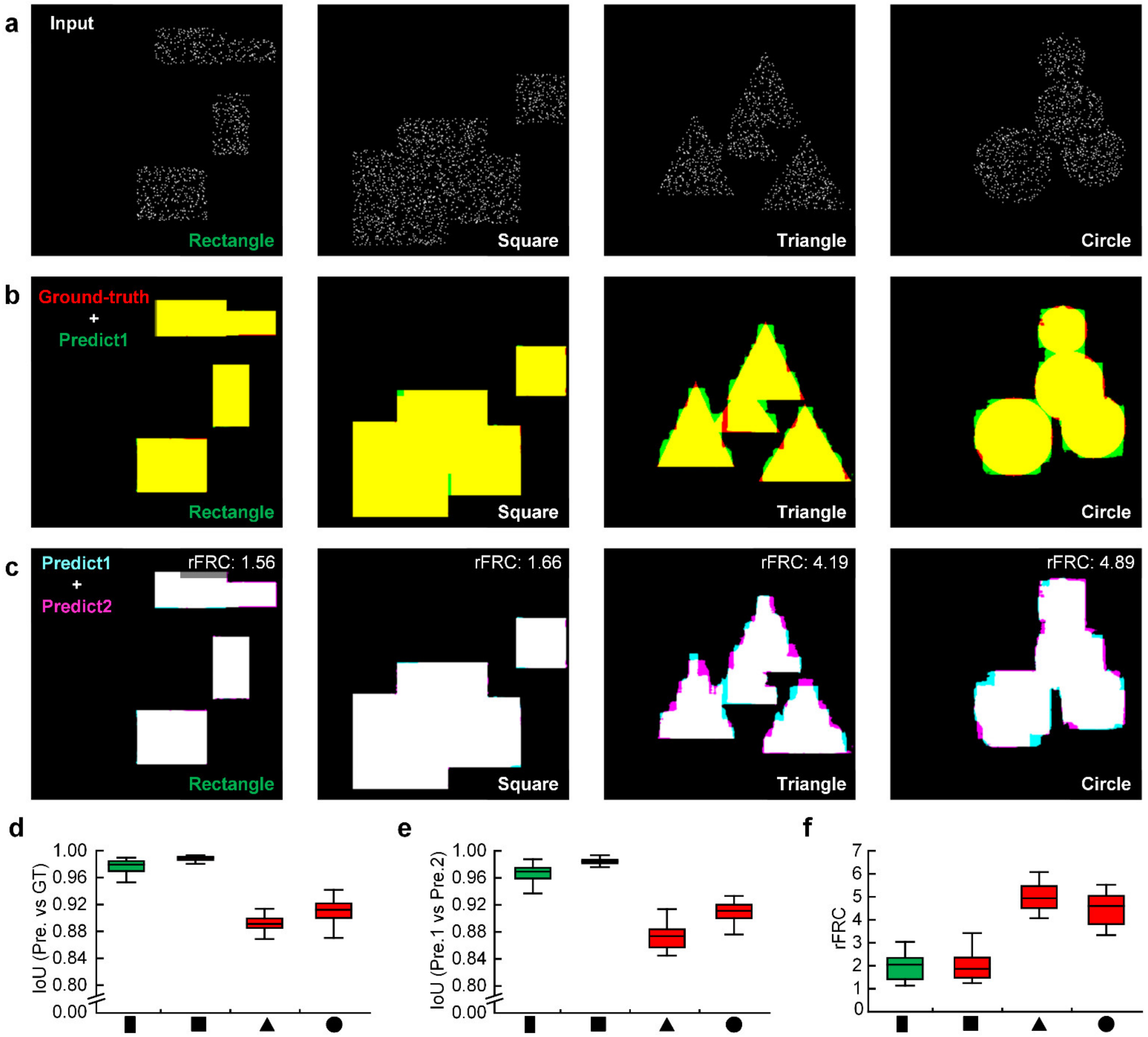


**Supplementary Fig. 24 | Simulation of univocal shapes uncovers that the model uncertainty leaks to the data uncertainty.** Rectangles (left) are used as the training dataset, and the other geometrical shapes (squares, circles, and triangles) denote the test dataset (from left to right). (**a**) Representative sparsely sampled input of corresponding geometry ('Input'). (b) Merged images using predicted images and the related ground-truth images (ground-truth: red channel; prediction 1: green channel). (**c**) Merged images using two predicted images (prediction 1: cyan channel; prediction 2: magenta channel). The corresponding input images of 'Prediction1' and 'Prediction2' are sampled independently. (**d**) Average IoU values from the prediction versus the ground truth (median values from left to right: 0.97, 0.99, 0.89, 0.91). (**e**) Average IoU values between two predictions (median values from left to right: 0.97, 0.98, 0.87, 0.91). (**f**) Average rFRC values between two predictions (median values from left to right: 2.05, 1.86, 4.94, 4.60).

**Supplementary Note 7.2 | A strategy for assessing both model and data uncertainties.**

Since exact Bayesian inference is computationally intractable for neural networks (Bayesian neural networks, BNNs)[17], many feasible approximations have been proposed, including the Monte Carlo dropout[18] (MC dropout), variational Bayesian methods[19], and deep ensemble[20]. Although these approximations allow the BNNs to function at some levels, they still rely on crucial assumptions. Compared to the conventional NNs, distribution forms of network predictions and weights need more complicated parameters, and most approximations need modifications to the network structures and training processes. These modifications may compromise the performance of NNs and were inconvenient in most application cases. Thus, the BNNs may not be a practical choice for uncertainty assessment when considering these imperfections.

In traditional optical imaging, the model uncertainty assessments rely on specific model calibration procedures. However, as a purely data-driven approach, the deep learning approach has no stationary form in weights which only learns the representations of training data. In another word, the predictions of the out-of-distribution input data (images) would be more sensitive (with more predicted fluctuations) to the potential influences, such as model weights changes. Like the deep ensemble approach[20], it conceptually utilized an ensemble of models for quantifying predictive uncertainty with respect to the model parameters (model uncertainty). Inspired by this, we also introduce an explicit framework to assess both model and data uncertainties. First, we independently trained two models on the same dataset (with different random initializations and optimization processes) to reflect the model uncertainty, and the rFRC mapping was used to represent the model uncertainty. This approach resembles the frequentist approach to estimating uncertainty, which is simple to implement, flexible to parallelize, and no hyperparameter tuning required. Second, we sampled the input data (images) twice to evaluate the data uncertainty, and we then applied the rFRC to assess the data uncertainty as did in the optical modalities.

Using a proof-of-principle experiment under the same configuration as in **Supplementary Fig. 24**, we verified the possibility of using this strategy to extend our rFRC for assessing both data uncertainty and model uncertainty (**Supplementary Fig. 25**). They are first quantified using the IoU metric against the ground truth as references. For model uncertainty, we trained 2 networks independently and then fed them with the same sparsely sampled data to obtain *Prediction1* and *Prediction2*. To explore the generality, we also trained 8 networks independently for obtaining 8 predictions, in which the patterns of the IoU (between *Predcition1* and *Prediction2*) and the average of 8 predictions are consistent with the IoU (between the predictions and ground truth) (**Supplementary Fig. 25b**). Next, for data uncertainty, we sampled the same data twice or eight times and followed the same evaluation procedure mentioned. Interestingly, we found the predictive differences of multiple models are rather small (right panel of **Supplementary Fig. 25b**, IoU (2) and (8)), reflecting that, most model uncertainty was leaked to the data uncertainty in this case. Overall, according to these results (**Supplementary Fig. 25**), we might use this strategy to evaluate data uncertainty and model uncertainty.

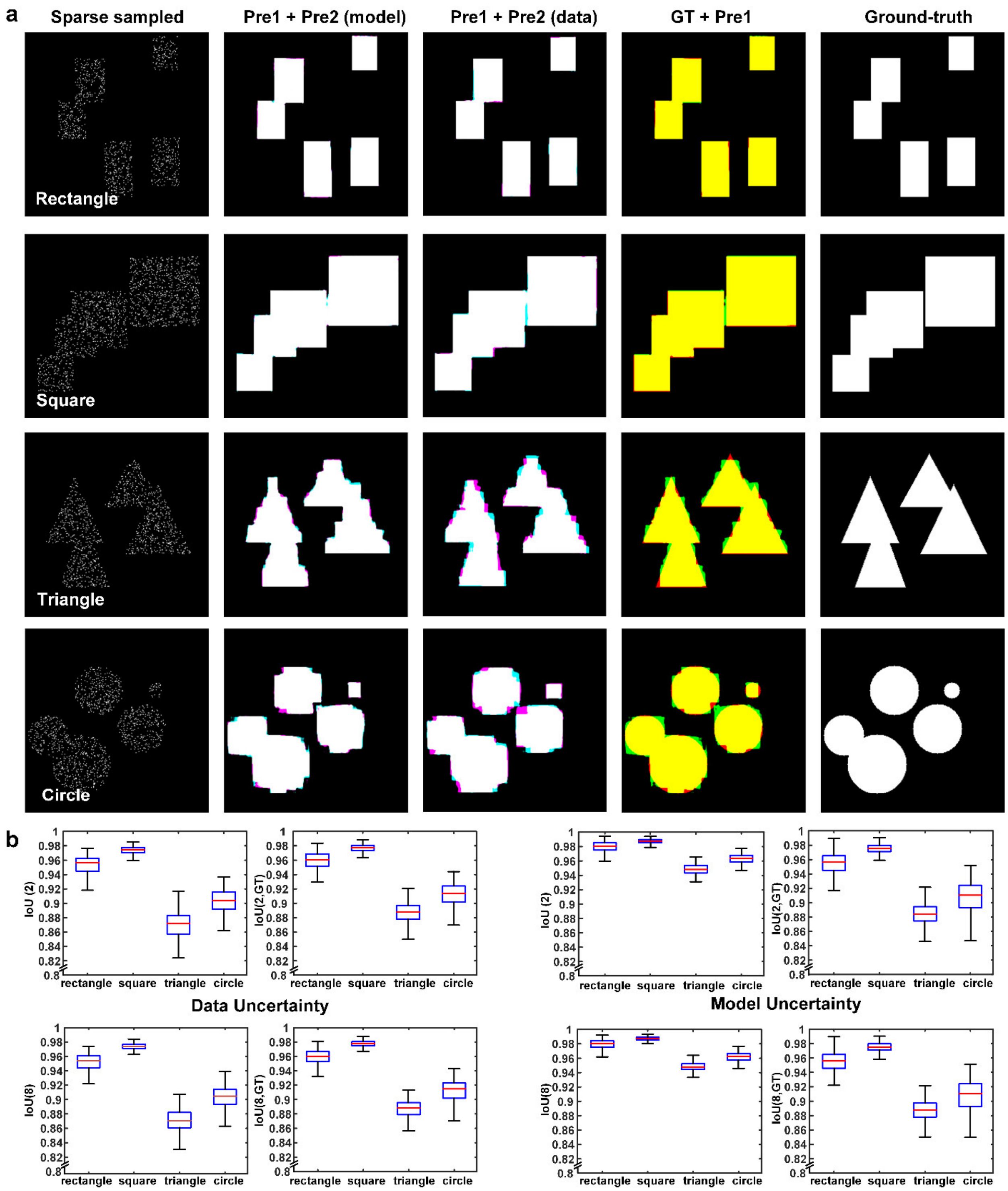


**Supplementary Fig. 25 | Data and model uncertainty quantification of sparse sampling simulated demonstration.** Rectangles are used as the training dataset, and other geometrical shapes (squares, triangles, and circles) are used as a test dataset (from top to bottom). (**a**) From left to right: Input (representative sparsely sampled input of the corresponding geometry); Prediction1 + Prediction2 (merged image of two predictions when two neural networks are **trained** independently); Prediction1 + Prediction2 (merged image of two predictions when two input images are **sampled** independently); GT + Prediction (merged image of PANEL in green channel and the predicted result in red channel); Ground truth (before sparse sampling). (**b**) Data uncertainty (top) and model uncertainty (bottom) quantification using the intersection over union (IoU) as an index. IoU (2): IoU between 2 predictions; IoU (8): IoU between 8 predictions; IoU (2, GT): IoU between 1 prediction and the ground truth; IoU (8, GT): IoU between 8 predictions and the ground truth.

#### Supplementary Note 8 | Single-frame strategies for rFRC mapping.

##### Supplementary Note 8.1 | Single-frame rFRC for optical imaging applications.

The FRC calculation needs statistically independent image pairs sharing exact details but different noise realizations. In some optical imaging applications, such as SMLM[21, 22], SRRF[23], and SOFI[24], these modalities can produce statistically independent images by dividing the input image sequence into two subsets and reconstructing them independently. Other modalities can even sample twice directly to create the necessary image pair[4]. However, some modalities, require multiple inseparable measurements to generate a single piece of SR content, resulting in difficulty in obtaining statistically independent image subsets.

Following the examples of Sami Koho et al.[5], we provided the single-frame rFRC as a supplement (**Supplementary Fig. 26a**). Considering each camera pixel being sampled independently, we divided a single frame into four subsets to create two image pairs with identical details but different noises. These two pairs were formed according to the pixels at (even, even) and (odd, odd) or (even, odd) and (odd, even) row/column indexes, as shown in *Step 1* in **Supplementary Fig. 26a**. This operation yields a single-pixel shift in both $x$ and $y$ directions in the image pairs and this spatial shift results in a $e^{-i2\pi sr/N}$ frequency phase modulation during FRC calculation. $s=\sqrt{x_0^{\ 2}+y_0^{\ 2}}$ is the total length of the shift, and $r$ is the radius in the FRC calculation. As the calibration procedure described by Sami Koho et al.[5], we correct this bias by the following equation:

$$r_{tf}=\frac{1}{a\cdot e^{c\cdot r_{sf}-b}+d}, \tag{13}$$

where $r_{tf}$ and $r_{sf}$ represent the two-frame and one-frame FRC cutoff frequencies (COFs), respectively. The 4 parameters $a$, $b$, $c$ and $d$ are all experimentally fitted from the data ($a$ = 0.9599, $b$ = 0.9798, $c$ = 13.9044, $d$ = 0.5515)[5]. In addition, we calculate the FRC values of two image pairs and average them to deal with special spectral-domain symmetries that arise when the details in an image are majorly oriented in a single direction. As the lateral dimensions of the four sub-images being identical and half the size of the original image, we resize the resulting rFRC map to the original image size with interpolation for better visualization.

##### Supplementary Note 8.2 | Single-frame rFRC for learning-based imaging.

Deep-learning reconstruction techniques differ from the conventional optical imaging methods in which the adjacent pixels in the output results from deep neural networks (DNNs) may share similar noise characteristics. Therefore, the above proposed single-frame strategy for the rFRC used in the optical imaging method is unsuitable for examining DNN reconstructions. Inspired by DeepFool[25] and other works[26], we also create a strategy to perform single-frame rFRC calculations for DNNs. DeepFool added small and invisible perturbations to the input image, and the corresponding classification result of the DNNs was far from the truth. Similarly, we add two independent Gaussian noises to the original input image for artificially creating two frames as the input pair (**Supplementary Fig. 26b**). Then, after the DNNs reconstruct these two input images individually, the resulting two output images are used to calculate the rFRC map following the same procedure. When the independent Gaussian noise is added into the input image, these small perturbations may profoundly change the DNN reconstructions. Remarkably, the more significant difference exists between these two reconstructions, the more the reconstructions will deviate from the real object. It also agrees with the data uncertainty concept proposed by Cameron Buckner for BNNs[27], in which the more considerable the variation induced by this perturbation, the worse the intrinsic error.

**Test on learning-based SIM:** In principle, 2D-SIM achieves SR by taking nine images of a wide-field microscope; thus, it provides the ideal pairs of low-resolution (LR) and SR images to train learning-based networks, such as the TIRF2SIM[28] and DFGAN-SIM[29] algorithms. Because SIM only modestly increases the spatial resolution (~two-fold), it may represent an ideal model for deep-learning algorithms to transform wide-field images into SR-SIM images. Despite their excellent visual effects, we wonder about the accuracy of the network prediction. By applying the single-frame rFRC calculation strategy, we evaluated spatial frequency extension by the TIRF2SIM algorithm (**Supplementary Fig. 27a**-**27d**)[28]. While the deep network deduced simple CCPs from an LR image with high accuracy, the PANEL map highlighted these erroneous regions (green regions, **Supplementary Fig. 27d**). For example, one large postulated CCP (the bottom arrow in **Supplementary Fig. 27b**) was two adjoined CCPs under TIRF-SIM (the bottom arrow in **Supplementary Fig. 27c**). Another algorithm, DFGAN-SIM, also predicted the simple microtubule filaments (under the grazing incidence illumination, GI)[30] with high fidelity. However, our rFRC revealed relatively increased uncertainties at the intersections (**Supplementary Fig. 27f-27g**). In contrast, when DFGAN-SIM was used to postulate intricate mitochondrial cristae structures from the raw wide-field images, the predicted image was different from the ground truth in many aspects, despite the overall visual resemblance (**Supplementary Fig. 27k-27o**). It is clear that the segmented rFRC map again highlighted these structural similarities and dissimilarities, underscoring its application in quantifying local uncertainties in SR images postulated by any learning-based method.

Then, we tested the single-frame rFRC maps in images corrupted with noises of different amplitudes (**Supplementary Fig. 28**). Small noise amplitudes cannot disturb the network predictions to detect the holistic view of uncertainty (black areas in 1% ~ 7% from **Supplementary Fig. 28**). Once the noise amplitude is capable of probing the overall uncertainty distribution of the predictions (9% from **Supplementary Fig. 28**), we stop increasing the noise amplitude and choose this noise level as a proper amplitude to generate two frames for our rFRC mapping.

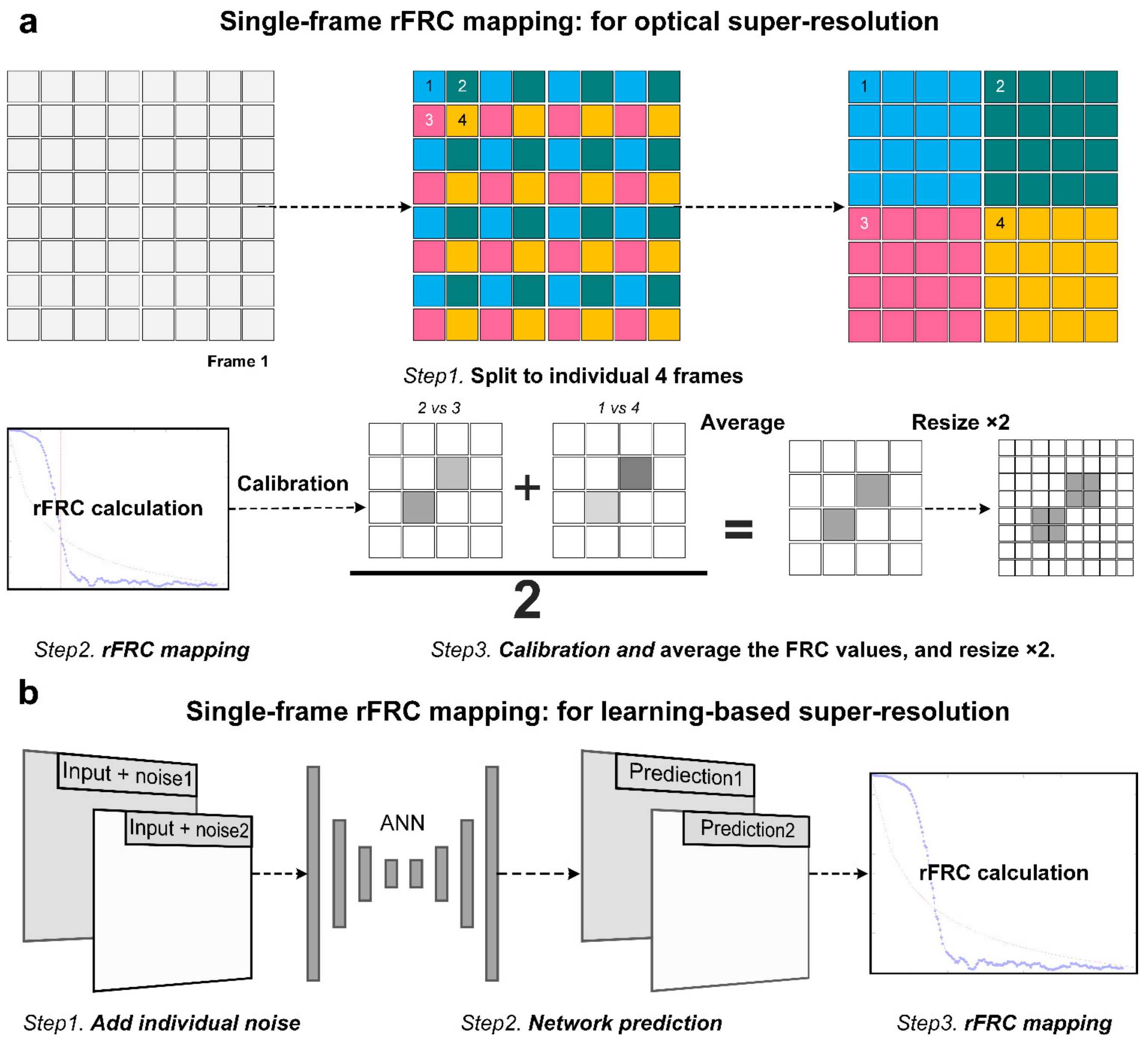


**Supplementary Fig. 26 | The rFRC calculation from a single frame.** (**a**) The strategy for optical super-resolution. (**b**) The strategy for learning-based super-resolution. ANN: Artificial neural network.

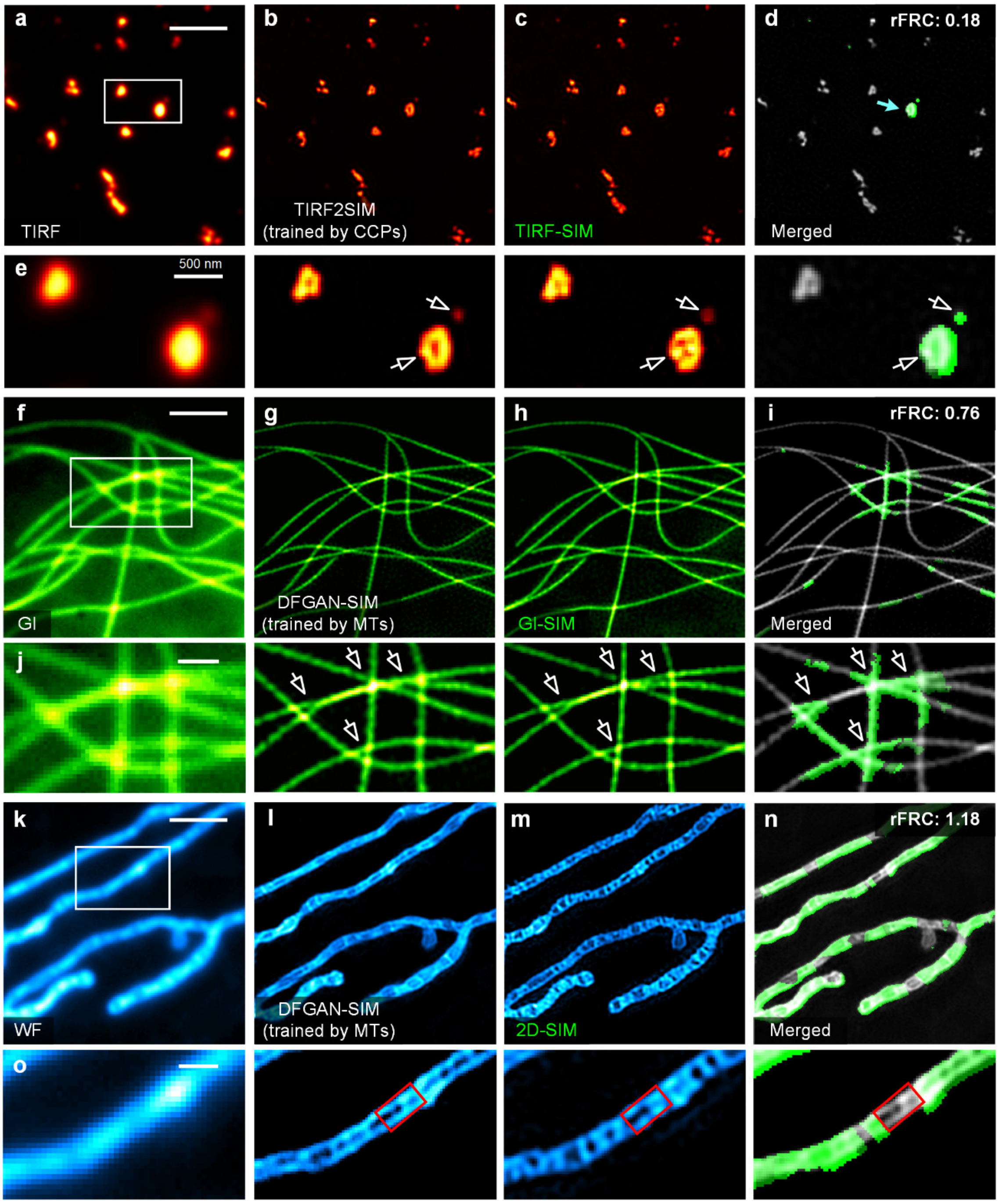


**Supplementary Fig. 27 | Learning-based SIM evaluation.** (**a**) TIRF2SIM result of CCPs (gene-edited SUM159 cells expressing AP2-eGFP). The network was trained on CCP structures ('CCPs'), and the input of which is a single-frame TIRF image. (**b**) Corresponding TIRF-SIM image. (**c**) Input TIRF image. (**d**) Merged image of the PANEL (green channel) and TIRF2SIM (gray channel) results. (**e**) Corresponding magnified views of the white box in (**a**). (**f**) DFGAN-SIM result of microtubules (enconsin-mEmerald in COS-7 cells). The network was trained on different microtubule structures ('MTs'), in which the input is 9 frames of raw data. (**g**) Corresponding GI-SIM image. (**h**) Averaged input GI image. (**i**) Merged image of the PANEL (green channel) and DFGAN-SIM (gray channel) results. (**j**) Corresponding magnified views of the white box in (**f**). (**f**) DFGAN-SIM results of mitochondrial cristae (MitoTracker Green in COS-7 cells). The network was trained on microtubule structures, and its input is nine frames of raw data. (**g**) Corresponding 2D-SIM image. (**h**) Averaged input WF image. (**i**) Merged image of the PANEL (green channel) and DFGAN-SIM (gray channel) results. (**j**) Corresponding magnified views of the white box in (**f**). Scale bars: (**a**, **f**, **k**) 2 μm; (**e**, **j**, **o**) 500 nm.

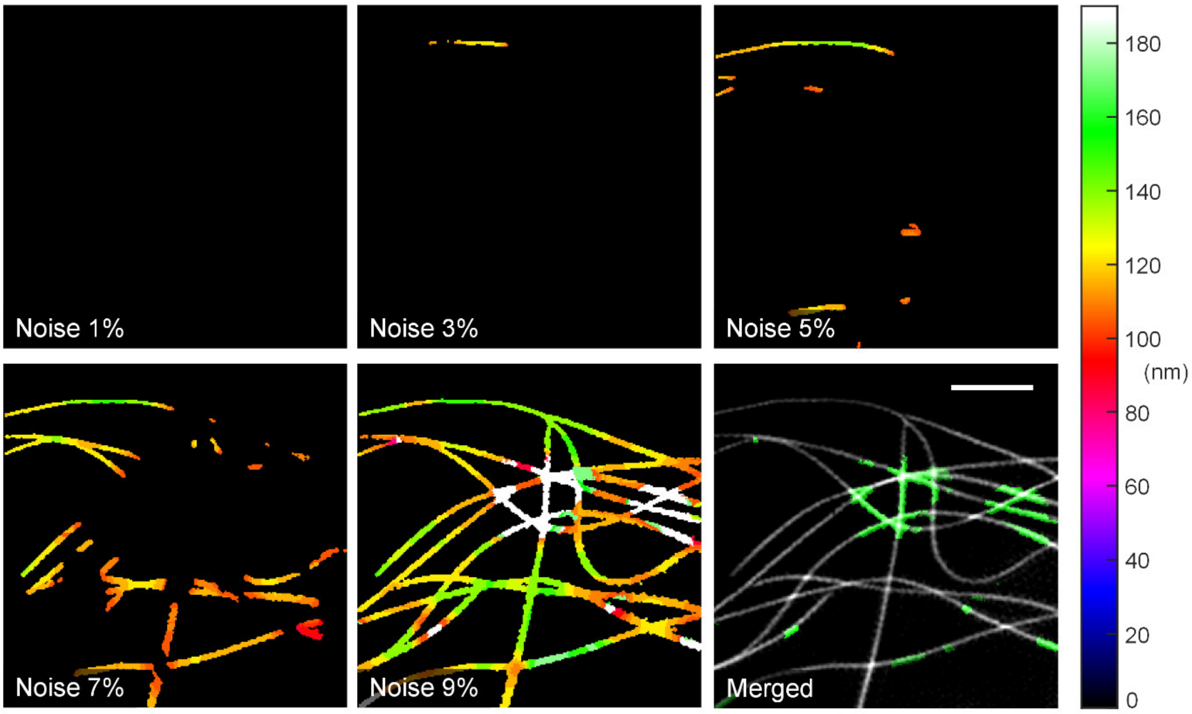


**Supplementary Fig. 28 | Illustration of rFRC maps with different noise amplitudes added (*c.f.*, Supplementary Fig. 27f-27j).** The rFRC maps except for the image at the right bottom. The corresponding percentage sign represents the amplitude (variance) of noise (1% equal to '0.01 × maximum intensity of image'). Right bottom: Merged image of the PANEL (green) and DFGAN-SIM (gray) result. Scale bars: 2 μm.

#### Supplementary Note 9 | Limitations.

In this part, we discussed the possible caveats of the rFRC and RSM maps, revealing the holistic view of our quantification, including its inherent limitations. Inspired by the Bayesian neural network[15], we defined two major types of uncertainty: the data uncertainty and the model uncertainty. The data uncertainty is mainly induced by the combined effects of noise/sampling. The model uncertainty is primarily caused by the existing distance between the established model and its real-world counterpart (or networks' ignorance of the out-of-distribution data). However, based on the underlying theory of the corresponding models, the model-related bias can be detected and reduced by careful system calibration[8, 31, 32] in optical imaging, or suppressed by a specifically designed strategy[33] and enough training data[15] in learning-based applications. On the other hand, data uncertainty is fundamentally model-independent, inevitable, and difficult to remove by system calibration (or adding more training datasets in learning-based scope).

In optical imaging applications, our model-independent rFRC can measure the data uncertainty to reflect the data error, and is limited to uncover the model uncertainty-induced reconstruction quality deterioration. For example, the PSF mismatch (theoretical PSF versus real PSF) induced by instrument imperfections and the sample-induced aberrations will introduce the biased estimation in SMLM[34], which can be compensated for by careful system calibration or the *in situ* point spread function retrieval (INSPR)[34]. Using a straight line as an example, if we localize it with the mismatched PSF, the reconstruction will be biased toward structural distortion. This model uncertainty is visible as seen in **Supplementary Fig. 29** (structural distortion, the original straight line to a tilted line), which could not be detected by the our rFRC.

In learning-based applications, as a purely data-driven approach that learns representations of training data, the model uncertainty and data uncertainty will not be mutually exclusive[15]. Our framework has shown the possibility of detecting both model (part of leaked model uncertainty) and data uncertainties, as demonstrated in **Fig. 3** and **Supplementary Fig. 24**. Alternatively, we estimated the data and model uncertainty in learning-based applications by data sampling twice and network training twice, respectively. By applying the rFRC map to the twin predictions from two inputs (obtained from data sampling twice) and two models (obtained from network training twice), we can effectively obtain both the data and model uncertainties.

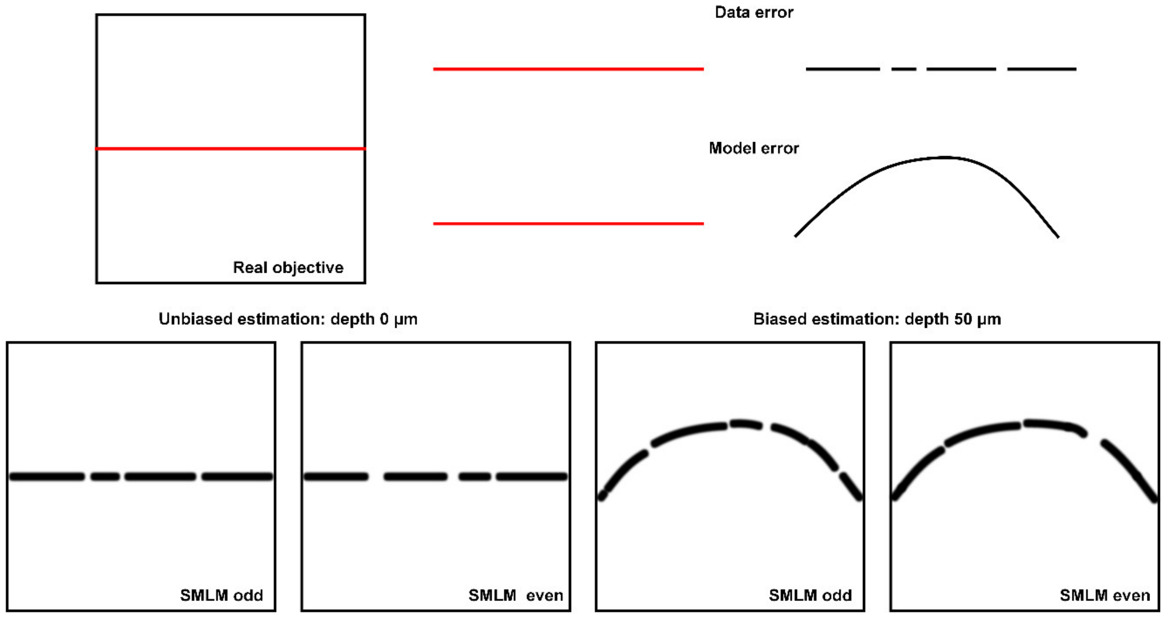


**Supplementary Fig. 29 | Unbiased and biased estimations of SMLM imaging.** The straight-line SMLM example enables visualizing unbiased and biased estimations. The real object is a straight line, and the PSF model is approximated as a Gaussian function. At a depth of 0 μm, such PSF model approximation is close enough to the real world (unbiased estimation) model reconstructing the straight-line structure. However, at a depth of 50 μm, due to the index mismatch and sample-induced aberrations, the real-world PSF model will deviate from the point-source Gaussian function. Therefore, the reconstruction (still using the Gaussian function) will distort the straight-line structure to a tilted line (biased estimation). In this case, such model uncertainties cannot be directly detected by the rFRC map.

#### Supplementary Note 8.2 | Limitations of the two-frame rFRC map.

**Limitation 1**: The normal rFRC method requires two statistically independent images with exact details. The single-frame rFRC procedure can moderate these requirements.

**Limitation 2**: If the errors are fixed patterns induced by the biased reconstruction model, they will be ignored in the rFRC map. For example, the rFRC using two independent captures cannot reveal errors due to the absence of the identical component in two measurements simultaneously. This limit is complemented by the RSM method to some extent. Besides, in learning-based applications, the model uncertainty can be uncovered by mapping two predictions from the two independently trained models.

**Limitation 3:** The rFRC can assess local qualities up to (≥) the corresponding SR scale. If the errors are smaller than the SR scale, such as the snowflake-like artifacts, they will be ignored in the rFRC map.

#### Supplementary Note 8.3 | Limitations of the single-frame rFRC.

##### For optical imaging.

**Limitation 1:** The diagonal of the pixel in the corresponding image should satisfy the Nyquist sampling criterion, in which it requires a pixel size that is $\sqrt{2}$ times smaller, i.e., $pixel < resolution / 2\sqrt{2}$. Compared to the two-frame rFRC, the single-frame approach is usually unstable.

**Limitations 2 and 3:** Similar to those of the two-frame rFRC.

**For deep-learning approaches.**

**Limitation 1:** Special care should be taken when determining the proper additive noise amplitude for the input image. An irrelevant small or large magnitude may lead to false positives or negatives. Appropriate noise magnitude should be chosen according to their specific situation.

**Limitations 2 and 3:** The same as for the two-frame rFRC.

**Supplementary Note 8.4 | Limitations of the modified RSM.**

**Limitation 1**: The RSM converts the SR image to its low-resolution scale; thus, it can detect only low-resolution errors. In contrast, errors at the SR scale (small-magnitude error components) estimated by the RSM may be false negatives, and using the rFRC map and segmentation may reduce the problem.

**Limitation 2**: The RSM map is the absolute residual image between the $I_L$ and $I_{HS}$. This map is highly corrupted by the intensity and illumination, leading to incorrect quantifications. This issue can be relieved by image segmentation.

**Limitation 3:** The RSM requires a high-SNR wide-field low-resolution image as a reference.

**Limitation 4**: The spatially invariant 2D Gaussian kernel convolution assumption may not apply to any optical system, not only introducing false negatives, but also limiting its application to 3D or non-Gaussian convolution data (e.g., denoising applications). Thresholding and segmentation can be used to mitigate this limitation.

In summary, we used a hard threshold of 0.5 in the RSM to detect only significant errors. The removal of minor errors reduces the number of potential false negatives posed by **Limitations 1**, **2**, and **4**. In addition, we apply the complemented rFRC map, which compensates for **Limitation 1** of RSM.

### Supplementary Tables.

**Supplementary Table 1 | Parameters of geometrical structure simulations.**

For a rectangle, $dimension_1$ is the height, and $dimension_2$ is the width. For a square, $dimension_1$ is the length of each side. For a triangle, $dimension_1$ is the base, and $dimension_2$ is the height. For a circle, $dimension_1$ is the radius.

| Geometry | $Dimension_1$ | $Dimension_2$ | Number |
|---|---|---|---|
| Rectangle | 5~40 pixel | 5~40 pixel | 3~5 |
| Square | 20~50 pixel | \ | 3~5 |
| Triangle | 35~50 pixel | 50~100 pixel | 3~5 |
| Circle | 10~50 pixel | \ | 3~6 |

**Supplementary Table 2 | Details on network architecture, loss function, and training procedure.** PSF SR: the 240 nm PSF to 120 nm PSF image transformation.

| **Parameters** | **Sparse sampling (Supplementary Fig. 24)** | **Noise2Noise (Fig. 4j-4m)** | **PSF super-resolution (Fig. 3)** |
|---|---|---|---|
| Training patch size | 256 × 256 | 256 × 256 | 128 × 128 |
| No. of epochs | 300 | 100 | 100 |
| Batch size | 8 | 32 | 32 |
| No. of images | 5000 | 16000 | 25 |
| Learning rate | $10^{-4}$ | $10^{-4}$ | $10^{-4}$ |
| Topology | U-net[1] | U-net[1] | U-net[1] |
| Parameters | $4.18 \times 10^{7}$ | $4.18 \times 10^{7}$ | $4.18 \times 10^{7}$ |
| Loss function | MAE | MAE | MAE |
| Optimizer | Adam | Adam | Adam |

### References.


1. Heel, M.v. & Schatz, M. Fourier shell correlation threshold criteria. *Journal of Structural Biology* **151**, 250-262 (2005).
2. Nieuwenhuizen, R.P. et al. Measuring image resolution in optical nanoscopy. *Nature Methods* **10**, 557-562 (2013).
3. Culley, S. et al. Quantitative mapping and minimization of super-resolution optical imaging artifacts. *Nature Methods* **15**, 263-266 (2018).
4. Tortarolo, G., Castello, M., Diaspro, A., Koho, S. & Vicidomini, G. Evaluating image resolution in stimulated emission depletion microscopy. *Optica* **5**, 32-35 (2018).
5. Koho, S. et al. Fourier ring correlation simplifies image restoration in fluorescence microscopy. *Nature Communications* **10**, 1-9 (2019).
6. Huang, X. et al. Fast, long-term, super-resolution imaging with Hessian structured illumination microscopy. *Nature biotechnology* **36**, 451-459 (2018).
7. Gustafsson, M.G. et al. Three-dimensional resolution doubling in wide-field fluorescence microscopy by structured illumination. *Biophysical Journal* **94**, 4957-4970 (2008).
8. Demmerle, J. et al. Strategic and practical guidelines for successful structured illumination microscopy. *Nature Protocols* **12**, 988-1010 (2017).
9. Richardson, W.H. Bayesian-based iterative method of image restoration. *Journal of The Optical Society of America A* **62**, 55-59 (1972).
10. Lucy, L.B. An iterative technique for the rectification of observed distributions. *The Astronomical Journal* **79**, 745 (1974).
11. Zheng, G., Horstmeyer, R. & Yang, C. Wide-field, high-resolution Fourier ptychographic microscopy. *Nature Photonics* **7**, 739-745 (2013).
12. Hell, S.W. & Wichmann, J. Breaking the diffraction resolution limit by stimulated emission: stimulated-emission-depletion fluorescence microscopy. *Optics Letters* **19**, 780-782 (1994).
13. Vicidomini, G. et al. Sharper low-power STED nanoscopy by time gating. *Nature Methods* **8**, 571-573 (2011).
14. Wang, Z., Bovik, A.C., Sheikh, H.R. & Simoncelli, E.P. Image quality assessment: from error visibility to structural similarity. *IEEE Transactions on Image Processing* **13**, 600-612 (2004).
15. Kendall, A. & Gal, Y. What Uncertainties Do We Need in Bayesian Deep Learning for Computer Vision? *Advances in Neural Information Processing Systems*, 5580-5590 (2017).
16. Moeckl, L., Roy, A.R. & Moerner, W. Deep learning in single-molecule microscopy: fundamentals, caveats, and recent developments. *Biomedical Optics Express* **11**, 1633-1661 (2020).
17. Goan, E. & Fookes, C. Bayesian Neural Networks: An Introduction and Survey. *Case Studies in Applied Bayesian Data Science* **2259**, 45-87 (2020).
18. Gal, Y. & Ghahramani, Z. Dropout as a Bayesian Approximation: Representing Model Uncertainty in Deep Learning. *Proceedings of Machine Learning Research*, 1050-1059 (2016).
19. Hinton, G.E. & Van Camp, D. Keeping the neural networks simple by minimizing the description length of the weights. *Proceedings of the sixth annual conference on Computational learning theory*, 5-13 (1993).
20. Lakshminarayanan, B., Pritzel, A. & Blundell, C. Simple and Scalable Predictive Uncertainty Estimation using Deep Ensembles. *Advances in neural information processing systems* **30** (2017).
21. Betzig, E. et al. Imaging Intracellular Fluorescent Proteins at Nanometer Resolution. *Science* **313**, 1642-1645 (2006).
22. Rust, M., Bates, M. & Zhuang, X. Sub-diffraction-limit imaging by stochastic optical reconstruction microscopy (STORM). *Nature Methods* **3**, 793-796 (2006).
23. Gustafsson, N. et al. Fast live-cell conventional fluorophore nanoscopy with ImageJ through super-resolution radial fluctuations. *Nature Communications* **7**, 1-9 (2016).
24. Dertinger, T., Colyer, R., Iyer, G., Weiss, S. & Enderlein, J. Fast, background-free, 3D super-resolution optical

fluctuation imaging (SOFI). *Proceedings of the National Academy of Sciences of the United States of America* **106**, 22287-22292 (2009).
